# *De Novo* Non-Canonical Nanopore Basecalling Enables Private Communication using Heavily-modified DNA Data at Single-Molecule Level

**DOI:** 10.1101/2024.08.15.606762

**Authors:** Qingyuan Fan, Xuyang Zhao, Junyao Li, Ronghui Liu, Ming Liu, Qishun Feng, Yanping Long, Yang Fu, Jixian Zhai, Qing Pan, Yi Li

**Affiliations:** School of Microelectronics, MOE Engineering Research Center of Integrated Circuits for Next Generation Communications, Southern University of Science and Technology, 518055, Shenzhen, China; School of Medicine, Southern University of Science and Technology, 518055, Shenzhen, China; National Clinical Research Center for Infectious Diseases, Shenzhen Third People’s Hospital, The Second Affiliated Hospital of Southern University of Science and Technology, Shenzhen, 518112, China; Department of Biology, School of Life Sciences, Southern University of Science and Technology, 518055, Shenzhen, China; College of Information Engineering, Zhejiang University of Technology, 310023, Hangzhou, China

**Author notes:** (Y.L.).

## Abstract

Hidden messages in DNA molecules by employing chemical modifications has been suggested for private data storage and transmission at high information density. However, rapidly decoding these “molecular keys” with corresponding basecallers remains challenging. We present DeepSME, a nanopore sequencing and deep-learning based framework towards single-molecule encryption, demonstrated by using 5-hydroxymethylcytosine (5hmC) substitution for individual nucleotide recognition rather than sequential interactions. This non-natural, motif-insensitive methylation disrupts ion current, resulting in a readout failure of 67.2%-100%, concealing the privacy within the DNAs. We further develop an alignment-free DeepSME basecaller as a key to reconstitute the digital information. Our three-stage training pipeline, expands k-mer size from 4^6^ to 4^9^, achieving over 92% precision and recall from scratch. DeepSME deciphers fully 5hmC concealed text and image within 16× coverage depth with an F1-score of 86.4%, surpassing all the state-of-the-art basecallers. Demonstrated on edge computing devices, DeepSME holds supreme potential for DNA-based private communications and broader bioengineering and medical applications.

## Introduction

DNA storage has emerged as a promising solution for the requirement of digital memories in the ‘zettabyte era’ due to its density and durability^1^. Design and realization of molecular protection^2^ places ever-increasing demands on the accurate storage, transmission and assessment of DNA data rapidly and privately. Many efforts have been devoted to exploring an accessible approach for information privacy, such as cryptoristors^3^, metahologram^4,5^, quantum photonic system^6,7^, DNA based technology^8–11^, synthetic macromolecules^12,13^, where concealed information would not be evident to unsuspecting persons’ examination^10^. Among them, biomolecular steganography, which utilizes chemical modifications or interactions instead of computational schemes, has been demonstrated using nucleic acids, proteins^14^, aptamers^15^, and bacteria^16^ for information concealment. For instance, DNA-based steganography developed by Clelland et al.^17^ was further leveraged by DNA origami cryptography, which creates a key with a size of over 700 bits with oligonucleotides^11^. However, these strategies exploiting molecular interactions with low information densities and time-consuming processing, have not fully explored the potential of chemical modifications to densely packed nucleobases, thus preventing them from being used in sequencing-based approaches.

Sequencing non-canonical DNA holds promise for private communication because current sequencing technologies and basecallers are designed for canonical DNA. When encountering modified nucleobases, these state-of-the-art sequencing technologies introduce various errors, making it challenging to retrieve the correct sequence^18^. Such methylated^19^ or oxidized^20^ modifications necessitate specialized methods across different sequencing platforms, such as next generation sequencing (NGS)^21^ or long-read ones. For NGS techniques, WGBS^22,23^, RRBS^24^, MeDIP-seq^25^, MRE-seq^26^ etc. have been developed for single specific types, where the non-canonical bases may not react completely or break the DNA strands into pieces^27^. These methods can also be applied onto the long-read fluorescent sequencing (PacBio), where the drawbacks are the same. Meanwhile, for long-read nanopore sequencing, non-canonical bases can be identified without any changes. The detection algorithms can be categorized into two classes: 1) Identification from alphabet errors. This includes Epinano for m6A^28^, differ for m6A^29^, DRUMMER for m6A^30^, nanoRMS for Ψ U/Nm^31^, ELIGOS for m6A^32^, NanoCEM for unspecific modifications^33^ and Dinopore for Inosine^34^; 2) Identification from current features, containing DeepSignal for 5mC^35^, DeepMod for 5mC^36^, MINES for m6A^37^, nanoDoc for unspecific modifications^38^, nanom6A for m6A^39^, Nanocompore for m6A^40^, Yanocomp for m6A^41^, xPore for m6A^42^, Penguin for Ψ ^43^ m6Anet for m6A^44^ and exos for 8-oxo-dG^20^; Rerio for 5mC/5hmC and IL-AD for 5mC/5hmC/6mA^45^. Aiming for biological science, these developments of basecallers limit themselves within the identification of light-weighted modification^46^ or motifs^47^. None of them - to the best of our knowledge – has been reported to deal with heavily or nearly complete modifications, which remains challenging because of the lack of basecallers and references.

To address this limitation, a *de novo* approach to basecalling heavily modified DNA is required for pristine raw data. Nanopore data readouts, maintaining sequential information of any nucleobase or analogues into ion flow blockades, nowadays utilize neural networks (NN)^48^ vastly for accurate basecalling. Well-trained NNs can act as the cipher to pair with a specific molecular key, the inherent non-reversibility of neural networks prevents direct determination of the key from the network’s weights. However, training NNs is often a joy with tears, since rounds of sophisticated alignments and time-consuming corrections of datasets may take place^49^. Besides, existing prior knowledge (k-mer dictionary) and references are critical to reduce the high error rates^50^. The bottlenecks for nanopore private communication have two aspects: 1) to arrange private messages wisely into non-canonical DNAs and 2) to rapidly and correctly build up correlated nanopore NN with limited correct alignments. To date, constructing a framework for heavily or completely modified DNAs, is therefore highly appreciated.

In this work, we propose a private framework utilizing motif-insensitive 5hmC modification combined with a nanopore sequencing-based *de novo* basecaller for privacy. We conceal text and image into fully 5hmC modified DNA sequences and validate our approach by nanopore sequencing and basecaller training, aiming to address the challenges to build the basecaller posed to heavily modified bases. We develop a three-stage alignment-free training framework for <u>Deep</u>-learning framework towards <u>S</u>ingle-<u>M</u>olecule <u>E</u>ncryption (DeepSME), which tackles the basecalling bottleneck of the heavily modified dataset by expanding k-mer dictionary.

Independent k-mer tables are generated from scratch, allowing us to process the modified sequences and their corresponded signal disruptions without prior references at single-molecule level. This framework, together with the training method of private DeepSME basecaller underpins the potential for concealed DNA-based data storage and communication with high information density, addressing the increasing demand for robust information privacy in an era of evolving biotechnological threats.

## Results

Fig. 1a illustrates the scenario for digital information stored in DNA for private communications. Texts and Images can be transcoded into DNA sequences, followed by the DNA synthesis as templates. Conventional polymerase chain reactions (PCR) amplify the templates into double strands and can be sequenced by either next generation sequencing (NGS)^51^ or nanopore sequencing (NPS)^52^ for public communications. Here, Alice acts as the sender who initiates the communication either publicly or privately. For the former case, canonical bases and conventional basecallers are publicly available. Here we proposed a <u>Deep</u>-learning framework towards <u>S</u>ingle-<u>M</u>olecule <u>E</u>ncryptions (DeepSME) for the latter case between Alice and Bob. This allows Alice to hide information within non-canonical DNAs and creates a basecaller that serves as the key for Bob. Moreover, unsuspecting person Eve is not capable of accessing the information using conventional basecallers.

**Fig. 1.**
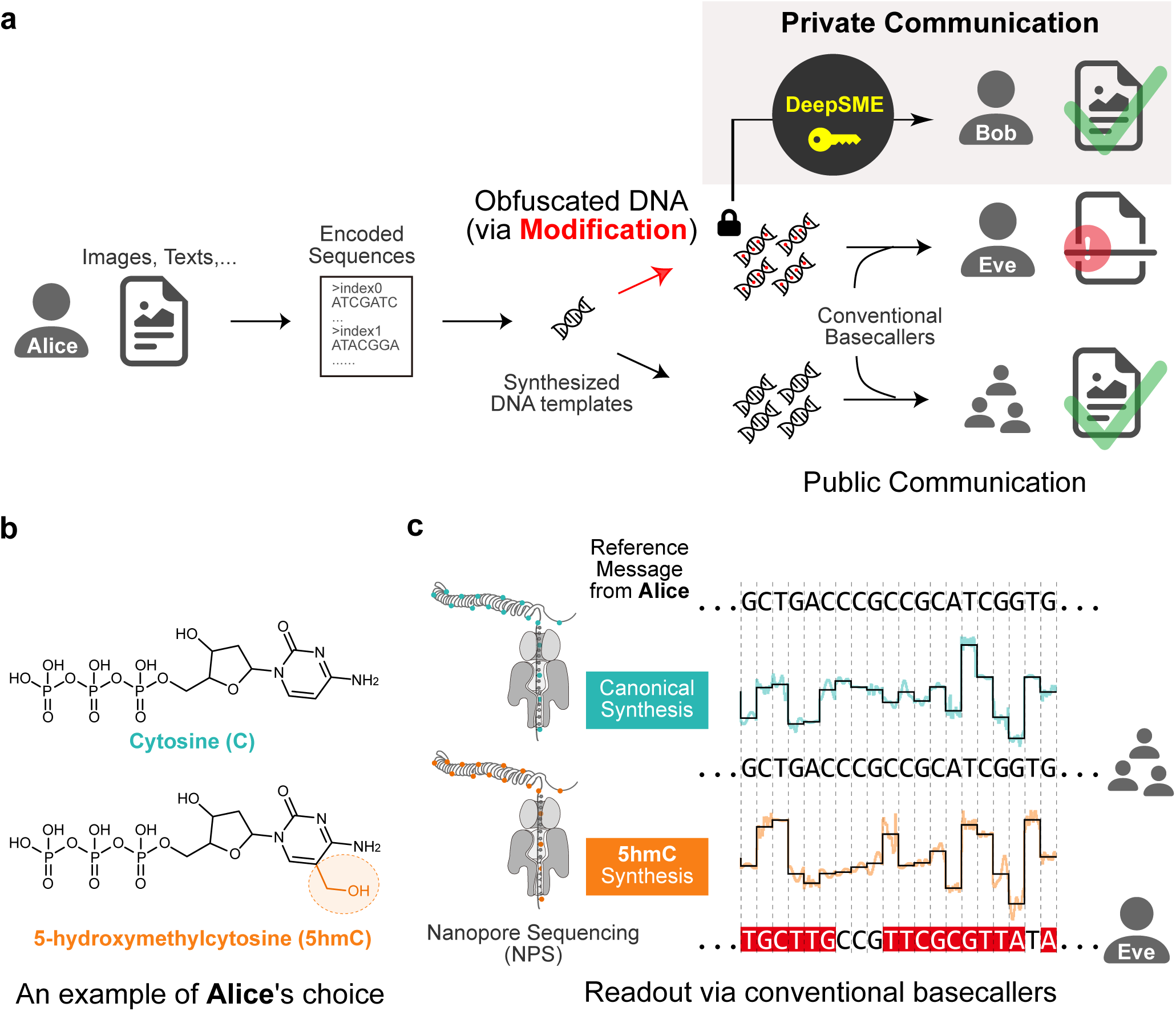
Obfuscating digital information stored in DNA for private communication via nanopore sequencing. (**a**) Schematics of private communication for files and texts stored in DNA. Steganography is achieved by obfuscating data stored inside DNA via modifications by the sender Alice. Canonical DNAs can be transmitted publicly and readout using conventional basecaller by everyone. Non-canonical DNAs in principle cannot be easily revealed by an unsuspecting observer Eve, while the private observer Bob should be able to use DeepSME as the key to decipher the messages from Alice. (**b**) An example of Alice’s choice with the chemical structure of deoxycytidine triphosphates (dCTP), which is labelled as cytosine (C) and 5-hydroxymethylcytosine (5hmC), respectively. (**c**) Nanopore sequencing of canonical and fully methylated 5hmC DNA strands using a state-of-the-art basecaller (Bonito/Dorado). The ionic current signals of C (cyan) and 5hmC (orange) are interpreted into DNA sequences and aligned with reference sequences/messages from Alice. The error bases are labelled in red, corresponding to the information received by Eve using the conventional basecaller.

Following the roles for Alice, Bob and Eve, Fig. 1b displays an example of chemical structures of Alice’s choice - deoxycytidine triphosphate (dCTP), where the two analogies are labelled as cytosine (C) and 5-hydroxymethylcytosine (5hmC). Due to an additional hydroxymethyl group at 5’, a distinct steric hinderance on the blockage of nanopore signals could be expected with close-to-neutral hydrophilicity. Following synthetic methods^36^, all the cytosines are substituted *in vitro* by the chemically stable 5hmCs regardless of any motifs or contexts. This complete replacement of 5hmC surpasses the medication ratio in any biological systems, in particular human genome^23,53,54^, where only 1.28% of C (2.62‰ of a given whole microbial genome) is currently modified to 5mC or 5hmC.

We next examined the privacy of full 5hmC modification using nanopore sequencing in Fig. 1c. As DNA molecules pass through the nanopore, they generate characteristic ionic current signals that are recorded as squiggle patterns (shown as black lines in the figure). For canonical DNA (top panel), the current signal (shown in cyan) is correctly interpreted by conventional basecallers to reveal the Alice’s reference sequence (GCTGACCCGCCGCATCGGTG). However, current signals in the bottom panel drastically deviated (shown in orange) when containing full 5hmC modifications. These heavily modified signals confuse conventional basecallers, for example Bonito/Dorado, resulting in severe misinterpretation of the message (TGCTTG<u>CCG</u>TTCGCGTTA<u>T</u>A, with 16 out of 20 bases incorrectly called), demonstrating the successful privacy protection away from Eve in this scenario.

For the corresponding key produced by either Alice or Bob, we propose a three-stage alignment-free pipeline for DeepSME basecaller, shown in Fig. 2a. Eleven samples of DNA sequences ranging from 1145 nt to 1341 nt are firstly sequenced to construct the 6-mer quality check dataset without any sequence-to-current alignments^50^. Tolerating these errors caused by the alignment-free segmentation, an initial 6-mer QC DeepSME is trained to provide a k-mer model as a new dictionary. Benefiting from investigated k-mer dictionary and nanopore simulators such as scrappie squiggle^55^, squigulator^56^, and deepsimulator^57^, *in silico* dataset can be generated for length variation and expanded the dictionary from 4^6^ to 4^9^. Not only mitigating the overall errors, simulated current signals can also expand the physical redundancy (also named as coverage depth, denoting the depth or completeness of DNA sequencing on the reference sequence). Depicted in Fig. 2b, the upper heatmap of full 6-mers shows a sparse occupation (2.60%) in the 9-mer space, while *in silico* dataset can cover almost all the 9-mers, comparable to the experimental microbial whole-genome sequences (gDNA) dataset with slightly different frequency (see Supplementary Table 5). Last, three types of microbial gDNA are sequenced by nanopore and processed by enhanced DeepSME, which leads to the final training of the reinforced DeepSME.

**Fig. 2.**
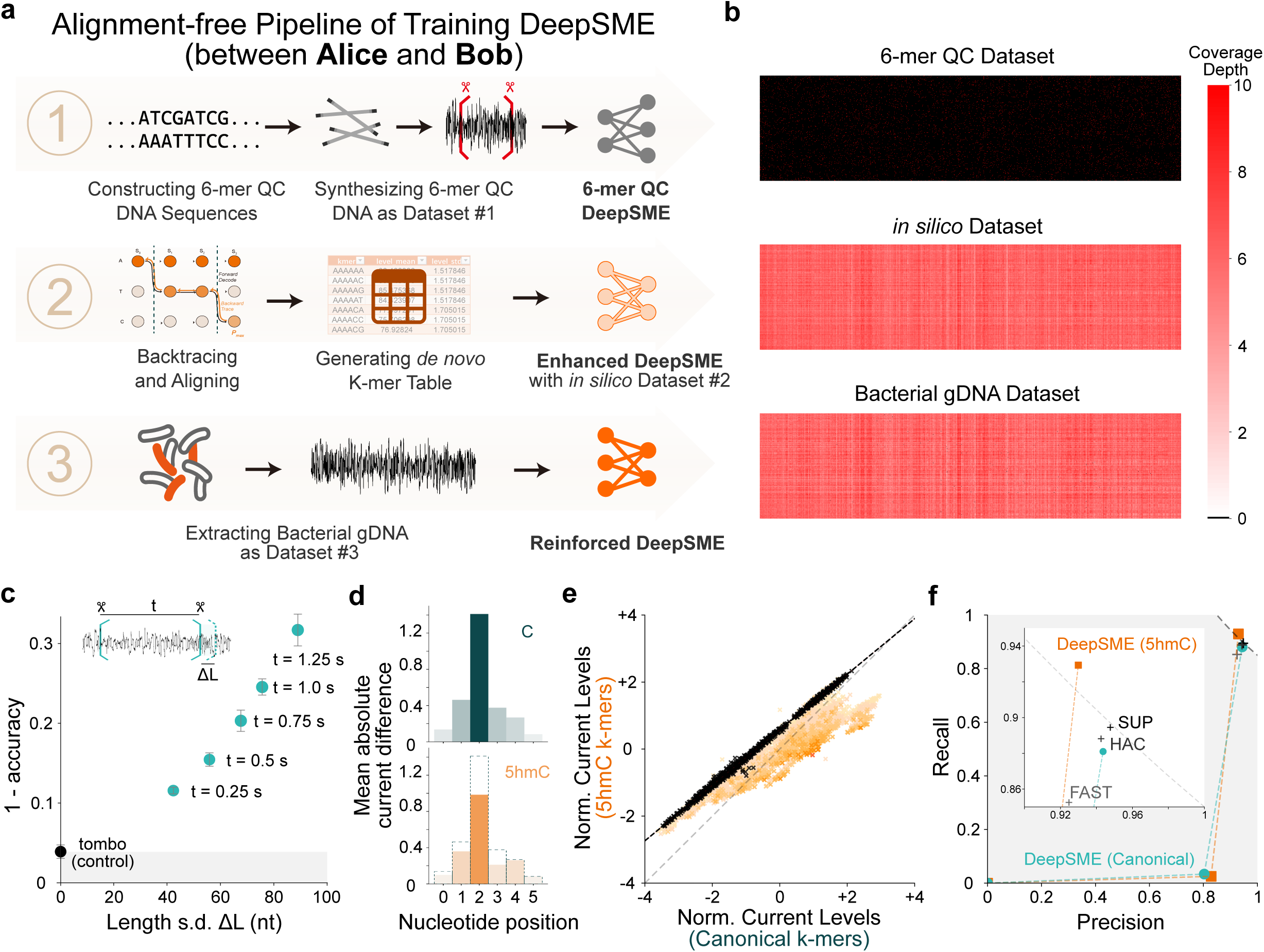
*De novo* construction of non-canonical DeepSME basecaller between Alice and Bob. (**a**) Training DeepSME with a three-stage alignment-free pipeline. (**b**) 9-mer coverages of three used training datasets. (**c**) Training error rate as a function of standard deviation of segmented lengths. Inset depicts the uncertainty (standard deviation of a certain segmented length) (**d**) Mean absolute current difference (MACD) of both canonical cytosine (top) and 5hmC (bottom) at different nucleotide positions of a k-mer oligonucleotide. (**e**) Methylated 5hmC vs. canonical cytosine of normalized current levels. (**f**) Recall vs. precision of DeepSME for both canonical and methylated 5hmC.

In the first stage, the 6-mer quality check (QC) DNA is sensitive to initiate the DeepSME. It is hardly practical to mark 500 annotations and segmentations for every 10,000 heavily modified nucleobases manually with no prior knowledge. Taking conventional segmentation tool Tombo^50^ (Nanopolish^58^ also works) on canonical dataset as a control, 3.9% errors will always exist. Considering billions of sequences and bases that are being used, Fig. 2c shows the overall error rate as a function of length variation. Given approximately 450 nt/s as the sequencing speed of a nanopore^59^, individual current signals segmented as 2.22 s (corresponding to roughly 1000 nt), 4.44 s (∼ 2000 nt), 6.66 s (∼ 3000 nt), 8.88 s (∼ 4000 nt) and 11.10 s (∼ 5000 nt) contribute to the length variations of 40 nt to 85 nt. The longer the segmentation goes, the more errors the preliminary DeepSME holds.

The inset of Fig. 2c illustrates the segmentation-induced length variation. A fixed number of bases can be assigned to segmented current blocks each time as a chunk, creating a dataset containing tens of thousands of chunks for the training of the basecaller. The length variation itself is proportional to the segmented time. Our preliminary DeepSME segmented chunks at 2.22 s is estimated to have a total error of 11.6%, which seems tolerable for further investigations.

We further analyzed the reduced ionic currents with the modification of 5hmC. Fig. 2d shows the normalized current difference for the canonical C and 5hmC in the extracted k-mer model from DeepSME in the second training stage. By substituting either C or 5hmC at each position (x-axis), the mean averaged current difference (MACD) is calculated over all k-mers in the model. In this 6-mer model, the base at index 2 has the highest impact on the current change. This is in a reasonable agreement with reported k-mer model from ONT^60^. Additionally, the weight of the bases at index #0, #1, #2, and #3 on the current significantly decreases when 5hmC modification takes place. This indicates that 5hmC modification leads to a reduction in the normalized currents at these corresponding base positions compared to the canonical state.

On the other hand, Fig. 2e presents a comparative analysis of normalized ionic current levels between canonical and 5hmC-modified DNA sequences. Each point in the figure represents a specific k-mer (6-mer), where those not containing any C’s shown in black, and those containing 1 to 6 C’s shown in orange. The intensity of the orange color correlates with the number of cytosines in the k-mer - darker orange indicates more cytosines. The black dashed line with a slope of 0.83 with the offset of 0.58 represents a linear fitting for 6-mers without C’s. Notably, this line deviates from the gray diagonal one, aligning with the k-mer for 5hmC modified DNA reported by Kovaka et al^60^. This significant deviation in slope indicates substantial changes in the normalized current signals, making the non-natural heavily modification more effective for private communications. Furthermore, k-mers containing 5hmC modifications consistently show lower normalized current levels compared to their canonical counterparts, which matches well with our previous observations in Fig. 2d.

The basecalling accuracy and recall are evaluated at the third stage. Fig. 2f illustrates the relationship between recall and precision of DeepSME at three stages for both canonical C and 5hmC. First, the preliminary basecallers are located at the bottom left, meaning low precision and low recall capabilities since only 6-mer features of DNA are learnt so far. Second, the enhanced basecallers perform much better on precision but remain low on the recall, which is reasonable since the simulated dataset lacks the ability of generalization for variant conditions. Last, the reinforced basecallers show significant improvements in both precision (0%-82.9%-93.0%) and recall (0%-2.5%-92.9%). The reinforced 5hmC basecaller has a slightly lower accuracy but a higher recall than that of canonical one. Compared with the commercial basecallers (Bonito FAST, HAC - high accuracy and SUP - super accuracy config), the reinforced 5hmC basecaller is beyond the F1-score of 0.85 (grey dashed line), surpassing Bonito SUP, HAC as well as FAST.

We turn our focus onto the decoding results of 5hmC modified DNA that conceals text file (sustech_introduction.txt, 978 bytes, 55 strands) and image file (sustech_logo.jpg, 7,775 bytes, 432 strands). Without knowing exact types of DNA modifications, Fig. 3a shows the decoding performance of the identical text file using state-of-the-art basecallers on modified DNA from the unsuspecting observer Eve’s view with 16× coverage. For the basecallers that could not reach the coverage amount, all basecalled sequences were obliged to use. Guppy 6.0 achieved a decode rate of 5.45% (3/55), Bonito/Dorado reached 3.64% (2/55), Rerio 5mC 5hmC managed 3.64% (2/55), while IL-AD was unable to decode any sequences (0/55) at all.

**Fig. 3.**
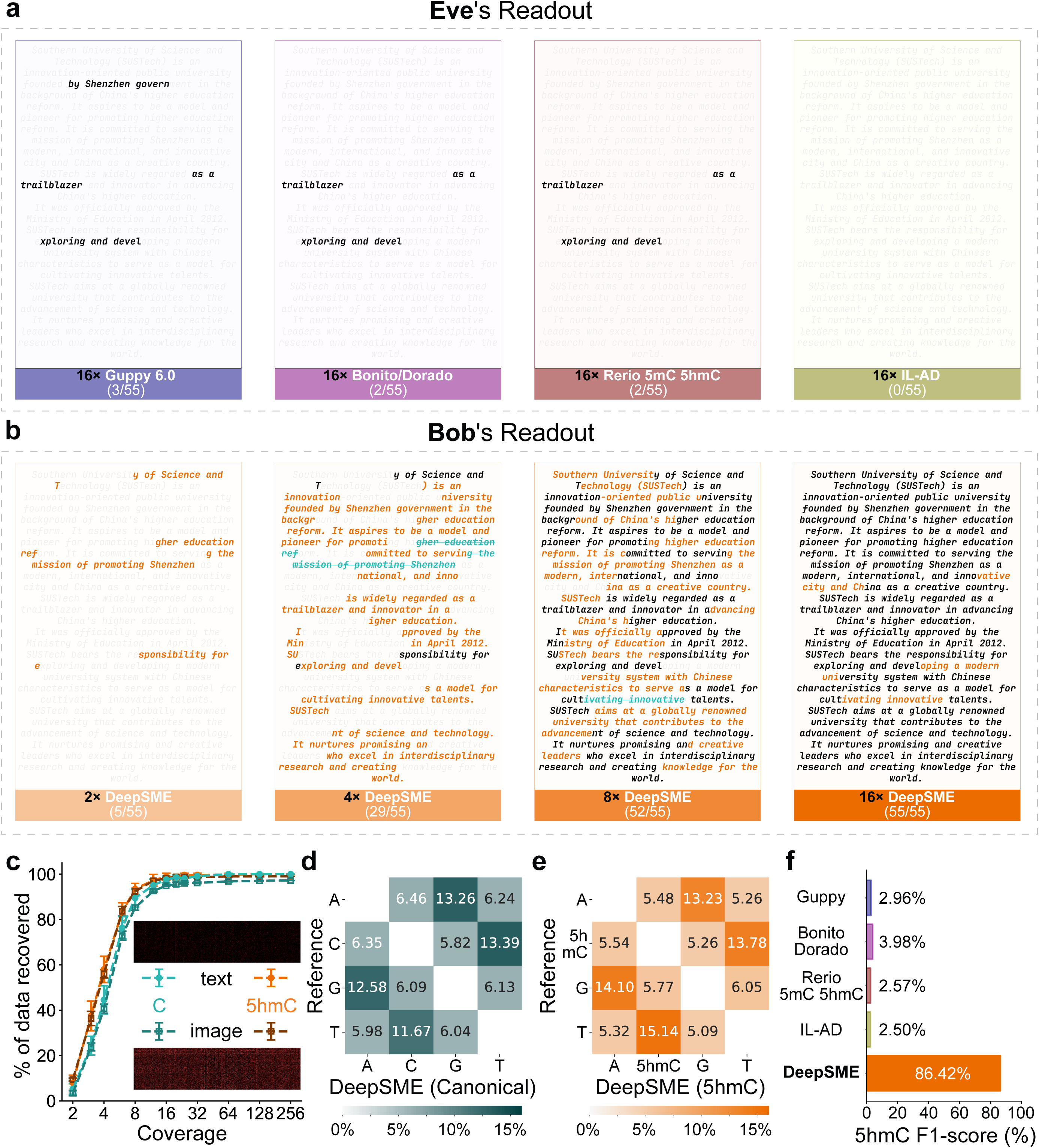
Deobfuscation performance of digital information enabled by DeepSME. (**a**) Decoding performance of the text file by the unsuspecting observer Eve using Guppy 6.0 (`dna_r9.4.1_450bps_hac` config), Bonito/Dorado (dna_r9.4.1_e8_sup@v3.3 config), Rerio 5mC 5hmC (`res_dna_r941_min_modbases_5mC_5hmC_v001` config), Incremental Learning and Anomaly Detection (IL-AD, 5hmC config) with coverage depths of 16×. (**b**) Decoding performance of the text file by the private observer Bob using DeepSME with coverage depths of 2×, 4×, 8× and 16×. The transcoded strand amounts are 5, 33, 52 and 55 of total 55. (**c**) The percentage of data recovered as a function of coverage for the text file stored in canonical (cyan) or 5hmC (orange) using corresponding canonical or 5hmC DeepSME, respectively (n=10). (**d**) Base bias of substitutions for canonical DeepSME. (**e**) Base bias of substitutions for 5hmC DeepSME. (**f**) F1-scores for variant basecallers on obfuscated (5hmC) DNA.

Encouraged by the high precision and recall of DeepSME, we perform decoding experiments displayed in Fig. 3b from the private observer Bob’s view. The recovered texts are shown with the increasing physical redundancy (the coverage depth from 2× to 16×). It is evident that DeepSME produced up to 52.73% (29/55) of the text after decoding with 4×, reached up to 94.5% (52/55) data retrieval with 8× coverage and correctly decoded all of the text with 16× coverage. Fig. 3c compares the text data retrieval performance of the two DeepSMEs as a function of coverage depth. They exhibit almost identical data recover rate, with small variations across experiments. At each coverage level, the data recover rate was measured 10 times with different random seeds.

To gain more insights from the two basecallers, the substitution probabilities of canonical and 5hmC DeepSME are illustrated in Fig. 3d and 3e. The substitutions from T to C and C to T are 11.67% and 13.39%, while these values go to 15.14% and 13.78% for T-5hmC and 5hmC-T. A-G and G-A are also high for both. The two matrixes are quite comparable, which are also in agreement with the matrix from Lopez et al^52^.

To analyze the comparison with alternative basecaller configs, Fig. 3f presents the F1-scores on our modified DNA storage dataset. DeepSME achieved a leading F1-score of 0.864 to date, while other basecallers such as Guppy 6.0, Bonito/Dorado Super, Rerio, and IL-AD had F1-scores ranging only from 0.025 to 0.040.

## Discussion

Our work demonstrates that the challenge of pairing key generation with nanopore basecallers can be coped with the DeepSME framework. This could enable private communication between Alice and Bob with a superior F1-score of 86.42%, shown in Fig. 3f Our DeepSME framework offers several advantages:

1. *Knowledge growing*. First, the initial 6-mer QC DeepSME basecaller does extract 5hmC-modified features. This is supported by the shift of k-mer current in Figure 2e, where 89.63% of C-containing k-mers deviate from the diagonal and only 12.35% of k-mers that don’t contain C deviate from the diagonal. Second, the Enhanced DeepSME basecaller could learn more abundant features from 6-mers to 9-mers, which can be ascribed to the substantial precision increase on the modified test dataset from 0% to 82.87% (see Figure 2f and Supplementary Table 7). Third, the reinforced DeepSME basecaller could get polished smoothly along with experimental 9-mer gDNA dataset. This can be accredited by the fact of achieving 92.99% precision and 92.93% recall.
2. *Alignment free.* We used the strategy of fixing chunk size of 1000 bp instead, shown in Fig. 2c. Based on the assumption of constant sequencing velocity at 450 bp, this strategy holds an error rate as high as 11.6%. Compared with 5.63% error as the minimal, this high error rate is no more a trouble as the overall basecalling errors went down to 7.01% in Fig. 2f.
3. *Prior-knowledge free*. Our training of fully modified dataset can be performed without pre-existing k-mer tables nor basecaller weights. This is underpinned by a smart use of the output from the Connectionist Temporal Classification (CTC) layer of our 6-mer QC DeepSME basecaller. Once this basecaller gets trained, one can directly derive a new k-mer table from the CTC layer, which is particularly beneficial to link with the trained weights. To this end, prior k-mer tables and basecaller weights can be detoured.
4. *Affordable*. We show in the Methods that 266.4 minutes of thee-stage basecaller training in 19.9 GB VRAM are sufficient for our open-sourced deep-learning neural network. While lightweight models can reduce computational demands, they led to much lower F1-scores (75.77%-84.36%). Different programming language-based software (Python-based Bonito and C-based Dorado) and hardware platform (NVIDIA RTX 3090 and Jetson Xavier AGX) were thoroughly evaluated (see Supplementary Table 11 and 12), confirming the operational feasibility of the current model size in edge-computing environments.

For the generalizability, DeepSME basecallers could be custom tailored and delivered for variant combinations on promise, including diverse modification types, modification ratios as well as nanopore sensors. For instance, our preliminary tests demonstrate successful alignment of the sequence primer current extracted from the 6-mer QC sequence (using an R9.4.1 pore) to its corresponding current in an R10.4.1 pore via Dynamic Time Warping^61^ (DTW) (Supplementary Information Figure 6). Although high error rates have been mitigated by Composite Hedges Nanopore (CHN) code, the advances on R10.4.1 flowcells could further manage the error rate below 1%^62^. This cross-pore compatibility suggests the adaptability of the DeepSME framework across different types of nanopores or other hardware such as CycloneSEQ^63^ or PolyseqOne^64^.

Our three-stage training pipeline is essential, as removing any stage would compromise the overall performance and reliability of the framework. First, the initial 6-mer QC DeepSME basecaller lays the foundation of generating a unique 6-mer table for subsequent steps. Without this initial step, fewer than 0.2% of the chunks can be identified for the rest steps, making basecaller training nearly impossible. Second, the *in silico* training of the Enhanced DeepSME basecaller boosts the accuracy to over 80% while expanding the k-mer coverage from 6-mers to 9-mers. Without 81,490 simulated strands, the 6-mer QC DeepSME basecaller would recognize less than 0.5% of chunks of gDNA dataset, hindering basecalling. However, the Enhanced DeepSME basecaller has the recall rate remaining at 2.5%, showing its overfitting of *in silico* data. Last, the reinforced DeepSME basecaller incorporating real-world gDNA data, reached 93.0% precision on DNA storage dataset (Fig. 3). This demonstrates the successful mitigation of overfitting risks.

For the unsuspecting observer Eve, it is hardly possible to circumvent DeepSME with public basecallers and their weights. Alternative basecallers have fewer than 5% of the correct sequences for modified sequences in Fig. 3f (see the Supplementary Table 7-11 for recall and precision of these basecallers). We believe this could be attributed to the shifted current deviations (Fig. 1c and Fig. 2e): some of the points are still located close to the black dashed line, which could be recognized by basecaller and aligned to the reference sequence. Meanwhile, other points are far away from the black dashed line, being ill-interpreted.

Beyond enumerating basecalling attempts, the interception of DeepSME could be executed by Eve as the weights lack inherent encryption. Thus, employing traditional^65^ or post-quantum cryptographic algorithms^66^ becomes essential and be a critical direction to avoid cybersecurity threats between Alice and Bob. Furthermore, reverse engineering of modification patterns represents another potential vulnerability. However, the diversity of existing modifications^19,67,68^ and the availability datasets for heavily modified DNA, may hinder the timely training of such reverse-engineering models, providing DeepSME with a valuable, albeit potentially temporary, window for privacy, though not absolute security.

Lastly, privacy may be inferred, assuming an ultimate modification caller that can accurately decode any level of modified data. While theoretically plausible, achieving this in practice presents considerable hurdles. One one hand, the complexity of the “molecular key space” increases with 429 existing types^19,67,68^ and upcoming new ones. On the other hand, the parameters and memory consumption will inflate drastically for training an 100-letter basecaller (may need 5.6 + 14.3 × 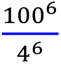 = 3.49 × 10^9^ GB of VRAM). These practical issues offers DeepSME a substantial layer of privacy against current threats and resource-constrained adversaries.

We speculate that DeepSME could evolve into a functional encryption system. The original DNA sequence could stand for the plaintext, the nanopore current readouts from modified DNA for the ciphertext, the specific modification types and their ratios for the key space, while *in vitro* modification process and DeepSME weights governed basecalling for encryption and decryption, respectively. While sharing similarities as a quintuple scheme for the definition of encryption^69^, we are at a very infant stage towards encryption. Given the key length is defined as log_2_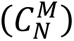, where *M* is the known nucleobase analogues and *N* is the chosen nucleobase types, our implementation with a single modification type achieves a key length of approximately log_2_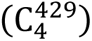 = 30.37 bits, supposing the existence of at least 429 known nucleobase modifications^19,67,68^. This is far behind National Institute of Standards and Technology’s (NIST) recommended minimum of 128 bits for secure encryption^65^. In this theory, 24 types of nucleobase analogues used in one *in vitro* process could fulfil the key length requirement of log_2_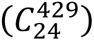 = 129.89 bits. Alternatively, block encryption offers a promising approach to increase key length. By splitting the data into blocks and employing different combinations of nucleobase analogues for each block, we can linearly increase the overall key length. For instance, using four blocks and focusing on a subset of 140 common base analogues mentioned by Juan Et. al.^19^, we can select four distinct sets of six analogues to encrypt each data block. This approach could theoretically achieve a key length of 4 × log_2_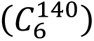 = 132.51 bits, thereby meeting NIST’s requirements. These methods, while beyond the scope of this work, highlight the potential of DeepSME for future development as a robust encryption system.

We also verified the integration of DeepSME with post-quantum cryptography (PQC) such as frameworks like FrodoKEM^71^. This could also be applied onto the raw texts and images. For tamper-evident DNA storage protocols; we foresee that techniques such as molecular tags, sequence watermarks, and enzymatic protection mechanisms could be explored^72^. Some simplied methods, like monitoring 5hmC/canonical cytosine ratios and basecaller discrepancies which reuse the developed basecallers, may also provide effective tamper-evidence. We further note that 5hmC DeepSME basecaller also performs well on canonical DNA sequences (see Supplementary Table 11), which shows its promise to be utilized as a framework to detect methylations from epigenomics.

For ethical considerations, we have implemented multi-layered biosecurity screening measures and advocate for adherence to guidelines like the International Gene Synthesis Consortium (IGSC) standards and ISO regulations^73^. Concerns regarding illicit data smuggling via microbes are taken into our thoughts. Although they could be mitigated by the likely toxicity of extensive 5hmC modifications^74^, proactive biosecurity, ethical guidelines, and dual-use risk assessments remain essential for responsible DeepSME development, as we have solely focused on *in vitro* investigations^75^.

In conclusion, we have proposed and experimentally validated a communication framework (DeepSME) with non-naturally existing hydromethylation DNAs for information privacy, paired with a *de novo* trained nanopore basecaller. Avoiding tremendous alignments and corrections, we demonstrate that our three-stage training method is essential for DeepSME to achieve both 92% recall and precision incrementally, exceeding state-of-the-art basecallers on the digital information stored in 5hmC DNA dataset. Within just 16× coverage depth and negligible prior knowledge requirements, our independent DeepSME framework fulfills the readout of heavily-modified single-molecule DNA, showcasing its great promise towards biomolecular-based private communications, unclonable functions and anticounterfeiting systems.

## Statistics & Reproducibility

Statistics used to analyze DNA storage decoding performance are described in the methods section. Analysis can be reproduced using datasets deposited in the NCBI SRA and Zenodo (see “Data availability” statement) and the code developed for this work (see “Code availability” statement). All sequencing data from 6-mer QC sequences and Bacterial Genome DNA (gDNA) were used to train DeepSME. All sequencing data from DNA storage strands were used to perform the DNA decoding experiments. The decoding rates under different coverages shown in Fig. 3c were determined by repeating the analysis 10 times using random seeds 0-9 when selecting reads from each sequence group.

## Methods

### Information Concealment via 5-hydroxymethylcytosine (5hmC) based Polymerase Chain Reaction

Synthetic DNA sequences with designed digital information were purchased from Twist Bioscience, while plasmids and primers were obtained from Tsingke Biotech. Polymerase chain reactions (PCR) were performed using 10 mM dATP, dTTP, dGTP and dCTP supplied by Sangon. The PCR process utilized Phanta Max Super-Fidelity DNA Polymerase (P505-d1) from Vazyme.

For each 50 µl reaction system, 15 µl of ddH2O was added along with 25 µl of 2× Phanta Max Buffer. Additionally, 1 µl of each 10 mM dATP, dTTP, dCTP, and dGTP was included. To this mixture, 2 µl of Forward Primer and 2 µl of Reverse Primer were added. Finally, 1 µl of Phanta Max Super-Fidelity DNA Polymerase (1 U/µl) and 1 µl of DNA were incorporated to complete the reaction mixture. Specifically, 5-Hydroxymethyl-dCTP (Cat. No. NU-932s) was purchased from Jena Bioscience as the alternative PCR substrate for dCTP. The PCR conditions were as follows: 1. Pre-denaturation at 95°C for 3 min, 2. Denaturation at 95°C for 15 seconds, 3. Annealing at 60°C for 15 seconds, 4. Extension at 72°C for 1.2 minutes or 1 minute per 1kb sequence, 5. Final Extension at 72°C for 5 minutes. After performing 30 cycles, the resulting products can be considered as a complete conversion from cytosine (C) to methylated 5hmC. To confirm the fidelity of amplification with DNA modification, the 5hmC PCR products were sent to Sanger sequencing showing identical sequences (Supplementary Figure 16).

### Electrical Readout via Nanopore Sequencing

Following PCR amplification, both the canonical (ATCG) and 5hmC-modified (AT^5hm^CG) DNA products were separately subjected to electrophoresis on a 1.5% agarose gel for length validation and purification. The bands corresponding to the reference sequence length were excised, followed by DNA extraction using the E.Z.N.A. Cycle Pure Kit (V-spin) from Omega Bio-Tek. A solution containing 200 ng of DNA was prepared using the ND608 kit from Vazyme for DNA damage repair, end preparation and adapter ligation. This includes two steps: 1) Damage repair and end preparation: 2 µl Damage Prep Enzyme, 5 µl End Prep Enzyme, and 10 µl End Prep Buffer were added, with ddH_2_O supplemented to a total volume of 65 µl. This mixture was incubated in a PCR thermal cycler at 30°C for 20 minutes, followed by 65°C for 15 minutes. 2) Adapter ligation: 25 µl Rapid Ligation Buffer 2, 5 µl Rapid DNA Ligase, and 5 µl AMX-F from the LSK-110 kit from Oxford Nanopore Technology were added to the reaction solution, and the mixture was incubated in a PCR thermal cycler at 20°C for 15 minutes.

Next, DNA purification was performed using VAHTS DNA Beads (N411-01) from Vazyme according to the bead purification protocol in the LSK-110 kit. The VAHTS DNA Beads were first resuspended by vortexing. Subsequently, 40 µl of the resuspended beads were added to the reaction in previous step and mixed by flicking the tube. The mixture was then incubated on a Hula mixer (rotator mixer) for 5 minutes at room temperature. After spinning down the sample and pelleting it on a magnet, the supernatant was pipetted off while keeping the tube on the magnet. The beads were then washed by adding 250 µl Short Fragment Buffer (SFB), flicking to resuspend, spinning down, and returning the tube to the magnetic rack to allow the beads to pellet. The supernatant was removed using a pipette and discarded. This washing step was repeated. After spinning down and placing the tube back on the magnet, residual supernatant was pipetted off and the beads were allowed to dry for about 30 seconds, ensuring the pellet did not crack. The tube was removed from the magnetic rack, and the pellet was resuspended in 15 µl Elution Buffer (EB). The mixture was spun down and incubated for 10 minutes at room temperature. The beads were then pelleted on a magnet until the eluate was clear and colorless, for at least 1 minute. Finally, 15 µl of eluate containing the DNA library was removed and retained in a clean 1.5 ml Eppendorf DNA LoBind tube.

Furthermore, DNA quantification was performed using a Qubit Fluorometer to measure 50 fmol of DNA. For flow cell priming and loading according to the LSK-110 protocol, 30 µl of thawed and mixed Flush Tether (FLT) was added directly to the tube of thawed and mixed Flush Buffer (FB) and mixed by vortexing at room temperature. Then, 800 µl of the priming mix was loaded into the flow cell (FLO-MIN106) via the priming port, avoiding the introduction of air bubbles, and left to sit for 5 minutes. During this time, the library for loading was prepared by mixing 37.5 µl Sequencing Buffer II (SBII), 25.5 µl Loading Beads II (LBII), and 12 µl DNA library including 50 fmol of DNA. An additional 200 µl of the priming mix was loaded into the flow cell via the priming port, again avoiding air bubbles. The prepared library was then mixed gently by pipetting up and down just before loading. Finally, 75 µl of the sample was added to the flow cell via the SpotON sample port in a dropwise fashion, ensuring each drop flowed into the port before adding the next.

Finally, the MinION was connected to a computer, and the MinKNOW v24.02.6 software was launched. The LSK-110 protocol and other default settings were selected. We also adjust the “sequence length cutoff options” on MinKNOW (200 bp for DNA Storage experiment, 1000 bp for 6-mer QC DeepSME Basecaller experiment). Finally, the sequencing button was clicked to begin collecting nanopore sequencing current data.

### Constructing of DeepSME Pipeline

#### Design and Synthesis of 6-mer QC sequence

The Cate_NAN plasmid (see Supplementary Table S17) of total length 5262 nt was used to constructing ten sequences of our 6-mer quality control (QC) dataset. Ten pairs of forward and reverse primers, ranging from 40 nt to 50 nt (see Supplementary Table S18), were designed and synthesized by Tsingke Biotech to generate corresponding ten samples of DNA sequences with length of 1330 nt, 1340 nt, 1340 nt, 1340 nt, 1340 nt, 1200 nt, 1200 nt, 1230 nt, 1226 nt and 1262 nt containing approximately 20 nt to 30 nt without cytosine (C) in their forward and reverse primers to serve as barcode of the sequence (see Supplementary Table S19). An additional sequence of 1144 nt was derived from a synthesized 1206 nt DNA template. These eleven key sequences contribute to 99.93% physical redundancy of 6-mers (4093/4096) with a median occurrence of three. The non-C segments on the eleven sequences were further used to facilitate subsequent data classification.

Both unmethylated and methylated PCR products were pooled and sequenced using an Oxford Nanopore Technology R9.4.1 flowcell with LSK-110 reagents. Specifically, the sequences were purified using the E.Z.N.A. Cycle Pure Kit (V-spin), followed by end repair and adapter ligation using the DNA Damage Repair Enzyme and End Prep Enzyme from the ND608 kit, and Rapid DNA Ligase, respectively. The final DNA samples were purified using VHATS DNA Beads, quantified with Qubit Flow, and loaded onto the flowcell for sequencing.

#### Preliminary Training of DeepSME with 6-mer datasets

The sequenced raw data were processed using Guppy 6.0 basecaller (`dna_r9.4.1_450bps_hac.cfg`) (https://nanoporetech.com/document/Guppy-protocol) as a control. The barcode region of basecalled fastq sequences were aligned back to the ionic currents using Tombo^50^ 1.5.1 (https://github.com/nanoporetech/tombo) to provide primer cutoff point. The current data were catalogued into 11 sequences, recognized by the non-C regions of the primers. These segments were also correctly recognized with 5hmC modification, ensuring automated classification of raw data into these 11 types of 6-mer QC sequences. The primer regions were then carefully trimmed, leaving only the 6-mer QC sequence regions.

Ionic current dataset from these 11 sequences were used for a preliminary training of a neural network with Encoder-Decoder architecture with Connectionist Temporal Classification (CTC) decoding layer (modified version of `bonito_bonitorev_ctc` based on `basecaller_benchmark` repository^48^, see supplementary information Table 14). The stride of the Encoder CNN layer was reconfigured to 1 to achieve high CTC layer resolution of DeepSME, resulting in the training convergence among 6-mer QC dataset in 151.8 minutes with an RTX 3090 GPU.

#### Extracting 6-mer statistics (k-mer model)

Ionic current data were segmented by the probability matrix obtained from the CTC segmentation^76^ of the trained 6-mer QC DeepSME, then enhanced by executing three times of k-mer extraction and alignment by Tombo to get the k-mer table (6-mer). Sequenced average currents, standard deviations as well as averaged dwell times and related standard deviations can be analyzed for either canonical or fully methylated 5-hmC sequenced by the R9.4.1 nanopore.

#### Expanding 6-mer to 9-mer with in silico simulations

Using the k-mer model from the previous step, simulated currents were generated for the whole genome fasta sequences of 50 microorganisms (Fasta can be download from https://github.com/marcpaga/nanopore_benchmark/blob/main/download/links_wick_data_tra in.txt) and simulated using Squigulator^56^ v0.3.0 (https://github.com/hasindu2008/squigulator) with a specified k-mer table csv file.

#### Enhanced training of DeepSME with Simulated Datasets

*In silico* experiments were performed to generate training datasets of 118,299 chunks with a chunk size of 3,600 (0.9 s) using the data preparation functionality of Bonito 0.8.1 (https://github.com/nanoporetech/bonito). These datasets were used to train the Encoder-Decoder architecture with Conditional Random Field (CRF) decoding layer (dna_r9.4.1_e8_sup@v3.3 model in Bonito 0.8.1, see supplementary Table. 15) over 5 epochs with a learning rate of 5e-4, resulting in the Enhanced DeepSME in 64.6 minutes with an RTX 3090 GPU.

#### Generating and Sequencing of Fully Methylated 5-hmC Bacteria Genome DNA (gDNA)

Genome DNA from Pseudomonas_aeruginosa_PAO1 (https://www.ncbi.nlm.nih.gov/datasets/genome/GCF_000006765.1/), Aeromonas_hydrophila_BJ054 (https://www.ncbi.nlm.nih.gov/datasets/genome/GCF_046708825.1/), and Vibrio_cholera_E1 (https://www.ncbi.nlm.nih.gov/datasets/genome/GCF_026013235.1/) microorganisms were extracted using the Qiagen Miniprep kit. The DNA was fragmented using the TD502 kit (TruePrep DNA Library Prep Kit V2 for Illumina) from Vazyme and modified using 5-Hydroxymethyl-dCTP (NU-932s) from Jena Bioscience. PCR was performed with Phanta Max Super-Fidelity DNA Polymerase (P505-d1) from Vazyme over 18 cycles under the following conditions of six steps: 1) Extension at 72°C for 3 minutes. 2) Pre-denaturation at 95°C for 3 min, 3) Denaturation at 95°C for 15 min, 4) Annealing at 60°C for 15 seconds, 5) Extension at 72°C for 60 s-72 s per kilobase sequence, 6) Final Extension at 72°C for 5 minutes. The modified DNA samples were sequenced using the Oxford Nanopore Technology R9.4.1 flowcell and LSK-110 reagents.

#### Reinforced DeepSME with Fully5-hmC Bacteria gDNA Datasets

The Enhanced DeepSME was used to basecall gDNA and map to the corresponding genome. The gDNA datasets from the three microbial samples also lead to 118,096 chunks with a chunk size of 3,600 (0.9 s). These data were then used to reinforce the Encoder-Decoder architecture with CRF decoding layer (dna_r9.4.1_e8_sup@v3.3 model in Bonito 0.8.1) over 5 epochs with a learning rate of 5e-4, yielding our final reinforced DeepSME, which takes 50.0 minutes with an RTX 3090 GPU. The weights of our basecallers could also be compatible with Dorado and be ready for deployed to edge devices like Jetson Xavier AGX for further use. We have also tried using FrodoKEM framework to encrypt our 5hmC-DeepSME basecaller weight with FrodoKEM-640-AES algorithm (see https://github.com/sparkcyf/FrodoKEM_demo) to demo the feasibility for integrating DeepSME with post-quantum cryptography.

### Recovering Data Stored in 5-hmC DNA using DeepSME

Dataset for digital information stored in DNA were prepared using our Composite Hedges Nanopore (CHN) codec^77^ in the four-letter configuration. The binary forms of two selected files (219 bytes and 4109 bytes) were extracted and split into payloads with 36 bytes. Next, Reed-Solomon (40,36) code was used to add redundancy to these payloads to generate segments. Then, a 34 nt barcode as addresses for random access followed with three 5-mer anchors are added to each segment. Finally, an oligo pool containing 487 single-stranded 243 nt DNA sequences was sent to Twist Biosciences, including 55 sequences that encode a .txt file and 432 sequence that encode a .jpeg image (see Supplementary Table 20 and Supplementary Table 21).

For the PCR amplification, a pair of carefully designed 23 nt 5’- and 11 nt 3’- flanking sequences was added to both ends of each DNA sequence. Canonical or modified 5hmC PCR were performed to obtain DNA sequences without or with modification, followed by above described nanopore sequencing procedure to obtain the current data.

The reinforced 5hmC DeepSME was used to process the sequenced raw data. The resulting fastq and aligned bam files were performed using the aboved-mentioned CHN decoder. For performance metrics, “the rate of the number of reads that could be aligned to the reference FASTA in total reads” was used as Recall, “the rate of the number of correct bases in total bases” was used as Precision, and their product was used as the F1-score.

## Data availability

The sequencing pod5 files from 5hmC-modified 6-mer QC sequences, Bacterial Genome DNA (gDNA) for training the 5hmC DeepSME basecaller and conducting the DNA storage decoding experiments have been deposited to Zenodo (https://doi.org/10.5281/zenodo.12704171). The FASTQ file generated by the 5hmC reinforced DeepSME from Bacterial Genome DNA (gDNA) has been deposited to NCBI Sequence Read Archive (SRA) accession number SRR32782578 (https://www.ncbi.nlm.nih.gov/sra/SRX28066745), SRR32782579 (https://www.ncbi.nlm.nih.gov/sra/SRX28066744), SRR32782580 (https://www.ncbi.nlm.nih.gov/sra/SRX28066743) under NCBI BioProject PRJNA1238011 (https://www.ncbi.nlm.nih.gov/bioproject/PRJNA1238011). The FASTQ file generated by the reinforced 5hmC DeepSME from the 5hmC DNA storage dataset has been deposited to NCBI Sequence Read Archive (SRA) accession number SRR32782578 (https://www.ncbi.nlm.nih.gov/sra/SRX28067610) under NCBI BioProject PRJNA1238011 (https://www.ncbi.nlm.nih.gov/bioproject/PRJNA1238011). All other data described in this work are available in the main text, provided in the supplementary materials, or can be reproduced using the deposited datasets and GitHub code (see “Code availavility” section). Source data are provided with this paper.

## Code availability

The code package for this study is available in the GitHub repository ( https://github.com/sparkcyf/DeepSME) and is also available on Zenodo under https://doi.org/10.5281/zenodo.15064695.

## Acknowledgements

This work was supported by the National Key Research and Development Program of China (no. 2022YFF1203400), National Natural Science Foundation of China (no. 62171211, 32371526, 32100021 and 32371372) and Science and Technology Innovation Commission of Shenzhen (JCYJ20220814170440001, JCYJ20220818100218039, JCYJ20220530113013030 and JCYJ20230807092459028), NSQKJJ under grant K21799109 and K21799116, Zhejiang Provincial Collaborative Innovation Center for High-end Digital Intelligence Diagnosis and Treatment Equipment and Center for Computational Science and Engineering at Southern University of Science and Technology.

## Author contributions

Q.Y.F. and Y.L. designed the experiment and the training pipeline. Q.Y.F., R.H.L. and Y.L. conducted the sequence design of the 6-mer QC sequence, Q.Y.F., R.H.L. Y.P.L. and J.X.Z. conducted the primers for the 6-mer QC sequence. M.L., Q.S.F. and Y.F. provided the plasmid samples for sequencing experiments. Q.Y.F. wrote the code for the k-mer extraction and the architecture of the preliminary basecaller. Q.Y.F., J.Y.L and X.Y.Z. conducted the nanopore basecalling and related quality check. Q.Y.F. conducted the construction of the datasets and training of the basecaller. Q.Y.F., X.Y.Z. and Y.L. prepared the figures and tables. Q.Y.F., Q.P. and Y.L. drafted the manuscript. Q.P. and Y.L. supervised the study. All authors read, revised, and approved the final manuscript.

