## Supplementary Information for "*De Novo* Non-Canonical Nanopore Basecalling Enables Private Communication using Heavily-modified DNA Data at Single-Molecule Level"

*Qingyuan Fan[1], Xuyang Zhao[1], Junyao Li[1], Ronghui Liu[1], Ming Liu[2], Qishun Feng[3], Yanping Long[4], Yang Fu[4], Jixian Zhai[4], Qing Pan[5], Yi Li[1,*]*

1. School of Microelectronics, MOE Engineering Research Center of Integrated Circuits for Next Generation Communications, Southern University of Science and Technology, 518055, Shenzhen, China

2. School of Medicine, Southern University of Science and Technology, 518055, Shenzhen, China

3. National Clinical Research Center for Infectious Diseases, Shenzhen Third People's Hospital, The Second Affiliated Hospital of Southern University of Science and Technology, Shenzhen, 518112, China

4. Department of Biology, School of Life Sciences, Southern University of Science and Technology, 518055, Shenzhen, China

5. College of Information Engineering, Zhejiang University of Technology, 310023, Hangzhou, China

 (Yi Li)

### Section I: Sequencing Length Distribution

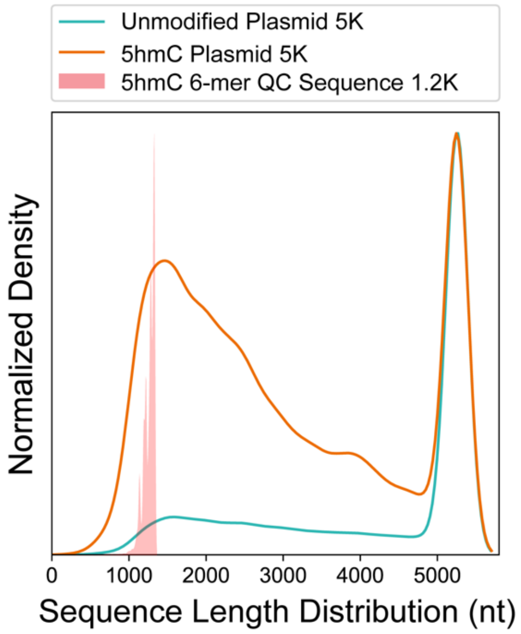

Figure S1 | Kernel density estimation (KDE) distribution of read length for Cate_NAN plasmid DNA (5262 bp) without and with 5hmC modification by nanopore sequencing (MinION R9.4.1).

The cyan line represents the read length distribution of unmodified Cate_NAN plasmid DNA (N=203,961 reads), while the orange line represents 5hmC-modified one (N=182,972 reads). The red peak indicates the read length distribution of the modified 6-mer quality check (QC) sequences (N=33,622 reads).

While nanopore basecallers and DeepSME are capable of processing long reads, we observed that heavy 5hmC modification tends to result in shorter sequenced reads (see supplementary Figure S1), potentially due to the modification's impact on the nanopore sequencing process itself, rather than limitations of the basecaller. For practical genome-scale applications, processing fragmented DNA of around 1 kilobases in size would align well with the current DeepSME framework and maintain efficient and accurate private communication.

### Section II: Translocation Speed

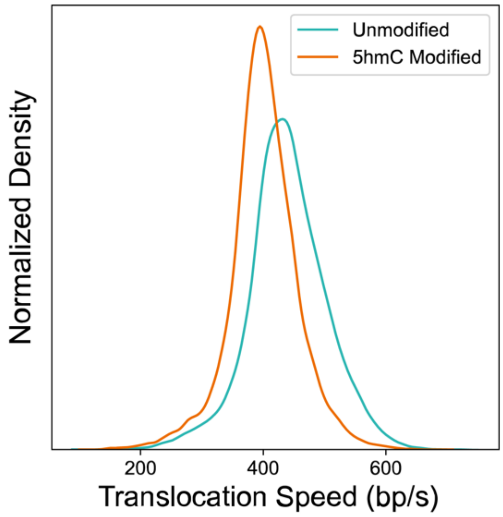

Figure S2 | Translocation speed distribution of Cate_NAN plasmid DNA without and with 5hmC modification.

The cyan line represents the translocation speed distribution of unmodified Cate_NAN plasmid DNA (N=20,000 reads), while the orange line represents the 5hmC-modified one (N=20,000 reads).

### Section III: Non-uniform Sequencing velocities

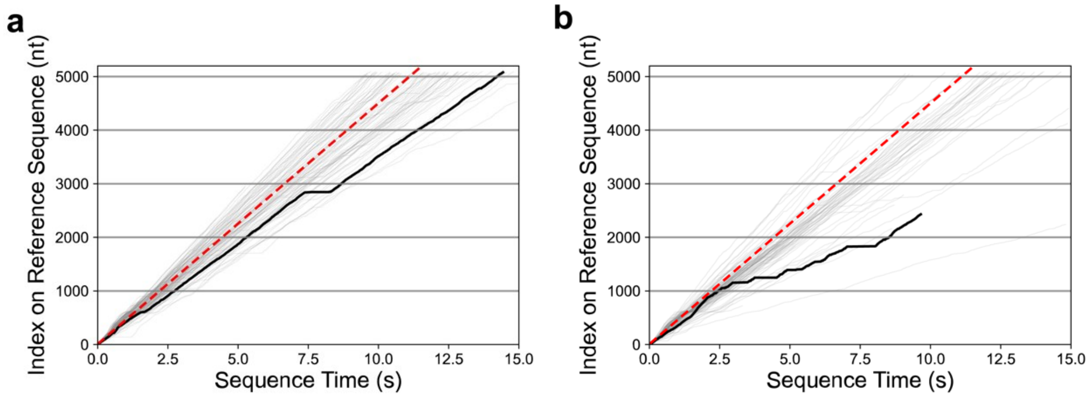

Figure S3 | Temporal nucleobase positions.

(**a**) Unmodified plasmid DNA strands. (**b**) 5hmC-modified plasmid DNA strands. The grey solid lines show 100 randomly selected DNA strands, while the black solid line highlights one of the representative DNA strands. The red dashed line represents a constant translocation speed of 450 nt/s.

5hmC-modified sequences are likely to "terminate at half a journey"(see Supplementary Figure S3). When sequencing a 5,262 nt long plasmid sequence, 57.9% of the unmodified sequences were longer than 5,100 nt. However, the value for extension exceeding 5,100 nt dropped to only 18.7% for the 5hmC-modified sequences. 4 folds of strands creased at 1,000 nt - 2,000 nt (see Supplementary Figure S1, S2). We speculate that this may be due to alteration in the chemical structure of 5hmC, which reduces the efficiency of helicase (motor protein) binding to the 5hmC methylated strands. Thus, the degradation on the binding strength leads to more sequences terminated much earlier before the finish line.

### Section IV: 9-mer Coverage Presentation for Datasets used in this work

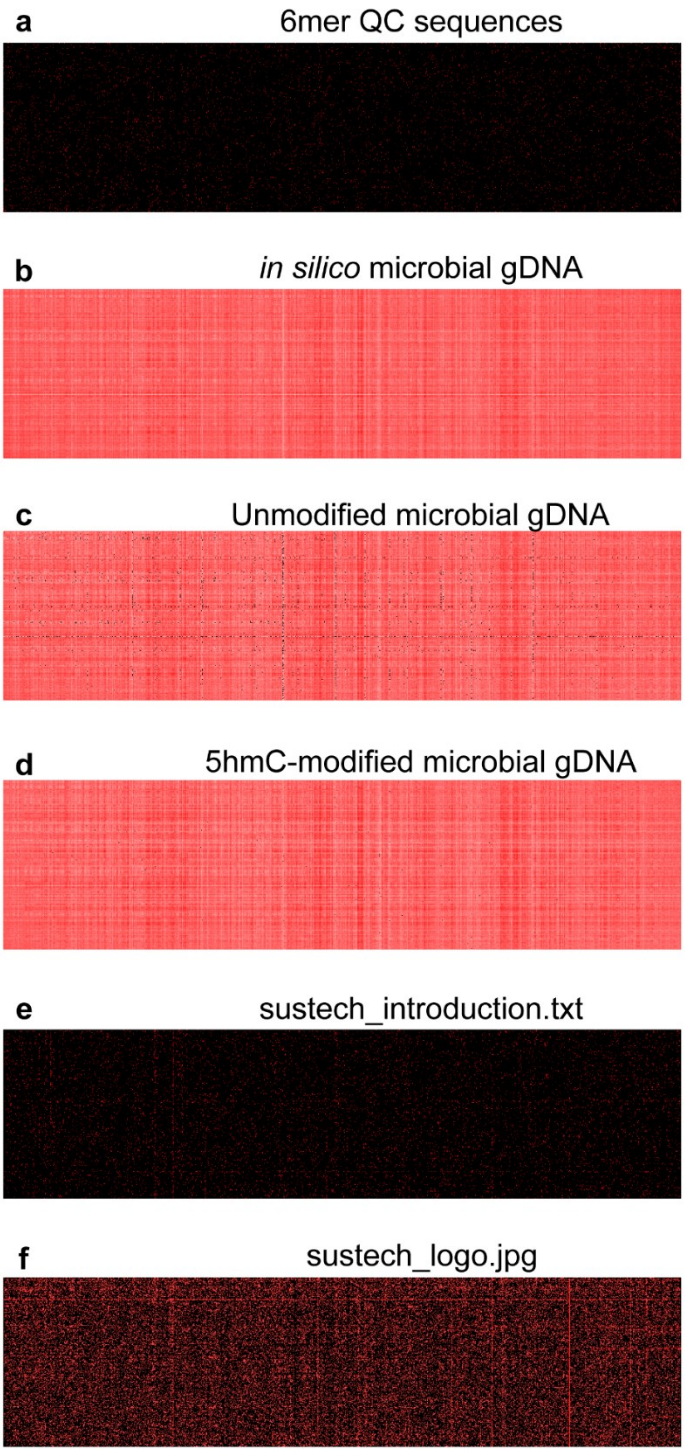

Figure S4 | Coverage depths for the datasets used in this work.

(**a**) 6-mer QC Sequence (**b**) *in silico* Microbial genome DNA (**c**) Unmodified Microbial genome DNA (**d**) 5hmC-modified Microbial genome DNA (**e**) CHN-coded text file (**f**) CHN-coded image file.

Table S5 | Datasets used in this work and their 9-mer coverages.

| FASTA Sequence | Usage | 9-mer Coverage |
|---|---|---|
| **6-mer QC Sequences** | Preliminary training | 2.40% |
| Simulated Sequences ( **https://github.com/marcpaga/nanopore_benchmark/blob/main/download/links_wick_data_train.txt** ) | Enhanced training | 100.00% |
| Experimental unmodified Sequences **(Klebsiella_variicola-KSB1_8J, Pseudomonas_aeruginosa-MINF_7A, Citrobacter_koseri-MINF_9D)** | Reinforced training | 99.12% |
| Experimental 5hmC Sequences **(Pseudomonas_aeruginosa_PAO1, Aeromonas_hydrophila_BJ054, and Vibrio_cholera_E1)** | Reinforced training | 99.96% |
| **sustech_introduction.txt 978 bytes, 55 strands** | Validation | 4.00% |
| sustech_logo.jpg 7,775 bytes, 432 strands | Validation | 26.43% |

### Section V: Cross-talking of Primer Classification

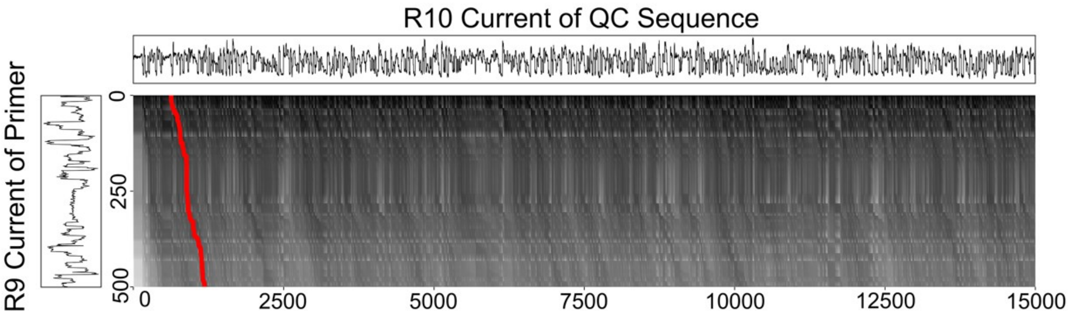

Figure S6 | Alignment between primer nanopore sequencing current from R9.4.1 Pore and simulated 6-mer QC sequence current from R10.4.1 Pore

This figure illustrates the possibility to align the primer and strand current among different types of nanopore.

### Section VI: Performance Metrics on DNA Storage Dataset

Table S7 | Precision, Recall and F1-score of 5hmC-modified DeepSME on 5hmC-modified DNA Storage Datasets

| DeepSME Basecaller | Precision | Recall | F1-score |
|---|---|---|---|
| **6-mer QC** | 0.0000 | 0.0000 | 0.0000 |
| **Enhanced** | 0.8287 | 0.0254 | 0.0211 |
| **Reinforced SUP** | **0.9299** | **0.9293** | **0.8642** |

Table S8 | Precision, Recall and F1-score of Unmodified DeepSME on Unmodified DNA Storage Datasets

| DeepSME Basecaller | Precision | Recall | F1-score |
|---|---|---|---|
| **6-mer QC** | 0.0000 | 0.0000 | 0.0000 |
| **Enhanced** | 0.8031 | 0.0338 | 0.0271 |
| **Reinforced SUP** | **0.9437** | **0.8810** | **0.8314** |

Table S9 | Precision, Recall and F1-score of Bonito on Unmodified DNA Storage Datasets

| Basecaller | Precision | Recall | F1-score |
|---|---|---|---|
| **Bonito SUP** | **0.9477** | **0.8945** | **0.8477** |
| **Bonito HAC** | 0.9426 | 0.8882 | 0.8372 |
| **Bonito FAST** | 0.9247 | 0.8525 | 0.7883 |

#### Table S10 | Precision, Recall and F1-score of different basecallers on 5hmC-modified DNA Storage Datasets

| Basecaller | Precision | Recall | F1-score |
|---|---|---|---|
| **Guppy 6.0** | 0.9294 | 0.0318 | 0.0296 |
| **Bonito SUP** | 0.9320 | 0.0428 | 0.0398 |
| **Dorado SUP** | 0.9535 | 0.0251 | 0.0239 |
| **Rerio 5mC 5hmC** | 0.9273 | 0.0278 | 0.0257 |
| **IL-AD 5hmC** | 0.8181 | 0.0306 | 0.0250 |
| **Reinforced DeepSME** | 0.9299 | 0.9293 | **0.8642** |

#### Table S11 | Precision, Recall and F1-score of 5hmC-modified DeepSME on Unmodified DNA Storage Datasets

| Basecaller | Precision | Recall | F1-score |
|---|---|---|---|
| **Reinforced SUP Bonito** | 0.9384 | 0.7723 | **0.7247** |

#### Table S12 | Performance of Reinforced DeepSME over different GPU platforms and software

| GPU | Basecaller Software | Precision | Recall | F1-score | Total Time (min)* | Throughput (samples/s)# |
|---|---|---|---|---|---|---|
| RTX3090 | Bonito | 0.9299 | **0.9293** | **0.8642** | 53.52 | 1.80×10$^{6}$ |
| | Dorado | 0.9299 | 0.9278 | 0.8628 | **23.58** | **3.98×10$^{6}$** |
| Xavier | | 0.9299 | 0.9278 | 0.8628 | 250.90 | 3.75×10$^{5}$ |

* $1.90 \times 10^6$ samples in this validation 5hmC-modified DNA storage datasets.
### The sampling rate is 4000 samples per second per channel.

Table S13 | Performance of Reinforced DeepSME over different GPU platforms and software

| | Cost | | | Performance | | |
|---|---|---|---|---|---|---|
| | Total Time | GPU Memory* | GPU Power | Precision | Recall | F1-score |
| Bonito v0.8.1 | 53.52 min | 14.02 GB | **220 W** | 0.9299 | **0.9293** | **0.8642** |
| Dorado v0.9.1 | **23.58 min** | **10.74 GB** | 250 W | 0.9299 | 0.9278 | 0.8628 |

* GPU Memory and GPU Power are measured when batchsize=768, chunksize=3600.

### Section VII: Model Architectures

#### Table S14 | Model architecture of DeepSME Preliminary 6-mer QC Basecaller

```
==================================================================================
Layer (type:depth-idx)                   Output Shape              Param #
==================================================================================
BonitoModel                              [2000, 64, 5]             --
├─Sequential: 1-1                        [64, 384, 2000]           --
│    └─Conv1d: 2-1                       [64, 4, 2000]             24
│    └─SiLU: 2-2                         [64, 4, 2000]             --
│    └─Conv1d: 2-3                       [64, 16, 2000]            336
│    └─SiLU: 2-4                         [64, 16, 2000]            --
│    └─Conv1d: 2-5                       [64, 384, 2000]           117,120
│    └─SiLU: 2-6                         [64, 384, 2000]           --
├─Sequential: 1-2                        [2000, 64, 384]           --
│    └─BonitoLSTM: 2-7                   [2000, 64, 384]           --
│    │    └─LSTM: 3-1                    [2000, 64, 384]           1,182,720
│    └─BonitoLSTM: 2-8                   [2000, 64, 384]           --
│    │    └─LSTM: 3-2                    [2000, 64, 384]           1,182,720
│    └─BonitoLSTM: 2-9                   [2000, 64, 384]           --
│    │    └─LSTM: 3-3                    [2000, 64, 384]           1,182,720
│    └─BonitoLSTM: 2-10                  [2000, 64, 384]           --
│    │    └─LSTM: 3-4                    [2000, 64, 384]           1,182,720
│    └─BonitoLSTM: 2-11                  [2000, 64, 384]           --
│    │    └─LSTM: 3-5                    [2000, 64, 384]           1,182,720
├─Sequential: 1-3                        [2000, 64, 5]             --
│    └─Linear: 2-12                      [2000, 64, 5]             1,925
│    └─LogSoftmax: 2-13                  [2000, 64, 5]             --
==================================================================================
Total params: 6,033,005
Trainable params: 6,033,005
Non-trainable params: 0
Total mult-adds (Units.GIGABYTES): 771.98
==================================================================================
Input size (MB): 0.51
Forward/backward pass size (MB): 2384.90
Params size (MB): 24.13
Estimated Total Size (MB): 2409.54
==================================================================================
```

*Windows Size 2000, Batch Size 64*

### Table S15 | Model Architecture of DeepSME Enhanced and Reinforced Basecallers

```
==============================================================================
Layer (type:depth-idx)                    Output Shape              Param #
==============================================================================
Model                                     [64, 720, 4096]           --
├─Sequential: 1-1                         [64, 720, 4096]           --
│    └─Convolution: 2-1                   [64, 4, 3600]             --
│    │    └─Conv1d: 3-1                   [64, 4, 3600]             24
│    │    └─Swish: 3-2                    [64, 4, 3600]             --
│    └─Convolution: 2-2                   [64, 16, 3600]            --
│    │    └─Conv1d: 3-3                   [64, 16, 3600]            336
│    │    └─Swish: 3-4                    [64, 16, 3600]            --
│    └─Convolution: 2-3                   [64, 768, 720]            --
│    │    └─Conv1d: 3-5                   [64, 768, 720]            234,240
│    │    └─Swish: 3-6                    [64, 768, 720]            --
│    └─Permute: 2-4                       [720, 64, 768]            --
│    └─LSTM: 2-5                          [720, 64, 768]            --
│    │    └─LSTM: 3-7                     [720, 64, 768]            4,724,736
│    └─LSTM: 2-6                          [720, 64, 768]            --
│    │    └─LSTM: 3-8                     [720, 64, 768]            4,724,736
│    └─LSTM: 2-7                          [720, 64, 768]            --
│    │    └─LSTM: 3-9                     [720, 64, 768]            4,724,736
│    └─LSTM: 2-8                          [720, 64, 768]            --
│    │    └─LSTM: 3-10                    [720, 64, 768]            4,724,736
│    └─LSTM: 2-9                          [720, 64, 768]            --
│    │    └─LSTM: 3-11                    [720, 64, 768]            4,724,736
│    └─Permute: 2-10                      [64, 720, 768]            --
│    └─LinearCRFEncoder: 2-11             [64, 720, 4096]           --
│    │    └─Linear: 3-12                  [64, 720, 4096]           3,149,824
│    │    └─Tanh: 3-13                    [64, 720, 4096]           --
==============================================================================
Total params: 27,008,104
Trainable params: 26,992,744
Non-trainable params: 15,360
Total mult-adds (T): 1.10
==============================================================================
Input size (MB): 0.46
Forward/backward pass size (MB): 1622.75
Params size (MB): 54.02
Estimated Total Size (MB): 1677.23
==============================================================================
```

*Windows Size 3600, Batch Size 64*

### Section VIII Sanger Sequencing Results for 5hmC modified 6-mer QC Sequence

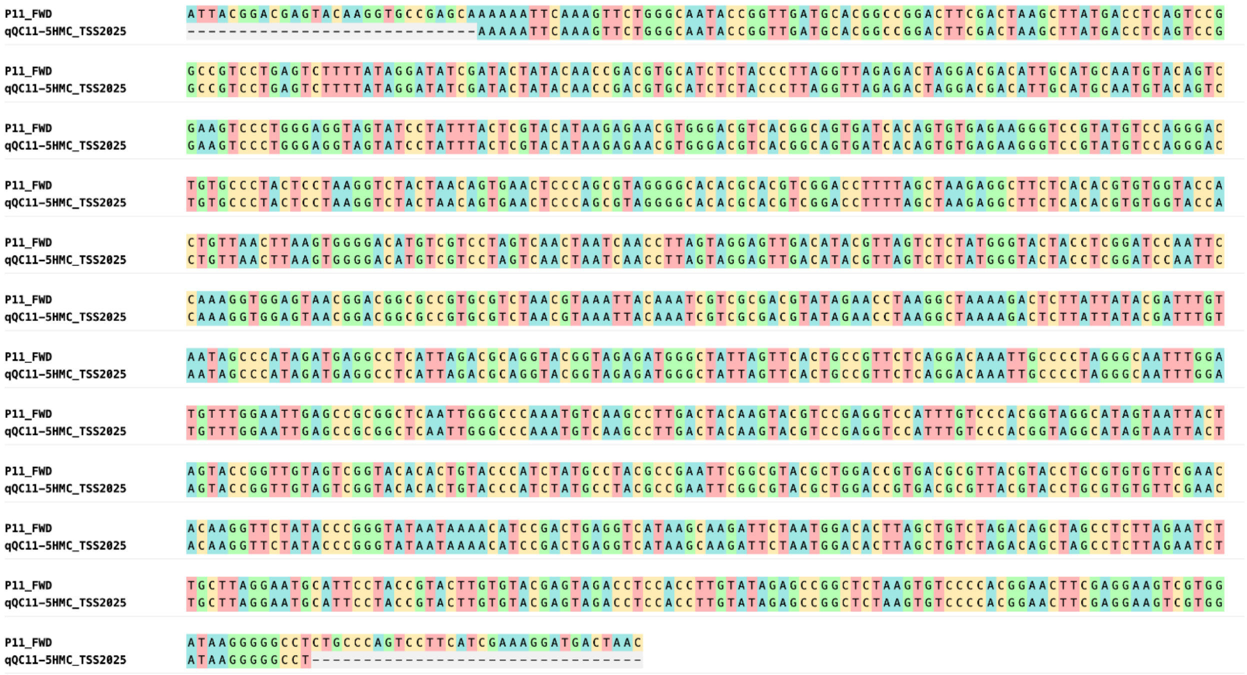

Figure S16 | Alignment results for P11_FWD (one of the 6-mer QC Sequences) and Sanger sequencing result of 5hmC modified P11_FWD sequence (qQC11-5HMC_TSS2025) from Tsingke Technology.

### Section IX FASTA Sequence of Cate_NAN Plasmid and synthesis sequence

#### Table S17 | Cate_NAN and synthesis sequence

```
>Cate_NAN
TGCCGTTCGCGTTATAAATCGGCTGACCCGCCGCATCGGTGCGATAGGTAATATCATCCGGTTCTTTGCGGGTGCGCGCTTGTTCCGCGCCCACCGCCAGCGCGCCAATCGGGG
TGCTGTTGGTAAACTGCACCATGCCGCCTTCTTCGCTAATCCACTGCACGCCCAGGTTAAAGCCATCGCCTTCAAACACTTCCACAATAATCGCTTCCACTTGCACTTGCGCGC
GGCGAATATCCAGCTGACGAATCACCGCTTCCAGGCTGCGCATCATATCCGGTTCCGCGGTAATCACCAGCGCGTTGCTATCCGGATGCGCATCAATGCTCACATCGCGCGCGC
TACTGCTACTGCTGCGTTTGTTGCTGCCGCCCGCTTTATCTTCCGCAATGCTACTGCTCACGCCTTGCAGCACTTTCACCAGATCTTCGGCCTTCGCATATTTCAGATAATACA
CTTTGGTGTTGCCGTTGGTTTCCAGTTCGCTATCCAGGCGCATAATCATATCAATGGTGCGCTGACGCGCTTTGCCTTCGCCGCTCACAATCACGCTGTTGGTGCGGTCATCCG
CCACCACTTTCGGAATCAGGTTATCCGGGGTGCGATCTTTCGCGCTGTTTTTGGTCAGGGTATCAATAATGCGCACCATTTCGCTCGCGCTCGCATATTTTAATTTCACAATTT
CCACTTCTTGATCCATGGCCATCGCCGGCTGGGCAGCGAGGAGCAGCAGACCAGCAGCAGCGGTCGGCAGCAGGTATTTCATATGTATATCTCCTTCTTAAAGTTAAACAAAAT
TATTTCTAGAGGGAAACCGTTGTGGTCTCCCTATAGTGAGTCGTATTAATTTCGCGGGATCGAGATCTCGGGCAGCGTTGGGTCCTGGCCACGGGTGCGCATGATCGTGCTCCT
GTCGTTGAGGACCCGGCTAGGCTGGCGGGGTTGCCTTACTGGTTAGCAGAATGAATCACCGATACGCGAGCGAACGTGAAGCGACTGCTGCTGCAAAACGTCTGCGACCTGAGC
AACAACATGAATGGTCTTCGGTTTCCGTGTTTCGTAAAGTCTGGAAACGCGGAAGTCAGCGCCCTGCACCATTATGTTCCGGATCTGCATCGCAGGATGCTGCTGGCTACCCTG
TGGAACACCTACATCTGTATTAACGAAGCGCTGGCATTGACCCTGAGTGATTTTTCTCTGGTCCCGCCGCATCCATACCGCCAGTTGTTTACCCTCACAACGTTCCAGTAACCG
GGCATGTTCATCATCAGTAACCCGTATCGTGAGCATCCTCTCTCGTTTCATCGGTATCATTACCCCCATGAACAGAAATCCCCCTTACACGGAGGCATCAGTGACCAAACAGGA
AAAAACCGCCCTTAACATGGCCCGCTTTATCAGAAGCCAGACATTAACGCTTCTGGAGAAACTCAACGAGCTGGACGCGGATGAACAGGCAGACATCTGTGAATCGCTTCACGA
CCACGCTGATGAGCTTTACCGCAGCTGCCTCGCGCGTTTCGGTGATGACGGTGAAAACCTCTGACACATGCAGCTCCCGGAGACGGTCACAGCTTGTCTGTAAGCGGATGCCGG
GAGCAGACAAGCCCGTCAGGGCGCGTCAGCGGGTGTTGGCGGGTGTCGGGGCGCAGCCATGACCCAGTCACGTAGCGATAGCGGAGTGTATACTGGCTTAACTATGCGGCATCA
GAGCAGATTGTACTGAGAGTGCACCATATATGCGGTGTGAAATACCGCACAGATGCGTAAGGAGAAAATACCGCATCAGGCGCTCTTCCGCTTCCTCGCTCACTGACTCGCTGC
GCTCGGTCGTTCGGCTGCGGCGAGCGGTATCAGCTCACTCAAAGGCGGTAATACGGTTATCCACAGAATCAGGGGATAACGCAGGAAAGAACATGTGAGCAAAAGGCCAGCAAA
AGGCCAGGAACCGTAAAAAGGCCGCGTTGCTGGCGTTTTTCCATAGGCTCCGCCCCCCTGACGAGCATCACAAAAATCGACGCTCAAGTCAGAGGTGGCGAAACCCGACAGGAC
TATAAAGATACCAGGCGTTTCCCCCTGGAAGCTCCCTCGTGCGCTCTCCTGTTCCGACCCTGCCGCTTACCGGATACCTGTCCGCCTTTCTCCCTTCGGGAAGCGTGGCGCTTT
CTCATAGCTCACGCTGTAGGTATCTCAGTTCGGTGTAGGTCGTTCGCTCCAAGCTGGGCTGTGTGCACGAACCCCCCGTTCAGCCCGACCGCTGCGCCTTATCCGGTAACTATC
GTCTTGAGTCCAACCCGGTAAGACACGACTTATCGCCACTGGCAGCAGCCACTGGTAACAGGATTAGCAGAGCGAGGTATGTAGGCGGTGCTACAGAGTTCTTGAAGTGGTGGC
CTAACTACGGCTACACTAGAAGGACAGTATTTGGTATCTGCGCTCTGCTGAAGCCAGTTACCTTCGGAAAAAGAGTTGGTAGCTCTTGATCCGGCAAACAAACCACCGCTGGTA
GCGGTGGTTTTTTTGTTTGCAAGCAGCAGATTACGCGCAGAAAAAAAGGATCTCAAGAAGATCCTTTGATCTTTTCTACGGGGTCTGACGCTCAGTGGAACGAAAACTCACGTT
AAGGGATTTTGGTCATGAGATTATCAAAAAGGATCTTCACCTAGATCCTTTTAAATTAAAAATGAAGTTTTAAATCAATCTAAAGTATATATGAGTAAACTTGGTCTGACAGTT
ACCAATGCTTAATCAGTGAGGCACCTATCTCAGCGATCTGTCTATTTCGTTCATCCATAGTTGCCTGACTCCCCGTCGTGTAGATAACTACGATACGGGAGGGCTTACCATCTG
GCCCCAGTGCTGCAATGATACCGCGAGACCCACGCTCACCGGCTCCAGATTTATCAGCAATAAACCAGCCAGCCGGAAGGGCCGAGCGCAGAAGTGGTCCTGCAACTTTATCCG
CCTCCATCCAGTCTATTAATTGTTGCCGGGAAGCTAGAGTAAGTAGTTCGCCAGTTAATAGTTTGCGCAACGTTGTTGCCATTGCTGCAGGCATCGTGGTGTCACGCTCGTCGT
TTGGTATGGCTTCATTCAGCTCCGGTTCCCAACGATCAAGGCGAGTTACATGATCCCCCATGTTGTGCAAAAAAGCGGTTAGCTCCTTCGGTCCTCCGATCGTTGTCAGAAGTA
```

```
AGTTGGCCGCAGTGTTATCACTCATGGTTATGGCAGCACTGCATAATTCTCTTACTGTCATGCCATCCGTAAGATGCTTTTCTGTGACTGGTGAGTACTCAACCAAGTCATTCT
GAGAATAGTGTATGCGGCGACCGAGTTGCTCTTGCCCGGCGTCAATACGGGATAATACCGCGCCACATAGCAGAACTTTAAAAGTGCTCATCATTGGAAAACGTTCTTCGGGGC
GAAAACTCTCAAGGATCTTACCGCTGTTGAGATCCAGTTCGATGTAACCCACTCGTGCACCCAACTGATCTTCAGCATCTTTTACTTTCACCAGCGTTTCTGGGTGAGCAAAAA
CAGGAAGGCAAAATGCCGCAAAAAAGGGAATAAGGGCGACACGGAAATGTTGAATACTCATACTCTTCCTTTTTCAATATTATTGAAGCATTTATCAGGGTTATTGTCTCATGA
GCGGATACATATTTGAATGTATTTAGAAAAATAAACAAATAGGGGTTCCGCGCACATTTCCCCGAAAAGTGCCACCTGAAATTGTAAACGTTAATATTTTGTTAAAATTCGCGT
TAAATTTTTGTTAAATCAGCTCATTTTTTAACCAATAGGCCGAAATCGGCAAAATCCCTTATAAATCAAAAGAATAGACCGAGATAGGGTTGAGTGTTGTTCCAGTTTGGAACA
AGAGTCCACTATTAAAGAACGTGGACTCCAACGTCAAAGGGCGAAAAACCGTCTATCAGGGCGATGGCCCACTACGTGAACCATCACCCTAATCAAGTTTTTTGGGGTCGAGGT
GCCGTAAAGCACTAAATCGGAACCCTAAAGGGAGCCCCCGATTTAGAGCTTGACGGGGAAAGCCGGCGAACGTGGCGAGAAAGGAAGGGAAGAAAGCGAAAGGAGCGGGCGCTA
GGGCGCTGGCAAGTGTAGCGGTCACGCTGCGCGTAACCACCACACCCGCCGCGCTTAATGCGCCGCTACAGGGCGCGTCCCATTCGCCAATCCGGATATAGTTCCTCCTTTCAG
CAAAAAACCCCTCAAGACCCGTTTAGAGGCCCCAAGGGGTTATGCTAGTTATTGCTCAGCGGTGGCAGCAGCCAACTCAGCTTCCTTTCGGGCTTTGTTAGCAGCCGGATCTCA
GTGGTGGTGGTGGTGGTGCTCGAGTGCGGCCGCAAGCTTGTCGACGGAGCTCGAATTcTCATTTTTCGAACTGCGGGTGGCTCCAAGCAGAATCGCGCTGCGCTTCATCTTTTT
TCAGATATTCATCCAGCGCGCCTTTTTTAATCAGATAATCTTCAAAGCTCGGCGGCAGCGCCATGCTATCATCCCAATCCGGCAGCACGCTAATATCATCGCTATAGCTCAGAA
AGTTGCCGTTCGCGCGGCGATCCAGCTGCTGCGCGCGCATAAAGTTATATTTGCGATGGCTAATTTTGTTCTGACTCAGGCCATCGCGAATAATGGTCGGGCGAATAAACACCA
TCAGGTTGCGTTTGCGTTTGGTGTTGCTCGTGCTTTTAAACAGGTTGCCAATAATCGGAATATCGCCCAGCAGCGGCACTTTGCTCACGCTTTCTTGAATATCTTCATCAATCA
GGCCGCCCAGCACAATGGTGCCGCCATCTTCCGCCATCACGGTGGTTTTAATTTCGCGTTTGTTAAAGCTCACATCAATGCCGGTCGCGCCCGCCACGCTGCTCACTTCTTGTT
CAATGGTCAGCATCACCGCATCGCCTTCGTTAATCTGCGGGGTCACTTTCAGTTTAATGCCCACTTCTTTGCGTTCCACGGTCTGAAACGGATTCGTGTTGCTACTGCTCGCGG
TCGCGCCGGTAATGGTCGGCACTTCTTGGCCCACAATAAAGCTCGCTTCTTGGTTATCCAGGGTGGTCAGGCTCGGGGTCGCCAGAATGTTGCTGTTGGTCGCGGTGGTCACCG
CTTGCAGCACCGCGCCCCAGCCATCTTTAATCACGCCCACCATCGTGCCGTTAATGCCGCCCAGCAGGCTCGCCAGGGTGCCAAAATCGCCTTGAATTTCTTTGCCTTCGGTAA
TGCTCGCAATCACGCGGT
>6mer_fill_syn_seq
CCATGGGTCGGCTGGGCCGTCATTACGGACGAGTACAAGGTGCCGAGCAAAAAATTCAAAGTTCTGGGCAATACCGGTTGATGCACGGCCGGACTTCGACTAAGCTTATGACCT
CAGTCCGGCCGTCCTGAGTCTTTTATAGGATATCGATACTATACAACCGACGTGCATCTCTACCCTTAGGTTAGAGACTAGGACGACATTGCATGCAATGTACAGTCGAAGTCC
CTGGGAGGTAGTATCCTATTTACTCGTACATAAGAGAACGTGGGACGTCACGGCAGTGATCACAGTGTGAGAAGGGTCCGTATGTCCAGGGACTGTGCCCTACTCCTAAGGTCT
ACTAACAGTGAACTCCCAGCGTAGGGGCACACGCACGTCGGACCTTTTAGCTAAGAGGCTTCTCACACGTGTGGTACCACTGTTAACTTAAGTGGGGACATGTCGTCCTAGTCA
ACTAATCAACCTTAGTAGGAGTTGACATACGTTAGTCTCTATGGGTACTACCTCGGATCCAATTCCAAAGGTGGAGTAACGGACGGCGCCGTGCGTCTAACGTAAATTACAAAT
CGTCGCGACGTATAGAACCTAAGGCTAAAAGACTCTTATTATACGATTTGTAATAGCCCATAGATGAGGCCTCATTAGACGCAGGTACGGTAGAGATGGGCTATTAGTTCACTG
CCGTTCTCAGGACAAATTGCCCCTAGGGCAATTTGGATGTTTGGAATTGAGCCGCGGCTCAATTGGGCCCAAATGTCAAGCCTTGACTACAAGTACGTCCGAGGTCCATTTGTC
CCACGGTAGGCATAGTAATTACTAGTACCGGTTGTAGTCGGTACACACTGTACCCATCTATGCCTACGCCGAATTCGGCGTACGCTGGACCGTGACGCGTTACGTACCTGCGTG
TGTTCGAACACAAGGTTCTATACCCGGGTATAATAAAACATCCGACTGAGGTCATAAGCAAGATTCTAATGGACACTTAGCTGTCTAGACAGCTAGCCTCTTAGAATCTTGCTT
AGGAATGCATTCCTACCGTACTTGTGTACGAGTAGACCTCCACCTTGTATAGAGCCGGCTCTAAGTGTCCCCACGGAACTTCGAGGAAGTCGTGGATAAGGGGGCCTCTGCCCA
GTCCTTCATCGAAAGGATGACTAACTTTGATAAAAATCTGCCTAACGAAAAGGTGCTTCCCTCGAG
```

### Table S18 | Primers

```
>seq1_fwd
GTGTAGATGATAGGGGTGGTGTTAAGAGGTGGAAATGAGTTGCCGTTCGCGTTA
TAAATCGGCTGACCCG
>seq1_rev
AACACAAATCAACTTCCAACTAACAAATCTTTCCACTAACTACTGGAACGTTGT
GAGGGTAAACAACTGG
>seq2_fwd
TTGAGAATAGAGTAAAGGGAGTTGAGAGGAGGGGTAGGGTGGTGTTGGAGAGCG
AACGTGAAGCGACTGC
>seq2_rev
ATTCTCTATTCACAAAACTTCACTCCCATTACACATACCTCCTCCTTACCGCTT
GGAGCGAACGACCTAC
>seq3_fwd
GTAATAGGGATTAAGAGGTTTGTGAAGTTTAGTGGGGTGAAATGTGGGGAATAG
GCTCCGCCCCCCTGAC
>seq3_rev
ATCTCCTAACTCATCCACCCTTTACTCACACATTTCCTAAATCCTCTATTACCA
TGAGTGATAACACTGC
>seq4_fwd
GTAGAATGGATGGGAAATGGGAGTTGTTATAGGGGATGTTTCCAGTCTATTAAT
TGTTGCCGGGAAGCTA
>seq4_rev
TACATCTATTCCCACACTTCTCCATCCATTCTCCACCTCCAGGGGTTTTTTGCT
GAAAGGAGGAACTATA
>seq5_fwd
GAAGTGTATATTTGGGGTGGGAGGTGGAGAATGGATGGAGAAGTGTGGGAGGGG
TCGAGGTGCCGTAAAG
>seq5_rev
ACCTCTTAATCCCTATTACCTCCAACACCACCCTACCCCTCCTCTCAACTCGAT
TTTGGCACCCTGGCGA
>seq6_fwd
GTGGTGTTAAGAGGTGGAAATGAGTTTGAGAATAGAGTAAAGGGAGTTGAACCG
CGTGATTGCGAGCATTACCGAAGGCA
```

```
>seq6_rev
TCCAACTAACAAATCTTTCCACTAACATTCTCTATTCACAAAACTTCACTTGGC
GAGAAAGGAAGGGAAGAAAGCGAAAG
>seq7_fwd
GAGGAGGGGTAGGGTGGTGTTGGAGGTAATAGGGATTAAGAGGTTTGTGACATA
ACCCCTTGGGGCCTCTAAACGGGTCT
>seq7_rev
CCCATTACACATACCTCCTCCTTACCATCTCCTAACTCATCCACCCTTTAGTGT
CACGCTCGTCGTTTGGTATGGCTTCA
>seq8_fwd
AGTTTAGTGGGGTGAAATGTGGGGAATATATGGTGGTAGAATGGATGGGAACGG
ATGGCATGACAGTAAGAGAATTATGC
>seq8_rev
CTCACACATTTCCTAAATCCTCTATTCAACCCTCCATACATCTATTCCCAGTGG
CGAAACCCGACAGGACTATAAAGATA
>seq9_fwd
AATGGGAGTTGTTATAGGGGATGTTGAAGTGTATATTTGGGGTGGGAGGTCTCA
AGACGATAGTTACCGGATAAGGCGC
>seq9_rev
CACTTCTCCATCCATTCTCCACCTCCCACCCCAAATATACACTTCAACATAAGT
CTGGAAACGCGGAAGTCAGCGCCCTG
>seq10_fwd
GGAGAATGGATGGAGAAGTGTGGGAATAGATGTATGGAGGGTTGAATAGAAACA
TGCCCGGTTACTGGAACGTTGTGAGG
>seq10_rev
ACATTTCACCCCACTAAACTTCACAAACCTCTTAATCCCTATTACCTCCATGCC
GTTCGCGTTATAAATCGGCTGACCCG
>seq11_fwd
GGATTTAGGAAATGTGTGAGTAAAGGGTGGATGAGTTAGGAGATGGTAAGATTA
CGGACGAGTACAAGGTGCCGAGCAAA
>seq11_rev
ACACCACCCTACCCCTCCTCTCAACTCCCTTTACTCTATTCTCAAACTCAGTTA
GTCATCCTTTCGATGAAGGACTGGGC
```

#### Table S19 | 6-mer QC Sequence After PCR

```
>P1_FWD
GTGTAGATGATAGGGGTGGTGTTAAGAGGTGGAAATGAGTTGCCGTTCGCGTTATAAATCGGCTGACCCGCCGCATCGGTGCGATAGGTAATATCATCCGGTTCTTTGCGGGTG
CGCGCTTGTTCCGCGCCCACCGCCAGCGCGCCAATCGGGGTGCTGTTGGTAAACTGCACCATGCCGCCTTCTTCGCTAATCCACTGCACGCCCAGGTTAAAGCCATCGCCTTCA
AACACTTCCACAATAATCGCTTCCACTTGCACTTGCGCGCGGCGAATATCCAGCTGACGAATCACCGCTTCCAGGCTGCGCATCATATCCGGTTCCGCGGTAATCACCAGCGCG
TTGCTATCCGGATGCGCATCAATGCTCACATCGCGCGCGCTACTGCTACTGCTGCGTTTGTTGCTGCCGCCCGCTTTATCTTCCGCAATGCTACTGCTCACGCCTTGCAGCACT
TTCACCAGATCTTCGGCCTTCGCATATTTCAGATAATACACTTTGGTGTTGCCGTTGGTTTCCAGTTCGCTATCCAGGCGCATAATCATATCAATGGTGCGCTGACGCGCTTTG
CCTTCGCCGCTCACAATCACGCTGTTGGTGCGGTCATCCGCCACCACTTTCGGAATCAGGTTATCCGGGGTGCGATCTTTCGCGCTGTTTTTGGTCAGGGTATCAATAATGCGC
ACCATTTCGCTCGCGCTCGCATATTTTAATTTCACAATTTCCACTTCTTGATCCATGGCCATCGCCGGCTGGGCAGCGAGGAGCAGCAGACCAGCAGCAGCGGTCGGCAGCAGG
TATTTCATATGTATATCTCCTTCTTAAAGTTAAACAAAATTATTTCTAGAGGGAAACCGTTGTGGTCTCCCTATAGTGAGTCGTATTAATTTCGCGGGATCGAGATCTCGGGCA
GCGTTGGGTCCTGGCCACGGGTGCGCATGATCGTGCTCCTGTCGTTGAGGACCCGGCTAGGCTGGCGGGGTTGCCTTACTGGTTAGCAGAATGAATCACCGATACGCGAGCGAA
CGTGAAGCGACTGCTGCTGCAAAACGTCTGCGACCTGAGCAACAACATGAATGGTCTTCGGTTTCCGTGTTTCGTAAAGTCTGGAAACGCGGAAGTCAGCGCCCTGCACCATTA
TGTTCCGGATCTGCATCGCAGGATGCTGCTGGCTACCCTGTGGAACACCTACATCTGTATTAACGAAGCGCTGGCATTGACCCTGAGTGATTTTTCTCTGGTCCCGCCGCATCC
ATACCGCCAGTTGTTTACCCTCACAACGTTCCAGTAGTTAGTGGAAAGATTTGTTAGTTGGAAGTTGATTTGTGTT
>P2_FWD
TTGAGAATAGAGTAAAGGGAGTTGAGAGGAGGGGTAGGGTGGTGTTGGAGAGCGAACGTGAAGCGACTGCTGCTGCAAAACGTCTGCGACCTGAGCAACAACATGAATGGTCTT
CGGTTTCCGTGTTTCGTAAAGTCTGGAAACGCGGAAGTCAGCGCCCTGCACCATTATGTTCCGGATCTGCATCGCAGGATGCTGCTGGCTACCCTGTGGAACACCTACATCTGT
ATTAACGAAGCGCTGGCATTGACCCTGAGTGATTTTTCTCTGGTCCCGCCGCATCCATACCGCCAGTTGTTTACCCTCACAACGTTCCAGTAACCGGGCATGTTCATCATCAGT
AACCCGTATCGTGAGCATCCTCTCTCGTTTCATCGGTATCATTACCCCCATGAACAGAAATCCCCCTTACACGGAGGCATCAGTGACCAAACAGGAAAAAACCGCCCTTAACAT
GGCCCGCTTTATCAGAAGCCAGACATTAACGCTTCTGGAGAAACTCAACGAGCTGGACGCGGATGAACAGGCAGACATCTGTGAATCGCTTCACGACCACGCTGATGAGCTTTA
CCGCAGCTGCCTCGCGCGTTTCGGTGATGACGGTGAAAACCTCTGACACATGCAGCTCCCGGAGACGGTCACAGCTTGTCTGTAAGCGGATGCCGGGAGCAGACAAGCCCGTCA
GGGCGCGTCAGCGGGTGTTGGCGGGTGTCGGGGCGCAGCCATGACCCAGTCACGTAGCGATAGCGGAGTGTATACTGGCTTAACTATGCGGCATCAGAGCAGATTGTACTGAGA
GTGCACCATATATGCGGTGTGAAATACCGCACAGATGCGTAAGGAGAAAATACCGCATCAGGCGCTCTTCCGCTTCCTCGCTCACTGACTCGCTGCGCTCGGTCGTTCGGCTGC
GGCGAGCGGTATCAGCTCACTCAAAGGCGGTAATACGGTTATCCACAGAATCAGGGGATAACGCAGGAAAGAACATGTGAGCAAAAGGCCAGCAAAAGGCCAGGAACCGTAAAA
AGGCCGCGTTGCTGGCGTTTTTCCATAGGCTCCGCCCCCCTGACGAGCATCACAAAAATCGACGCTCAAGTCAGAGGTGGCGAAACCCGACAGGACTATAAAGATACCAGGCGT
TTCCCCCTGGAAGCTCCCTCGTGCGCTCTCCTGTTCCGACCCTGCCGCTTACCGGATACCTGTCCGCCTTTCTCCCTTCGGGAAGCGTGGCGCTTTCTCATAGCTCACGCTGTA
GGTATCTCAGTTCGGTGTAGGTCGTTCGCTCCAAGCGGTAAGGAGGAGGTATGTGTAATGGGAGTGAAGTTTTGTGAATAGAGAAT
>P3_FWD
GTAATAGGGATTAAGAGGTTTGTGAAGTTTAGTGGGGTGAAATGTGGGGAATAGGCTCCGCCCCCCTGACGAGCATCACAAAAATCGACGCTCAAGTCAGAGGTGGCGAAACCC
GACAGGACTATAAAGATACCAGGCGTTTCCCCCTGGAAGCTCCCTCGTGCGCTCTCCTGTTCCGACCCTGCCGCTTACCGGATACCTGTCCGCCTTTCTCCCTTCGGGAAGCGT
GGCGCTTTCTCATAGCTCACGCTGTAGGTATCTCAGTTCGGTGTAGGTCGTTCGCTCCAAGCTGGGCTGTGTGCACGAACCCCCCGTTCAGCCCGACCGCTGCGCCTTATCCGG
TAACTATCGTCTTGAGTCCAACCCGGTAAGACACGACTTATCGCCACTGGCAGCAGCCACTGGTAACAGGATTAGCAGAGCGAGGTATGTAGGCGGTGCTACAGAGTTCTTGAA
GTGGTGGCCTAACTACGGCTACACTAGAAGGACAGTATTTGGTATCTGCGCTCTGCTGAAGCCAGTTACCTTCGGAAAAAGAGTTGGTAGCTCTTGATCCGGCAAACAAACCAC
CGCTGGTAGCGGTGGTTTTTTTGTTTGCAAGCAGCAGATTACGCGCAGAAAAAAAGGATCTCAAGAAGATCCTTTGATCTTTTCTACGGGGTCTGACGCTCAGTGGAACGAAAA
```

```
CTCACGTTAAGGGATTTTGGTCATGAGATTATCAAAAAGGATCTTCACCTAGATCCTTTTAAATTAAAAATGAAGTTTTAAATCAATCTAAAGTATATATGAGTAAACTTGGTC
TGACAGTTACCAATGCTTAATCAGTGAGGCACCTATCTCAGCGATCTGTCTATTTCGTTCATCCATAGTTGCCTGACTCCCCGTCGTGTAGATAACTACGATACGGGAGGGCTT
ACCATCTGGCCCCAGTGCTGCAATGATACCGCGAGACCCACGCTCACCGGCTCCAGATTTATCAGCAATAAACCAGCCAGCCGGAAGGGCCGAGCGCAGAAGTGGTCCTGCAAC
TTTATCCGCCTCCATCCAGTCTATTAATTGTTGCCGGGAAGCTAGAGTAAGTAGTTCGCCAGTTAATAGTTTGCGCAACGTTGTTGCCATTGCTGCAGGCATCGTGGTGTCACG
CTCGTCGTTTGGTATGGCTTCATTCAGCTCCGGTTCCCAACGATCAAGGCGAGTTACATGATCCCCCATGTTGTGCAAAAAAGCGGTTAGCTCCTTCGGTCCTCCGATCGTTGT
CAGAAGTAAGTTGGCCGCAGTGTTATCACTCATGGTAATAGAGGATTTAGGAAATGTGTGAGTAAAGGGTGGATGAGTTAGGAGAT
>P4_FWD
GTAGAATGGATGGGAAATGGGAGTTGTTATAGGGGATGTTTCCAGTCTATTAATTGTTGCCGGGAAGCTAGAGTAAGTAGTTCGCCAGTTAATAGTTTGCGCAACGTTGTTGCC
ATTGCTGCAGGCATCGTGGTGTCACGCTCGTCGTTTGGTATGGCTTCATTCAGCTCCGGTTCCCAACGATCAAGGCGAGTTACATGATCCCCCATGTTGTGCAAAAAAGCGGTT
AGCTCCTTCGGTCCTCCGATCGTTGTCAGAAGTAAGTTGGCCGCAGTGTTATCACTCATGGTTATGGCAGCACTGCATAATTCTCTTACTGTCATGCCATCCGTAAGATGCTTT
TCTGTGACTGGTGAGTACTCAACCAAGTCATTCTGAGAATAGTGTATGCGGCGACCGAGTTGCTCTTGCCCGGCGTCAATACGGGATAATACCGCGCCACATAGCAGAACTTTA
AAAGTGCTCATCATTGGAAAACGTTCTTCGGGGCGAAAACTCTCAAGGATCTTACCGCTGTTGAGATCCAGTTCGATGTAACCCACTCGTGCACCCAACTGATCTTCAGCATCT
TTTACTTTCACCAGCGTTTCTGGGTGAGCAAAAACAGGAAGGCAAAATGCCGCAAAAAAGGGAATAAGGGCGACACGGAAATGTTGAATACTCATACTCTTCCTTTTTCAATAT
TATTGAAGCATTTATCAGGGTTATTGTCTCATGAGCGGATACATATTTGAATGTATTTAGAAAAATAAACAAATAGGGGTTCCGCGCACATTTCCCCGAAAAGTGCCACCTGAA
ATTGTAAACGTTAATATTTTGTTAAAATTCGCGTTAAATTTTTGTTAAATCAGCTCATTTTTTAACCAATAGGCCGAAATCGGCAAAATCCCTTATAAATCAAAAGAATAGACC
GAGATAGGGTTGAGTGTTGTTCCAGTTTGGAACAAGAGTCCACTATTAAAGAACGTGGACTCCAACGTCAAAGGGCGAAAAACCGTCTATCAGGGCGATGGCCCACTACGTGAA
CCATCACCCTAATCAAGTTTTTTGGGGTCGAGGTGCCGTAAAGCACTAAATCGGAACCCTAAAGGGAGCCCCCGATTTAGAGCTTGACGGGGAAAGCCGGCGAACGTGGCGAGA
AAGGAAGGGAAGAAAGCGAAAGGAGCGGGCGCTAGGGCGCTGGCAAGTGTAGCGGTCACGCTGCGCGTAACCACCACACCCGCCGCGCTTAATGCGCCGCTACAGGGCGCGTCC
CATTCGCCAATCCGGATATAGTTCCTCCTTTCAGCAAAAAACCCCTGGAGGTGGAGAATGGATGGAGAAGTGTGGGAATAGATGTA
>P5_FWD
GAAGTGTATATTTGGGGTGGGAGGTGGAGAATGGATGGAGAAGTGTGGGAGGGGTCGAGGTGCCGTAAAGCACTAAATCGGAACCCTAAAGGGAGCCCCCGATTTAGAGCTTGA
CGGGGAAAGCCGGCGAACGTGGCGAGAAAGGAAGGGAAGAAAGCGAAAGGAGCGGGCGCTAGGGCGCTGGCAAGTGTAGCGGTCACGCTGCGCGTAACCACCACACCCGCCGCG
CTTAATGCGCCGCTACAGGGCGCGTCCCATTCGCCAATCCGGATATAGTTCCTCCTTTCAGCAAAAAACCCCTCAAGACCCGTTTAGAGGCCCCAAGGGGTTATGCTAGTTATT
GCTCAGCGGTGGCAGCAGCCAACTCAGCTTCCTTTCGGGCTTTGTTAGCAGCCGGATCTCAGTGGTGGTGGTGGTGGTGCTCGAGTGCGGCCGCAAGCTTGTCGACGGAGCTCG
AATTcTCATTTTTCGAACTGCGGGTGGCTCCAAGCAGAATCGCGCTGCGCTTCATCTTTTTTCAGATATTCATCCAGCGCGCCTTTTTTAATCAGATAATCTTCAAAGCTCGGC
GGCAGCGCCATGCTATCATCCCAATCCGGCAGCACGCTAATATCATCGCTATAGCTCAGAAAGTTGCCGTTCGCGCGGCGATCCAGCTGCTGCGCGCGCATAAAGTTATATTTG
CGATGGCTAATTTTGTTCTGACTCAGGCCATCGCGAATAATGGTCGGGCGAATAAACACCATCAGGTTGCGTTTGCGTTTGGTGTTGCTCGTGCTTTTAAACAGGTTGCCAATA
ATCGGAATATCGCCCAGCAGCGGCACTTTGCTCACGCTTTCTTGAATATCTTCATCAATCAGGCCGCCCAGCACAATGGTGCCGCCATCTTCCGCCATCACGGTGGTTTTAATT
TCGCGTTTGTTAAAGCTCACATCAATGCCGGTCGCGCCCGCCACGCTGCTCACTTCTTGTTCAATGGTCAGCATCACCGCATCGCCTTCGTTAATCTGCGGGGTCACTTTCAGT
TTAATGCCCACTTCTTTGCGTTCCACGGTCTGAAACGGATTCGTGTTGCTACTGCTCGCGGTCGCGCCGGTAATGGTCGGCACTTCTTGGCCCACAATAAAGCTCGCTTCTTGG
TTATCCAGGGTGGTCAGGCTCGGGGTCGCCAGAATGTTGCTGTTGGTCGCGGTGGTCACCGCTTGCAGCACCGCGCCCCAGCCATCTTTAATCACGCCCACCATCGTGCCGTTA
ATGCCGCCCAGCAGGCTCGCCAGGGTGCCAAAATCGAGTTGAGAGGAGGGGTAGGGTGGTGTTGGAGGTAATAGGGATTAAGAGGT
>P6_FWD
ACCGCGTGATTGCGAGCATTACCGAAGGCAAAGAAATTCAAGGCGATTTTGGCACCCTGGCGAGCCTGCTGGGCGGCATTAACGGCACGATGGTGGGCGTGATTAAAGATGGCT
GGGGCGCGGTGCTGCAAGCGGTGACCACCGCGACCAACAGCAACATTCTGGCGACCCCGAGCCTGACCACCCTGGATAACCAAGAAGCGAGCTTTATTGTGGGCCAAGAAGTGC
CGACCATTACCGGCGCGACCGCGAGCAGTAGCAACACGAATCCGTTTCAGACCGTGGAACGCAAAGAAGTGGGCATTAAACTGAAAGTGACCCCGCAGATTAACGAAGGCGATG
```

CGGTGATGCTGACCATTGAACAAGAAGTGAGCAGCGTGGCGGGCGCGACCGGCATTGATGTGAGCTTTAACAAACGCGAAATTAAAACCACCGTGATGGCGGAAGATGGCGGCA
CCATTGTGCTGGGCGGCCTGATTGATGAAGATATTCAAGAAAGCGTGAGCAAAGTGCCGCTGCTGGGCGATATTCCGATTATTGGCAACCTGTTTAAAAGCACGAGCAACACCA
AACGCAAACGCAACCTGATGGTGTTTATTCGCCCGACCATTATTCGCGATGGCCTGAGTCAGAACAAAATTAGCCATCGCAAATATAACTTTATGCGCGCGCAGCAGCTGGATC
GCCGCGCGAACGGCAACTTTCTGAGCTATAGCGATGATATTAGCGTGCTGCCGGATTGGGATGATAGCATGGCGCTGCCGCCGAGCTTTGAAGATTATCTGATTAAAAAGGCG
CGCTGGATGAATATCTGAAAAAAGATGAAGCGCAGCGCGATTCTGCTTGGAGCCACCCGCAGTTCGAAAAATGAgAATTCGAGCTCCGTCGACAAGCTTGCGGCCGCACTCGAG
CACCACCACCACCACCACTGAGATCCGGCTGCTAACAAAGCCCGAAAGGAAGCTGAGTTGGCTGCTGCCACCGCTGAGCAATAACTAGCATAACCCCTTGGGGCCTCTAAACGG
GTCTTGAGGGGTTTTTTGCTGAAAGGAGGAACTATATCCGGATTGGCGAATGGGACGCGCCCTGTAGCGGCGCATTAAGCGCGGCGGGTGTGGTGGTTACGCGCAGCGTGACCG
CTACACTTGCCAGCGCCCTAGCGCCCGCTCCTTTCGCTTTCTTCCCTTCCTTTCTCGCCA
>P7_FWD
CATAACCCCTTGGGGCCTCTAAACGGGTCTTGAGGGGTTTTTTGCTGAAAGGAGGAACTATATCCGGATTGGCGAATGGGACGCGCCCTGTAGCGGCGCATTAAGCGCGGCGGG
TGTGGTGGTTACGCGCAGCGTGACCGCTACACTTGCCAGCGCCCTAGCGCCCGCTCCTTTCGCTTTCTTCCCTTCCTTTCTCGCCACGTTCGCCGGCTTTCCCCGTCAAGCTCT
AAATCGGGGGCTCCCTTTAGGGTTCCGATTTAGTGCTTTACGGCACCTCGACCCCAAAAAACTTGATTAGGGTGATGGTTCACGTAGTGGGCCATCGCCCTGATAGACGGTTTT
TCGCCCTTTGACGTTGGAGTCCACGTTCTTTAATAGTGGACTCTTGTTCCAAACTGGAACAACACTCAACCCTATCTCGGTCTATTCTTTTGATTTATAAGGGATTTTGCCGAT
TTCGGCCTATTGGTTAAAAAATGAGCTGATTTAACAAAAATTTAACGCGAATTTTAACAAAATATTAACGTTTACAATTTCAGGTGGCACTTTTCGGGGAAATGTGCGCGGAAC
CCCTATTTGTTTATTTTTCTAAATACATTCAAATATGTATCCGCTCATGAGACAATAACCCTGATAAATGCTTCAATAATATTGAAAAAGGAAGAGTATGAGTATTCAACATTT
CCGTGTCGCCCTTATTCCCTTTTTTGCGGCATTTTGCCTTCCTGTTTTTGCTCACCCAGAAACGCTGGTGAAAGTAAAAGATGCTGAAGATCAGTTGGGTGCACGAGTGGGTTA
CATCGAACTGGATCTCAACAGCGGTAAGATCCTTGAGAGTTTTCGCCCCGAAGAACGTTTTCCAATGATGAGCACTTTTAAAGTTCTGCTATGTGGCGCGGTATTATCCCGTAT
TGACGCCGGGCAAGAGCAACTCGGTCGCCGCATACACTATTCTCAGAATGACTTGGTTGAGTACTCACCAGTCACAGAAAAGCATCTTACGGATGGCATGACAGTAAGAGAATT
ATGCAGTGCTGCCATAACCATGAGTGATAACACTGCGGCCAACTTACTTCTGACAACGATCGGAGGACCGAAGGAGCTAACCGCTTTTTTGCACAACATGGGGGATCATGTAAC
TCGCCTTGATCGTTGGGAACCGGAGCTGAATGAAGCCATACCAAACGACGAGCGTGACAC
>P8_FWD
ACGGATGGCATGACAGTAAGAGAATTATGCAGTGCTGCCATAACCATGAGTGATAACACTGCGGCCAACTTACTTCTGACAACGATCGGAGGACCGAAGGAGCTAACCGCTTTT
TTGCACAACATGGGGGATCATGTAACTCGCCTTGATCGTTGGGAACCGGAGCTGAATGAAGCCATACCAAACGACGAGCGTGACACCACGATGCCTGCAGCAATGGCAACAACG
TTGCGCAAACTATTAACTGGCGAACTACTTACTCTAGCTTCCCGGCAACAATTAATAGACTGGATGGAGGCGGATAAAGTTGCAGGACCACTTCTGCGCTCGGCCCTTCCGGCT
GGCTGGTTTATTGCTGATAAATCTGGAGCCGGTGAGCGTGGGTCTCGCGGTATCATTGCAGCACTGGGGCCAGATGGTAAGCCCTCCCGTATCGTAGTTATCTACACGACGGGG
AGTCAGGCAACTATGGATGAACGAAATAGACAGATCGCTGAGATAGGTGCCTCACTGATTAAGCATTGGTAACTGTCAGACCAAGTTTACTCATATATACTTTAGATTGATTTA
AAACTTCATTTTTAATTTAAAAGGATCTAGGTGAAGATCCTTTTTGATAATCTCATGACCAAAATCCCTTAACGTGAGTTTTCGTTCCACTGAGCGTCAGACCCCGTAGAAAAG
ATCAAAGGATCTTCTTGAGATCCTTTTTTTCTGCGCGTAATCTGCTGCTTGCAAACAAAAAAACCACCGCTACCAGCGGTGGTTTGTTTGCCGGATCAAGAGCTACCAACTCTT
TTTCCGAAGGTAACTGGCTTCAGCAGAGCGCAGATACCAAATACTGTCCTTCTAGTGTAGCCGTAGTTAGGCCACCACTTCAAGAACTCTGTAGCACCGCCTACATACCTCGCT
CTGCTAATCCTGTTACCAGTGGCTGCTGCCAGTGGCGATAAGTCGTGTCTTACCGGGTTGGACTCAAGACGATAGTTACCGGATAAGGCGCAGCGGTCGGGCTGAACGGGGGGT
TCGTGCACACAGCCCAGCTTGGAGCGAACGACCTACACCGAACTGAGATACCTACAGCGTGAGCTATGAGAAAGCGCCACGCTTCCCGAAGGGAGAAAGGCGGACAGGTATCCG
GTAAGCGGCAGGGTCGGAACAGGAGAGCGCACGAGGGAGCTTCCAGGGGGAAACGCCTGGTATCTTTATAGTCCTGTCGGGTTTCGCCAC
>P9_FWD
CTCAAGACGATAGTTACCGGATAAGGCGCAGCGGTCGGGCTGAACGGGGGGTTCGTGCACACAGCCCAGCTTGGAGCGAACGACCTACACCGAACTGAGATACCTACAGCGTGA
GCTATGAGAAAGCGCCACGCTTCCCGAAGGGAGAAAGGCGGACAGGTATCCGGTAAGCGGCAGGGTCGGAACAGGAGAGCGCACGAGGGAGCTTCCAGGGGGAAACGCCTGGTA
TCTTTATAGTCCTGTCGGGTTTCGCCACCTCTGACTTGAGCGTCGATTTTTGTGATGCTCGTCAGGGGGGCGGAGCCTATGGAAAAACGCCAGCAACGCGGCCTTTTTACGGTT

```
CCTGGCCTTTTGCTGGCCTTTTGCTCACATGTTCTTTCCTGCGTTATCCCCTGATTCTGTGGATAACCGTATTACCGCCTTTGAGTGAGCTGATACCGCTCGCCGCAGCCGAAC
GACCGAGCGCAGCGAGTCAGTGAGCGAGGAAGCGGAAGAGCGCCTGATGCGGTATTTTCTCCTTACGCATCTGTGCGGTATTTCACACCGCATATATGGTGCACTCTCAGTACA
ATCTGCTCTGATGCCGCATAGTTAAGCCAGTATACACTCCGCTATCGCTACGTGACTGGGTCATGGCTGCGCCCCGACACCCGCCAACACCCGCTGACGCGCCCTGACGGGCTT
GTCTGCTCCCGGCATCCGCTTACAGACAAGCTGTGACCGTCTCCGGGAGCTGCATGTGTCAGAGGTTTTCACCGTCATCACCGAAACGCGCGAGGCAGCTGCGGTAAAGCTCAT
CAGCGTGGTCGTGAAGCGATTCACAGATGTCTGCCTGTTCATCCGCGTCCAGCTCGTTGAGTTTCTCCAGAAGCGTTAATGTCTGGCTTCTGATAAAGCGGGCCATGTTAAGGG
CGGTTTTTTCCTGTTTGGTCACTGATGCCTCCGTGTAAGGGGGATTTCTGTTCATGGGGGTAATGATACCGATGAAACGAGAGAGGATGCTCACGATACGGGTTACTGATGATG
AACATGCCCGGTTACTGGAACGTTGTGAGGGTAAACAACTGGCGGTATGGATGCGGCGGGACCAGAGAAAAATCACTCAGGGTCAATGCCAGCGCTTCGTTAATACAGATGTAG
GTGTTCCACAGGGTAGCCAGCAGCATCCTGCGATGCAGATCCGGAACATAATGGTGCAGGGCGCTGACTTCCGCGTTTCCAGACTT
>P10_FWD
AACATGCCCGGTTACTGGAACGTTGTGAGGGTAAACAACTGGCGGTATGGATGCGGCGGGACCAGAGAAAAATCACTCAGGGTCAATGCCAGCGCTTCGTTAATACAGATGTAG
GTGTTCCACAGGGTAGCCAGCAGCATCCTGCGATGCAGATCCGGAACATAATGGTGCAGGGCGCTGACTTCCGCGTTTCCAGACTTTACGAAACACGGAAACCGAAGACCATTC
ATGTTGTTGCTCAGGTCGCAGACGTTTTGCAGCAGCAGTCGCTTCACGTTCGCTCGCGTATCGGTGATTCATTCTGCTAACCAGTAAGGCAACCCCGCCAGCCTAGCCGGGTCC
TCAACGACAGGAGCACGATCATGCGCACCCGTGGCCAGGACCCAACGCTGCCCGAGATCTCGATCCCGCGAAATTAATACGACTCACTATAGGGAGACCACAACGGTTTCCCTC
TAGAAATAATTTTGTTTAACTTTAAGAAGGAGATATACATATGAAATACCTGCTGCCGACCGCTGCTGCTGGTCTGCTGCTCCTCGCTGCCCAGCCGGCGATGGCCATGGATCA
AGAAGTGGAAATTGTGAAATTAAAATATGCGAGCGCGAGCGAAATGGTGCGCATTATTGATACCCTGACCAAAAACAGCGCGAAAGATCGCACCCCGGATAACCTGATTCCGAA
AGTGGTGGCGGATGACCGCACCAACAGCGTGATTGTGAGCGGCGAAGGCAAAGCGCGTCAGCGCACCATTGATATGATTATGCGCCTGGATAGCGAACTGGAAACCAACGGCAA
CACCAAAGTGTATTATCTGAAATATGCGAAGGCCGAAGATCTGGTGAAAGTGCTGCAAGGCGTGAGCAGTAGCATTGCGGAAGATAAAGCGGGCGGCAGCAACAAACGCAGCAG
TAGCAGTAGCGCGCGCGATGTGAGCATTGATGCGCATCCGGATAGCAACGCGCTGGTGATTACCGCGGAACCGGATATGATGCGCAGCCTGGAAGCGGTGATTCGTCAGCTGGA
TATTCGCCGCGCGCAAGTGCAAGTGGAAGCGATTATTGTGGAAGTGTTTGAAGGCGATGGCTTTAACCTGGGCGTGCAGTGGATTAGCGAAGAAGGCGGCATGGTGCAGTTTAC
CAACAGCACCCCGATTGGCGCGCTGGCGGTGGGCGCGGAACAAGCGCGCACCCGCAAAGAACCGGATGATATTACCTATCGCACCGATGCGGCGGGTCAGCCGATTTATAACGC
GAACGGCA
>P11_FWD
ATTACGGACGAGTACAAGGTGCCGAGCAAAAAATTCAAAGTTCTGGGCAATACCGGTTGATGCACGGCCGGACTTCGACTAAGCTTATGACCTCAGTCCGGCCGTCCTGAGTCT
TTTATAGGATATCGATACTATACAACCGACGTGCATCTCTACCCTTAGGTTAGAGACTAGGACGACATTGCATGCAATGTACAGTCGAAGTCCCTGGGAGGTAGTATCCTATTT
ACTCGTACATAAGAGAACGTGGGACGTCACGGCAGTGATCACAGTGTGAGAAGGGTCCGTATGTCCAGGGACTGTGCCCTACTCCTAAGGTCTACTAACAGTGAACTCCCAGCG
TAGGGGCACACGCACGTCGGACCTTTTAGCTAAGAGGCTTCTCACACGTGTGGTACCACTGTTAACTTAAGTGGGGACATGTCGTCCTAGTCAACTAATCAACCTTAGTAGGAG
TTGACATACGTTAGTCTCTATGGGTACTACCTCGGATCCAATTCCAAAGGTGGAGTAACGGACGGCGCCGTGCGTCTAACGTAAATTACAAATCGTCGCGACGTATAGAACCTA
AGGCTAAAAGACTCTTATTATACGATTTGTAATAGCCCATAGATGAGGCCTCATTAGACGCAGGTACGGTAGAGATGGGCTATTAGTTCACTGCCGTTCTCAGGACAAATTGCC
CCTAGGGCAATTTGGATGTTTGGAATTGAGCCGCGGCTCAATTGGGCCCAAATGTCAAGCCTTGACTACAAGTACGTCCGAGGTCCATTTGTCCCACGGTAGGCATAGTAATTA
CTAGTACCGGTTGTAGTCGGTACACACTGTACCCATCTATGCCTACGCCGAATTCGGCGTACGCTGGACCGTGACGCGTTACGTACCTGCGTGTGTTCGAACACAAGGTTCTAT
ACCCGGGTATAATAAAACATCCGACTGAGGTCATAAGCAAGATTCTAATGGACACTTAGCTGTCTAGACAGCTAGCCTCTTAGAATCTTGCTTAGGAATGCATTCCTACCGTAC
TTGTGTACGAGTAGACCTCCACCTTGTATAGAGCCGGCTCTAAGTGTCCCCACGGAACTTCGAGGAAGTCGTGGATAAGGGGGCCTCTGCCCAGTCCTTCATCGAAAGGATGAC
TAAC
```

### Section X FASTA Sequences of Digital Information Stored in DNA

#### Table S20 | sustech_introduction.txt

```
>index0_0
TTTCTGTTGGTGCTGATATTGCTAACAGGACCAGGCGAAGAACAGGACCAGGCG
AAGTCAGCAGACAGGATACATCTCGGCGTAGTCGGATAATACGGGCCTAACAAT
CAGACCCTCTGGAATTTGGAATACCTCTGGACGAGGCCTCGAAAAGTTACGCTT
GGGCAAGCCTCACACTGGATCTCCCGGCCTGAGGAACGAAAACTTCCAACCATG
ACTAGGGGAGGCGCACGAAGATAGAGC
>index1_0
TTTCTGTTGGTGCTGATATTGCTAACAGGACCAGGCGAAGAAAGAAACAACGGA
TGATCTGCACACAGACACTGTCGATGAGTTGAGTGTCAACGAAGGCCTGCCATG
GATTGGTCATCGTCTACGGTTTATCCCCTGCGCAGGCCTCCTATGAACTCATTA
CGTGGTATCTACAAGACATCTCACGGGCCTGTAACGGCAGTCTCTTTTGATCAT
GGGTGTCCAAATTGGCGAAGATAGAGC
>index2_0
TTTCTGTTGGTGCTGATATTGCTAACAGGACCAGGCGAAGAAAGTTTCCACCCA
CTCTCCCTTACTGTTCTGTGCTCACACGTTAGCCAAAATGGCTGGCCTTGAGAC
GAGCCAACGGAGCCAATGGAAATCCACCCTTGCCGGCCTTGCTAGTCAGGGGAG
ACGGTTTGGACGGCAACCCTCACTAGGCCTGTCTAACCCAATACCGTGAGCGAT
ACCGGATAGAGGCTAAGAAGATAGAGC
>index3_0
TTTCTGTTGGTGCTGATATTGCTAACAGGACCAGGCGAAGAATGGAGGAAAGAA
GTCTCTGCATCTCTCGATTCAGCATACGACGATGTGCCCGGAGGGCCTCAATCG
CATCAGTGGTAGCACGCTTATTACCGCTGCGGTTGGCCTTCCCTAAGTACGGGA
CTTACCAATCCCTAGTAAGTCGCGGGGCCTCCCGCAGTGACGACAGTAGGCAGA
CATGGTCATAGTGCCGGAAGATAGAGC
>index4_0
TTTCTGTTGGTGCTGATATTGCTAACAGGACCAGGCGAAGAAAGAAAGGAACAT
ACCTCCACAAGCATGTAGCCCTTGGTTCTGCAGATAGTTTGAGGGCCTTACAAG
TGGCCCGGTTTACACCAATTGTAATGCAAGACTCGGCCTACGTTCTCGTTACGG
```

```
TGCTTAGGGACGCGGTCGCTGCCACGGCCTGACGCACTGGGGACAATGGCTTGT
CTATCTGGCGTATGGCGAAGATAGAGC
>index5_0
TTTCTGTTGGTGCTGATATTGCTCCGGCCTGAACTAGTTCCCGGCCTGAACTAG
TTCTCACCTTGCTAGGGAACGAAGTGTTCTCGCCGACACAGTGGGCCTTACGAT
AGCCGATCTGGCTTACCGTGACTGTACGCGATGAGGCCTGGATTACCAGCTCGG
ATAAAAGGAGGCGTATAACCATTACGGCCTGGGTAGACGGAATAGTGACTGATA
CTCTAATCCAACCTATGAAGATAGAGC
>index6_0
TTTCTGTTGGTGCTGATATTGCTCCGGCCTGAACTAGTTCAGTCGCTCAGGGAA
GATTCAGTCAGGTACATCGGCATGCATGCGATGGCTGAACCAAGGCCTGGTCTA
TTTCGCGATGAAGCGATAGCCAAATTGCCAGTACGGCCTCCTAGAATTGGAGTG
AGAGCAAAACCCTGTAACATGTGCGGGCCTGGAGGCGTATGATTGCCTGCAATG
CCGTTATATGGCCCTTGAAGATAGAGC
>index7_0
TTTCTGTTGGTGCTGATATTGCTCCGGCCTGAACTAGTTCAGGGAGGAAACAGA
TCATCCACATCCTTGCCTGAGACACATCCTCGTAACCTAAGGCGGCCTACATGT
AAATACCGGCGGTTAGGGGCGGCAACTTATTCGTGGCCTCAGCGCGGGTGCTTC
CAACCGACTCTCTGTACACCGATGGGGCCTTATGGGCTGATAAAGTACTGGAGA
CGAGAAATCGTACAATGAAGATAGAGC
>index8_0
TTTCTGTTGGTGCTGATATTGCTAAACCACCAAGTCCCACTCAATAACCACCCA
CTCTCATGGGAGTAAGTGCAACCCACGTGTTTAGATTGCGGGGGGCCTCCCTTA
TAAATATCTTGCAGTTAGCGCATATGCCAAGGGCGGCCTGTTTAATTCTCCGTT
AATCATGCGCAGACACGCGCAGATCGGCCTCTGCCATAGTGTCGAAAGCTTGAC
GCAATAAATCCAAGGCGAAGATAGAGC
>index9_0
TTTCTGTTGGTGCTGATATTGCTAAACCACCAAGTCCCACTCCTACAACAGGGA
GGATCACACTGTCCCATGATCGGGATGCTAGGAACCAGATCGGGGCCTATAGAT
```

TGAATCCCGACCCTAGTATCTGGCATCTGGTGCGGGCCTGCAAGAGTTTCCCGG
GATACGTGGCTGTTTATAAGTAAGCGGCCTTGGAGGACTGATCGAATCAGGCAT
TCCCTTAGGTAAACGTGAAGATAGAGC
>index10_0
TTTCTGTTGGTGCTGATATTGCTAAACCACCAAGTCCCACGTTTGCTTCCCACG
TTCTCATGGGATGAGCATGACCTGGATTTCGACGGCTCCTTAGGGCCTTAACCC
ATACAAACCCAGTCAGATGTTGCGACTGCCCTACGGCCTGCCTCGCGCAGGTAC
CACATCAATCGCTTGTTTTCCTACAGGCCTTCTCGCAAAGACACAAGTGTTCGA
AGGAGGTGGTTGGTCAGAAGATAGAGC
>index11_0
TTTCTGTTGGTGCTGATATTGCTAAAGACAGACCCTCCCAAACATGTTACCATC
TTTTCATGGGATGTTGGATCACAGATGCGTTGAGGAGCCGCATGGCCTTTCAGG
CTGCTCTATTTAGAGAGTAGATTGATTGGCGCCCGGCCTTAACAGCGGCACTTA
GCCCCTAGTTAGCTCCGTTGGTGACGGCCTTATCCAACCTCGGATTACCTAGTC
CCTGCGTATAACGACCGAAGATAGAGC
>index12_0
TTTCTGTTGGTGCTGATATTGCTAAAGACAGACCCTCCCAAGGCGAAGAAACCA
GACTCTGCACTGTAAGTCTCGAGCTTGGATCGTTCTCCGGCTAGGCCTTTGAGC
GTGGACTTGAGCAAACTAAGTGCATGCCAGAGATGGCCTCTACGTCGTCTGAGG
CAGGCTCGGTGTCGTGTGGGAATTTGGCCTACCCAATCTAGTGCCCGAATGTAG
CCTTAACGACTCGCAGGAAGATAGAGC
>index13_0
TTTCTGTTGGTGCTGATATTGCTAAAGACAGACCCTCCCAACGGGTTTCCGGAC
TTTTCACCTTGCAGCAGTCAGACGTCTAGCACACTAAGTGGGCGGCCTGACAGA
GAGTAGGCTGAAGCTATCAGGAGCCAGACAAGGCGGCCTCTTGCAACCTTCAGG
CTTGGTGCTGTGAATCACGCGAACCGGCCTCTTTCCAGTTGATGATGTGCTATC
TGTTGATTAACGAGCTGAAGATAGAGC
>index14_0
TTTCTGTTGGTGCTGATATTGCTAAAGTTTCATTATGTTTCCGGCCTGAACTCA
CCCTCACCTTGTCTACTGCCACCGTGAGTGATGTAGAGGAATGGGCCTAAGGGA
GGGTAAATTGTTTCCCACACACCTGCCAAATGTAGGCCTTGCAAAGGCCCAATG
AAGGGTGGGTCAAATCCGGCAGCGTGGCCTGCGAACATGCACAAAATCGCTTAG
ACTCACCGTGCGATCAGAAGATAGAGC
>index15_0
TTTCTGTTGGTGCTGATATTGCTAAATTTCATCCGCAAAGAAACCACCAAGTCC
CACTCATGGGAGTATCACAGTCTCGAAGGACAGAATCCATCACGGCCTGTCGTT
GCGCTAAGTGCAGATCAGAGACTCGGGCTGATAAGGCCTCCGACACAACAGATA
GTCTTCTTAACTCTCCTAATGTCCCGGCCTCTTCCGCTGAGGTGAGTCATCACA
AGCCGCCTGAGTAGCTGAAGATAGAGC
>index16_0
TTTCTGTTGGTGCTGATATTGCTAAATTTCATCCGCAAAGAAAGGGATGGGTAG
GTTTCACACTGTCCACGATATCACGGGGTTGCAACTAAATCCTGGCCTGTGATC
GCTCCCAACTGTAATTTTCGCAGCCACGCATTAGGGCCTGCGTTCGAACGGAAG
CGCAGCAGGTTTACCTCGAAAGTTAGGCCTAGGAGTTCATCGGCTTGCACTTGT
GGATGATGTGGGTATGGAAGATAGAGC
>index17_0
TTTCTGTTGGTGCTGATATTGCTAACATGTTACCATCTTTAAACCACCAAGTCC
CACTCAGGTGATGCAATCAGCTCCGGTTCCATGCGTCTAAACCGGCCTCCATGT
TGGTACGCCTTACTACTTGTTACATACCGTTAAAGGCCTAGTACGCGAGTTGTT
AGCAAGCTGGTGTAGCTCATCCGGGGGCCTAAAGAGTTCCCGGATAGTATCCAG
TTATGGTCGTCCTATAGAAGATAGAGC
>index18_0
TTTCTGTTGGTGCTGATATTGCTAACATGTTACCATCTTTAGGTGCGTCAGGAA
AGATCTACCCTTCAGATGAGACATCTGGAAGTGCTAACTGGAGGGCCTAGTCCC
CGGTCTAATAGAATTCGCTTGATCGCTAGCGTTTGGCCTTCGCAGTTCGACAAA
TCGCATGGAGTTTGTTAAACGGGGCGGCCTCGCCACGTAACAATAGCGGGTCTA
GCATGAACCTTTTAAAGAAGATAGAGC
>index19_0
TTTCTGTTGGTGCTGATATTGCTAACATGTTACCATCTTTCAGGCGTTTCCGCA
AAGTCACCTTGCTAACACGTTCCTACCGCAGAGTAAGGAGTACGGCCTCTTGGG
ACACTAAAGGCTTCTAAGAATTAGGCACGACAGGGGCCTGAAATATGAGACGCG
CACACATCGCTACGTAGGAGCTTTTGGCCTCTTTGTAGCGCGTCCGGTAGTTTG
CCAGAACGCAGGGGTTGAAGATAGAGC
>index20_0
TTTCTGTTGGTGCTGATATTGCTAACATGTTACCATCTTTCCCACGTTCATCGT
CATTCAGCATGCACGAGTCTTCGAGCAGTGTTGGCAATCTGCTGGCCTCCTTAC
GGTAATAAACACGTCTGGTGTCTTATCAGGCTTCGGCCTACACTACGAGTAGGT
TATCACCGTCATGTGCGCGACACAAGGCCTTCCCATGGATGAGATCGTATCTGT
GAATATAATTCGTTTTGAAGATAGAGC
>index21_0
TTTCTGTTGGTGCTGATATTGCTAACATGTTACCATCTTTAACTCACCCACCAG
ATCTCTGCACTTCGAGCAAAAGCATCTTCGGTCTAGCATGGACGGCCTTTACGA

```
GTTACACCTCACCACACCCGCTGCGAGAATTCATGGCCTAGGTTCAATCCATAT
ATGCTACATCATATCCATTGACACAGGCCTTCAGCAGGACGATTACATGAAGCC
CTATACCGAGTAAAGGGAAGATAGAGC
>index22_0
TTTCTGTTGGTGCTGATATTGCTAACATGTTACCATCTTTAAGTCGGAAAGGGC
GTCTCAGCTGAAGTGCTGACTTGCATCTTTGCCTTCCGTACGAGGCCTCCATCT
TAGACCCTTAACGGTTTAGGTGGCACACATTATGGGCCTATTTGCTCGATCCTA
CTCCCTCACGCACGGATCCAAGTTCGGCCTTTGTTCCATCGACACTTTGTCAGA
ATGCACCCGTGTCATTGAAGATAGAGC
>index23_0
TTTCTGTTGGTGCTGATATTGCTAACATGTTACCATCTTTAGGGAGGAAAGTCC
CACTCTGCACTGTTCATGAGCGCGTATCGGTACGAGCTGATACGGCCTTCTAGT
TAGAGATCATCGCCTTTCACTTCCTTGGTTCGTCGGCCTCAGGCCGTATAGTTG
CTTCGGACCGCATCACTAGGTGATTGGCCTTCCACGGCATCATGAGACGTCCAG
TTATAGTCTTGTAATCGAAGATAGAGC
>index24_0
TTTCTGTTGGTGCTGATATTGCTAACTCGTTAATCATGTCAGTCGCTCACTCCA
TCGTCGCTACTGATGTCTCGCTGCAATGTGAAGGATCAGCACCGGCCTCCTACC
ATCCCGAACATTGTTGACCCTGTTAGGGAGATTGGGCCTCTTTTCGGACGATTC
AGCGGTGCGTCCGCATAACCTAGCAGGCCTGTATCGCTTTCCTGGCGAACCTGT
TTATCGCCACCTGGATGAAGATAGAGC
>index25_0
TTTCTGTTGGTGCTGATATTGCTAAGTAGTCAATCTGTTTACCACCCATCCTAC
AACTCTACCCTGTTACGGTCCTATTACCGTAGGCGTACTGGCAGGCCTTTACCG
ATATAGACCGGCGTGCTCACGCCTTGGCATGTGCGGCCTCAGACTGTGACTTAT
TCGCTAGAAGGGCAAGAGTGCGGGCGGCCTTTAACCTTTTGGGCCCGGACGTCA
CACTTATGGTGTTGCAGAAGATAGAGC
>index26_0
TTTCTGTTGGTGCTGATATTGCTAAGTAGTCAATCTGTTTACCGTTCTTAGGGA
TCCTCAGCTTGGTCGACAAGTTAAGCGACGATCATATGCATCGGGCCTGCCATC
TTCAATGCTGGCTAGGGCTGTGGCGACTTTTGCAGGCCTTGAAAATCATAGAGC
AAGGCACCGTTAGGTCCCTATCCATGGCCTCATACATTTCGTAGCAGCGACAGC
GTCCGGTGCCTACATTGAAGATAGAGC
>index27_0
TTTCTGTTGGTGCTGATATTGCTAATACCAGACCGATCTATGGCCGTCTTCGAT
GTCTCTGCACTGTACCTAAGAGTACGCATTGTCCGTTAGGGTCGGCCTGACCGC
```

```
CAGAGACTCATGGAAACAGTTGCTGATGACCTGTGGCCTAGATGGCGTTCATCC
ACCTCAGCGTTGGTTTCACTTGATCGGCCTCTCCGCAGCCTTCAATTGCGCGGT
AGCCGGAATTATATGTGAAGATAGAGC
>index28_0
TTTCTGTTGGTGCTGATATTGCTAATACCAGACCGATCTAAGGCGAAGAAAGAA
AGGTCAGCATCCTTGGACTATGGGTGTGGGAGTCACTATGGCCGGCCTGTCATA
GCCAACAACGCCTACGAAACGCACAGCATGTCGCGGCCTCAAAACTCCTCGGCA
CTAGTAATTGCTGACGGAGGAAAAGGGCCTAGGAAAGCACGGTCATTCCGCGTA
CCTCTAGAGAGGTCTTGAAGATAGAGC
>index29_0
TTTCTGTTGGTGCTGATATTGCTAATACCAGACCGATCTAATCGGACTTACCTC
ACCTCGCAATCGATAGCAGTTGCTCTCTAGGAATGTACTCGAAGGCCTTGTGTC
TCGGGGAGACTCCAGCAACGACATACATCTGCCCGGCCTGCGTTGAGAATTTGA
ACGGTGCATCCGTGCAATGAGCTTAGGCCTGCAGCGGCGGCAGGCAAATACCGC
TAGGACACCATCTCACGAAGATAGAGC
>index30_0
TTTCTGTTGGTGCTGATATTGCTAATCTTGTCCTCAGTACAAAGACAGACCCTC
CCATCTGCACTGTTCATGAGCGCTTCCACGTTGCTACCGATTCGGCCTTAATAC
TGCCCGGTGGAGACTACGGCTTAGGAGGTCTTAGGGCCTTCCAGGCTGAACGTC
AATGAATTGCCAGAAGCGCGGTAGAGGCCTAGGGCGAGACACGCTATGTTTAAG
CCTTTCTTTGGATGGCGAAGATAGAGC
>index31_0
TTTCTGTTGGTGCTGATATTGCTAATCTTGTCCTCAGTACAACATGTTACCGGA
GGATCGCTACTGATTCCCCTTTTGAAGGCGGAGTTGTGGCATAGGCCTACCAAA
GTCTGATCAAGAAATCCTCAACTGGCCCTGTATCGGCCTAGATGGGACGCCGAG
AGATGGGTATAAAGGTTACTTACAGGGCCTTAGCGGGTAACATCAACCTTTACG
TTCTTAACGACTGTAAGAAGATAGAGC
>index32_0
TTTCTGTTGGTGCTGATATTGCTAATTTAACAGGCGAAAGAAAGACAGACCCTC
CCATCAGCAGAAGGGTCTTGGTAGAGGCAGAAGCCATATATATGGCCTACTCGT
TCCGTTAAGACTGAACCCATCAACGCCGGATGGGGGCCTCCCTTAACAACTTCG
AAATACTTCATTAGTATCGAACCGCGGCCTCAGGAATTAACAGCGGCAATTAGG
CTGAGAACGTGGATGCGAAGATAGAGC
>index33_0
TTTCTGTTGGTGCTGATATTGCTAATTTAACAGGCGAAAGCCGGCCTGAAAGAA
ACATCAGCTGTGTGATACTCGTTCTGGCCGTAGATTTCCTCGTGGCCTTTCTAA
```

GGCGGGGTACTCAGTAGCCAATACTATGCAACGCGGCCTCTCAAGATTGCGTAA
AGCTGTCAGCAGATGCTATCCAGCGGGCCTATCCACGGTTGGTCGCCTCGTAAC
CCGGTGTCTATCCATGGAAGATAGAGC
>index34_0
TTTCTGTTGGTGCTGATATTGCTAATTTAACAGGCGAAAGACCTCACCATCGGT
TGTTCGTTGTCGTTTCGTGGACCGAGTGACTTCTTAACGCGCCGGCCTATGAGG
CAATCCGCATCCAACTATGGCACTGTTCTCATATGGCCTCCATCCCCTTAATCG
CCATCCTCTTACTTACTTATGTAATGGCCTGCGCGTCAGTCTCCCCGCCCATTC
CAACTCGTAGTCGGTGGAAGATAGAGC
>index35_0
TTTCTGTTGGTGCTGATATTGCTACCACCCATCCTACAACAATCTTGTCCTCAG
TACTCATGGGATGTTGACCGTGGAATCGCCAACCAGGTACGTAGGCCTTAATCA
AGTTATGTCCCGCGATCCGAGAAGTAAATAACAGGGCCTGTAAAGCTGTAAGAG
TGTGGATAGTGGGAATGATCCTGTTGGCCTTCTGAGGCCACACATCAGAACTCG
GGAAAAGAGCGCTGGCGAAGATAGAGC
>index36_0
TTTCTGTTGGTGCTGATATTGCTACCACCCATCCTACAACAGTTAAGGAAATCA
AACTCAGCATGCACCTTGCAGACCTTACAGTGGGCTAAAAGCCGGCCTAAGTAG
AAAAGTCTCTGGAACTCGCCATCCATTAATTACCGGCCTTACGTGACCGGACCC
CTATTCATTTTCCCAGAGCTGCTCCGGCCTGCGATGCCGAGCTGAGGATTATGA
AGTGCTGAGCCGCACTGAAGATAGAGC
>index37_0
TTTCTGTTGGTGCTGATATTGCTACCACCCATCCTACAACAAAGAAAGGAGGAA
AGATCGTACCAACTTGAGCTTGTAGTGGGCATTTGGGGTGCGTGGCCTAGTTAC
ATAGATTGTTGTCACTTGACTGTCCCAGTCAGATGGCCTAAAGCTACGTAGTGC
GATGAACCGATTCTTTCTCCCAAAAGGCCTCAGTGGTGGTGACTCTACACGCGG
CATCAGTGTCCCGTAAGAAGATAGAGC
>index38_0
TTTCTGTTGGTGCTGATATTGCTACCGACGAAACTTTGTTACTCAATAACCCGT
AGATCAGGTGATGCACTTCCCATTTTGTCTGCATGCTAGTGTTGGCCTAGTCCC
TCCCTCACAGGGCATCGGTAGCCTTAGATATAAGGGCCTCGACTTTCGATATTG
CTAACCCATCTGTGGCTAGTCAGAGGGCCTCTTCACCCGCTTTACAAGACGCTG
AGTTGCTCACGCTGGCGAAGATAGAGC
>index39_0
TTTCTGTTGGTGCTGATATTGCTACCGACGAAACTTTGTTAATTCCGTTACTCG
GACTCAGCTGTGTGACGAAATCAACTGAGCCTTACCGCCGTCAGGCCTCGGTCA

AGGTGCAGTGAACTCCAGGATGAGATCGCCTGAAGGCCTAAGTCTCTAGATAAT
CCTTCATAGTCATGTTATGGCGGGAGGCCTCGGAGCGGTATGTAGTTCCTTCCC
AACTTTAAGGGATTGAGAAGATAGAGC
>index40_0
TTTCTGTTGGTGCTGATATTGCTACCGACGAAACTTTGTTAGGCGAAGAATCTT
TACTCTGCACTTCGTAGCCTGACTAGGCTGATGATCCCAGAAGGGCCTTTGGAT
CTTGCCCAACTATTGTGGCAGACGCGATACGTGCGGCCTGTTTCTTATTTGGTG
CATTTATATTTGTACTACGTCGCGGGGCCTCCCGCGGTGCACAAGCGACGTTGT
AAGACGCTCTGGGCGAGAAGATAGAGC
>index41_0
TTTCTGTTGGTGCTGATATTGCTACCGACGAAACTTTGTTTCCCGTTTAAGGGC
GTCTCGCTACTGTCCAGATGACAAGCGAACAGGGGAAGATCTTGGCCTCCTCCA
ACTACTGAACGTGAGATGAGACCCGATAGTGACGGGCCTGCAATACTAGAGAGT
CCGTGCCGCATCGTCTTACGTTCCGGGCCTGAGATCCATCGACACTTTGTCAGA
ACGGCTTAGATCGCGCGAAGATAGAGC
>index42_0
TTTCTGTTGGTGCTGATATTGCTACCGACGAAACTTTGTTACCACGGAAATCAA
TCATCTACCCTTGGTCACTCACGACAGTTGGGAGCTCAGAGTTGGCCTGACAAT
GAATGCATCGTTTCGACACATTTATGTAGAGCCAGGCCTCCCTACACTGATTAC
TTCTGACCGCATGGGACGCTAGTCGGGCCTTGCCGAGCTTGGTCGGGATAAACA
ATCGATCATTTTCGCTGAAGATAGAGC
>index43_0
TTTCTGTTGGTGCTGATATTGCTACCTAATCAATCAATCATCCGCCTCAATTTG
CTGTCGCTACTGTGGAACTGTTATCGATGTGCATGGTGGTAGTGGCCTTTGAAC
GATGTCGACTACGGGCTGTTTATAGGGCAAGCAGGGCCTCCTCGTTAGGTTAAT
CAGGAGGTTCTTTTGGCCGATGGGAGGCCTGAGTTAATACGCATTGTATGCACC
GTCCACATAGCAGCGTGAAGATAGAGC
>index44_0
TTTCTGTTGGTGCTGATATTGCTACTCAATAACCCGTAGAACCGACGAAACTTT
GTTTCAGCATCCAGGTAGACAACAAAGGTCATGTGTGACATCAGGCCTTCCCTA
GTGTAACTGGAACTGCCGGAATTCCCTGGTGGTTGGCCTTCAACTTCTGAATTA
AGTCCGGGGATTCGCAAGACGGCTGGGCCTTAGCATCAATTCTTATTTCTCCGA
ATTTTGGAGAATCTCTGAAGATAGAGC
>index45_0
TTTCTGTTGGTGCTGATATTGCTACTCAATAACCCGTAGAAAGTAGTCACGGAT
GACTCAGCTGTGTGACGAATCTGGGTGACACGTTGTAAGTTATGGCCTAGCAAC

```
CCTTCATTGGTTGGATTCGCTAAGCCCCGGAACCGGCCTACATTTAAAGTTTTG
CAAGAGACGCTCTCGATAGATTTGAGGCCTGGAGGGCCGGTCCAAGCTCAAAGG
GTTCACCTCCCACGGAGAAGATAGAGC
>index46_0
TTTCTGTTGGTGCTGATATTGCTACTCAATAACCCGTAGAAACCCTAAAGGGAA
GATTCACCTTGTCATCTGTCACATCCTGCGCTCGGACTTTAGCGGCCTGGAGAT
GCCAGGTAAGTAGAATACTGACGACATGAAGGCGGGCCTTATATACACGGTGCG
ATCAGAGAATAGTCTGCCGTACCTCGGCCTCCTGTGGTCGGCTTTAGGCACCAG
TCACCGGGATCTGCGTGAAGATAGAGC
>index47_0
TTTCTGTTGGTGCTGATATTGCTAGGCGAAGAAACCAGACAATCTTGTTCAGCT
GGATCAGCATCCAAGACGAGCTTGATGTCGAAGCGCAGGTGTCGGCCTGAGAAA
TACGACAGCTCCATGTACAGGAGGTAGGGCATCGGGCCTAGAAAGCGCACTTAG
TCGTACATACTTGCTTGGCAGGAAGGGCCTACCGGTCTCGCTCATCGGACATCA
CTACTCAGGGAGGTGAGAAGATAGAGC
>index48_0
TTTCTGTTGGTGCTGATATTGCTAGGCGAAGAAACCAGACTCCTACAACAGGGA
GGATCAGCATCCAAGTGTTCAGAGGCACGCAAAGACGACGAAAGGCCTACGGTA
GGTCGCTCGGTCTACTCGAACGGTATAGGAATGTGGCCTACCAGGAGCATCAAC
TCCCCGTGCTGGCGACGTGATAACAGGCCTGCTGAGGGGTAATCCTACGACCCC
GTGTAACATATCATATGAAGATAGAGC
>index49_0
TTTCTGTTGGTGCTGATATTGCTAGGCGAAGAAACCAGACTCCTACTACCCACC
TCCTCAGCTTGGTCAAGGGATTGAACGCACGACTCTGCTAAGGGGCCTGGCATC
TTATGGTTGCCGTGTTGATTTGGTACTGTCGGTAGGCCTACATCCTCACTGGTA
GCACCAGGCCAACGATCTGTGCTCGGGCCTGGGCCACACGGGATGCTCTTGCTG
CAGACAACGCAAGAGGGAAGATAGAGC
>index50_0
```

```
TTTCTGTTGGTGCTGATATTGCTAGGCGAAGAAACCAGACAAAGAAAGGACCGA
AAGTCGCTACTGAAAGCAGGGTACGAGCTCAACCTATCGACTAGGCCTTACTGG
TCACATAGAGAAATAGCTGGTTTCAGAAGAGCAAGGCCTTGTCTGGAACCGGTG
TCCTTTATCGCCGTGGAACATTAAAGGCCTGGTCGCGACGACGATACCTGATAT
GGCCATTTCCGTCGTAGAAGATAGAGC
>index51_0
TTTCTGTTGGTGCTGATATTGCTAGTCGCTCACGGATGACTGTCGACGGAAAGA
TCATCAGGTGAGACGACTGTGACAGTTGACAGTCTGCGCACACGGCCTTACGCA
CAGTACCCTTGAACGGCTTATTCTACGGCACGGTGGCCTTGCTCTAGTAAGCAG
CCTATACCCGGGCTGTCACTCTCCCGGCCTTCCAGATCGAGCATGGATGATAGA
ACAGACAGCCGTGGAGGAAGATAGAGC
>index52_0
TTTCTGTTGGTGCTGATATTGCTATCGTCATGGGTCATTTATTGCATTACCCTC
CCATCTACCCTGTCGTTAGTCGCAACGACTGATCCACTGGTCCGGCCTCAGAAC
GTGACATCCACAGAGAGTGTTCATAGAAAATTAGGGCCTCGTGCGATCCCTAGT
CCCTCACGATTTACCGTACTAGAAAGGCCTTGAGGCCTCTGTTCACACGTAACC
TCTACTTTTCGTAGGCGAAGATAGAGC
>index53_0
TTTCTGTTGGTGCTGATATTGCTATCGTCATGGGTCATTTACCCTCCCAAAGTG
GACTCACCACAGTCACACATACCCGTCTGTGTACGCCATCACTGGCCTATCGCT
CAGGTGGTGTACCCTCTCAGAAGTATTAAGAGGGGGCCTATCCTGGAAGTGATT
ACCCGCGCAGACTACGTGTCGCACAGGCCTGAGTACCATGAAGTCCCTACCCCA
CCTTGTTGAGAAGCCTGAAGATAGAGC
>index54_0
TTTCTGTTGGTGCTGATATTGCTATCGTCATGGGTCATTTAGGAAAGAACGGAT
GACTCACAGTCGTAAGGGTGTCAAAAGGGGCCTCAGAGTCGTAGGCCTTGAATG
CATTGTTTTGAGGATGGCTAGCTTTCGGCGGCCTGGCCTCAGATTTCATTTAAT
AGTGCAAGTCCTCACGACACCTCTCGGCCTTGACGTTTAAGGCACACCACCTCG
ACGCGGTTGGTTGTATGAAGATAGAGC
```

### Table S21 | sustech_logo.jpg

```
>index55_0
TTTCTGTTGGTGCTGATATTGCTAACAGGACCAGGCGAAGAACAGGACCAGGCG
AAGTCATGCTGCAACACTGATCTGGTTCAAGACGATCTAGACAAAGGCCGGCGT
GTACACACCGGTATCTGTTTCAACGGACGGAGATAAGGCGTCGTTGAATGCGTC
GAGCAGCCTTATGAGACGGGGCCTCAAGGCGATTTGCTATATGTAGCTTCATAT
ATCCCTCAGTGCGGCGGAAGATAGAGC
>index56_0
TTTCTGTTGGTGCTGATATTGCTAACAGGACCAGGCGAAGAAAGAAACAACGGA
TGATCATGGACAGTTCTGGAACGACGCATACTCTCTAAGGCACAAGGCCTCAGC
GATAAATCCGTGATATAGAGGTGTTAGGAGTAAAAAGGCTGAGTCGGGCCGCCC
TGCACCTTGTACCATCTTGATAACGAAGGCATTTTAAATCAAATCCGACTAGGC
TGAACAGACTCTACTAGAAGATAGAGC
>index57_0
TTTCTGTTGGTGCTGATATTGCTAACAGGACCAGGCGAAGAAAGTTTCCACCCA
CTCTCACAGCACACTCTGCACACCTGACTCTCATCGGATATCCAAGGCGGGACA
TTTTAGGCCACACGTACCAGTGCTGCGAGTCTAAAAGGCCACATGGGAACGGTA
ACTTGCCGTGAGTCAAAAGGGCTATAAGGCTGCCTCAATCAAATACGACTAGGC
GCAGCAAAACGCCAGCGAAGATAGAGC
>index58_0
TTTCTGTTGGTGCTGATATTGCTAACAGGACCAGGCGAAGAATGGAGGAAAGAA
GTCTCACAGCACACTCTGCACACCTGACTCTCATCGGATATCCAAGGCGGGACA
TTTTAGGCCACACGTACCAGTGCTGCGAGTCTAAAAGGCCACATGGGAACGGTA
ACTTGCCGTGAGTCAAAAGGGCTATAAGGCTGCCTCAATCAAATACGCCCTCTA
CCACATTTGAATGCGGGAAGATAGAGC
>index59_0
TTTCTGTTGGTGCTGATATTGCTAACAGGACCAGGCGAAGAAAGAAAGGAACAT
ACCTCAACGCTTGTCCCTGACAGCTATGACTGCCCTCAACAATAAGGCTCGAGG
TCCCTGCATCAAGCCCGTAATAGGAAGCTTTACCAAGGCGAAGGTCCCTATACA
AACGAAAGACGTATAACACCGCTCAAAGGCGATTAGGATCACCTCCTTGTTATG
GGCTAGCGATTTCCTCGAAGATAGAGC
>index60_0
TTTCTGTTGGTGCTGATATTGCTCCGGCCTGAACTAGTTCCCGGCCTGAACTAG
TTCTCAGCTCTTGTTCAGGCTTACACCGACGGAAGGCTCTCTTAAGGCGGCATT
CTGCATCTTAAACGTACCAGTGCTGCGAGTCTAAAAGGCCACATGGGAACGGTA
ACTTGCCGTGAGTCAAAAGGGCTATAAGGCTGCCTCAATCAAATACGACTAGGC
TGCTGGACGGCTACCGGAAGATAGAGC
>index61_0
TTTCTGTTGGTGCTGATATTGCTCCGGCCTGAACTAGTTCAGTCGCTCAGGGAA
GATTCACAGCACACTCTGCACACCTGACTCTCATCGGATATCCAAGGCGGGACA
TTTTAGGCCACACGTACCAGTGCTGCGAGTCTAAAAGGCCACATGGGAACGGTA
ACTTGCCGTGAGTCAAAAGGGCTATAAGGCTGCCTCAATCAAATACGACTAGGC
GCAGCAAAACGCCAGCGAAGATAGAGC
>index62_0
TTTCTGTTGGTGCTGATATTGCTCCGGCCTGAACTAGTTCAGGGAGGAAACAGA
TCATCACAGCACACTCTGCACACCTGACTCTCATCGGATATCCAAGGCGGGACA
TTTTAGGCCACACGTACCAGTGCTGCGAGTCTAAAAGGCCACGATCTCGACCCG
GATCAGCATATGTTTTCTATCCAATAAGGCATTGTATGATAGCCGCCGGCCGCC
CGCAGTGATGGACCCGGAAGATAGAGC
>index63_0
TTTCTGTTGGTGCTGATATTGCTAAACCACCAAGTCCCACTCAATAACCACCCA
CTCTCGCTTAGCTATCTGTCTGAACCCTGAGTGCGCCAGACTAAAGGCCCTCTC
CACAGTGATTCATCGTCGAGCACTAGTCAATCATAAGGCCAGGTGTGGATCGCA
ATTGGTAGCAGACAAGGCCCCTCGCAAGGCGCTAAAGGTACAGTCGTCCGGACG
TTAGTCTATAGACGATGAAGATAGAGC
>index64_0
TTTCTGTTGGTGCTGATATTGCTAAACCACCAAGTCCCACTCCTACAACAGGGA
GGATCGTAGGAAGCGTCAATCATTCAGGTCAAGATGCTCTAGAAAGGCTCCAAG
AGGACCTGTCACGTGACGCTAATACGCCCCACGCAAGGCTAAGACAAAAGAGCG
CTTCGTTATTTCTAGAGGAGGGAGAAAGGCTGGACGTTCAAACCGCTACCCTCC
GGAAGGTCGGATCTTAGAAGATAGAGC
>index65_0
TTTCTGTTGGTGCTGATATTGCTAAACCACCAAGTCCCACGTTTGCTTCCCACG
TTCTCGTTGACCTGTTCAGGGAAAGTTACGGCTAAGAACCAATAAGGCGTACTG
ACGCCCTGCGAAACACTTATATCTGCAGTACTTAAAGGCTGCCGGACCTCGTAG
CTATCACCGCCTCACGACACCTCTCAAGGCTGACGTTTAAGGCACACCACCTCG
GGTGACAGCGTATTCGGAAGATAGAGC
```

>index66_0
TTTCTGTTGGTGCTGATATTGCTAAAGACAGACCCTCCCAAACATGTTACCATC
TTTTCGCTTAGCACAGCTTGGTTTTACCCGCTACCACGTATGTAAGGCGATCGG
AGGTACGCCTTCAGGGAAACGGCAGACGTTTCCTAAGGCCGACCTGTGATAAGA
TGTGCCACCAACCAAGCCCCGTCTCAAGGCCTGTACGACACTCCTAGGTCCTCA
GTGTTTTGGTGCTATTGAAGATAGAGC
>index67_0
TTTCTGTTGGTGCTGATATTGCTAAAGACAGACCCTCCCAAGGCGAAGAAACCA
GACTCGCTTAGCACTGGTAGCAGTAACTGTGGCTAGGCCACTAAAGGCGATGTT
CGTACGGCTCCCTCTGAGGAACGTAGGCAGTAGAAAGGCTAATAGACCTCAGGC
AACCAATCACAATTGAAAGGGCCCAAAGGCTTAGTACTCGTACCATCCTTGCCG
GGGACTCAATCAGGCGGAAGATAGAGC
>index68_0
TTTCTGTTGGTGCTGATATTGCTAAAGACAGACCCTCCCAACGGGTTTCCGGAC
TTTTCAGCTCAGAGACAAGGCTCGTACCTCTCTTGCCTACACCAAGGCTCCAGT
TCCTGAGACGTCAAATAATTCATAACTTAGATGAAAGGCTGCGTCTAGATCAAG
AGAGGGCAGGCTAACTTGTGGCGGCAAGGCGAGTTGCGTGGCCTCGCGGCTATA
AGAGAAAATCAATATGGAAGATAGAGC
>index69_0
TTTCTGTTGGTGCTGATATTGCTAAAGTTTCATTATGTTTCCGGCCTGAACTCA
CCCTCTACCTCAGGAATCCCCATGATTTGGTTGTCCAGCTACAAAGGCCTACGT
CGGCCCCAATCTGCCACTCGCTCTCGATTGTAACAAGGCATCCTGGTAACACCC
CGTGGTAATGATGTCAGTGTTGGAAAAGGCAAGTCCTCCCGTGGGCTGCACAAA
CGAGTGAGTTTTAGACGAAGATAGAGC
>index70_0
TTTCTGTTGGTGCTGATATTGCTAAATTTCATCCGCAAAGAAACCACCAAGTCC
CACTCTACCTCTGCACCTGTAGAGCCTAGTGCAAACCCTCGATAAGGCTCGCCA
ATTTACTGCTACTCGCCCACAAGAGGAACAACGCAAGGCGTTACAGGTGGTAGC
TTAGCTTGTTGAGTCGAGGATCACCAAGGCGGTCGTTCATAGGCTAGCTTCCGA
CCAACCGACCATCCTGGAAGATAGAGC
>index71_0
TTTCTGTTGGTGCTGATATTGCTAAATTTCATCCGCAAAGAAAGGGATGGGTAG
GTTTCTACCACCAAGAACCTCCGAGTCATGCCTAAAGACAACGAAGGCACAAAC
TACTAACGAACGTGCCGAAAGATGTCGCGGTCACAAGGCAGCCGGTGGTCTTGT
GTCTCTTAAGCTTAGTGTATGGTAGAAGGCGCTGACATGGGCCCCTGCTGTTAC
TTTTACGGTGTTGCGCGAAGATAGAGC
>index72_0
TTTCTGTTGGTGCTGATATTGCTAACATGTTACCATCTTTAAACCACCAAGTCC
CACTCAGCTCATCTCCTGAATGTGCGCACTACCGGTACCTGTAAAGGCACATCC
TACCCCACGACAGGCTGTTCCAAGCTTACGCAGGAAGGCAGAATATCCCCAGGC
AGACTAATGTCTCCGTCACCGATTAAAGGCGGCAATAAAACTTGAGTGCCGAGC
TCATCCCCGATCACCTGAAGATAGAGC
>index73_0
TTTCTGTTGGTGCTGATATTGCTAACATGTTACCATCTTTAGGTGCGTCAGGAA
AGATCAGTCAGGTATGGGGTGTTAATCCCATAGGTAACCCGCCAAGGCCACACA
TCGTTGTCTCTCCAGCCCACGTACATGATGTACCAAGGCTTGTTCATGACTTGG
ACCTGAAGCCCACAAAAGCGGGTGGAAGGCTATAAAGGTGTCCAGTCACTGCAT
GGAGCTCCTCGCGTCTGAAGATAGAGC
>index74_0
TTTCTGTTGGTGCTGATATTGCTAACATGTTACCATCTTTCAGGCGTTTCCGCA
AAGTCGCTTAGCACCTATCAACAGTGCGCATTGCGCATCGCAGAAGGCACCAGT
GATCTGACTAGCTTACCCTGGAATCCTCACACGTAAGGCGATTTACTATTTCTC
GGCGTGAGTCCTCACGACACCTCTCAAGGCTGACGTTTAAGGCACACCACCTCG
GGGCACTCCAAGTTATGAAGATAGAGC
>index75_0
TTTCTGTTGGTGCTGATATTGCTAACATGTTACCATCTTTCCCACGTTCATCGT
CATTCGCTTAGCACTGGTAGCAGTAACTGTGGCTAGGCCACTAAAGGCGATGTT
CGTACGGCTCCCTTGGAGGTCTGAACTAACGTGTAAGGCCATATGACGGGGCAA
GGCGTGAGTCCTCACGACACCTCTCAAGGCTGACGTTTAAGGCACACCACCTCG
ACGTGATTGGTCACTCGAAGATAGAGC
>index76_0
TTTCTGTTGGTGCTGATATTGCTAACATGTTACCATCTTTAACTCACCCACCAG
ATCTCGCTTAGCACTGGTAGCAGTAACTGTGGCTAGGCCACTAAAGGCGATGTT
CGTACGGCTCCCAGTAACAAGACATCCTCAATTGAAGGCATTGCTACCACGCTG
AGTGTGAGGACATGCGTTTACGGGTAAGGCCTTTAGGATGATATCGCAAGTTGC
AAGTTGATCTCGCTGAGAAGATAGAGC
>index77_0
TTTCTGTTGGTGCTGATATTGCTAACATGTTACCATCTTTAAGTCGGAAAGGGC
GTCTCACCATGCACGTTGTCTGACACGGATCTCACGCACTCTCAAGGCCTATGT
TCAGACTAATCGAAGCGAATAGTTTTCGGTGCCGAAGGCAGGAAAAGAATGTTG
CCAGTGCCGTTAGCGGCGAGCTGATAAGGCCACTGTCATATCCAGGGTAGAGAT
ACGCACTGTTGCCACGGAAGATAGAGC

>index78_0
TTTCTGTTGGTGCTGATATTGCTAACATGTTACCATCTTTAGGGAGGAAAGTCC
CACTCATGGGAACAAGTCAGACTCGAAGCGCCTCCTACTAATAAAGGCTGGAGT
TGTGCGTGGGAGCGACATGCACCGGTTCCGTACCAAGGCAGACCTGTAAGTTTA
CACCTAGCAAACAGACACTGCCTTCAAGGCTAGGAAAATTGGTGCCCCTTCCCT
GGGCAATTAATTCGGAGAAGATAGAGC
>index79_0
TTTCTGTTGGTGCTGATATTGCTAACTCGTTAATCATGTCAGTCGCTCACTCCA
TCGTCCAACCTCTCCTGTTGTCCCTACGTTTGTTTCCACCCTAAAGGCCGCATG
GACAGTCTGGGGATCAGCATTGTCGCGGTGTGTTAAGGCCCCTGACAACTGGTT
CGTTTATGTAATGATGTCGGACACAAAGGCACGCACGAGAAAGGATAATTCAAA
TGTATTTAATGACGTCGAAGATAGAGC
>index80_0
TTTCTGTTGGTGCTGATATTGCTAAGTAGTCAATCTGTTTACCACCCATCCTAC
AACTCAGCTAGGTTGCTCTGGTTGTTAACAGTCTAGACAGAGTAAGGCCATGGA
CGACGGATCCACTTGGTCACAGCACAGAGCGGAGAAGGCTCAAGCAAAGCTCAG
GATATGGAAGCCGTGTGGGGCTACAAAGGCATGCTAAGAGACAAAGAGAGATGA
AGTCCAATGCGTAGCGGAAGATAGAGC
>index81_0
TTTCTGTTGGTGCTGATATTGCTAAGTAGTCAATCTGTTTACCGTTCTTAGGGA
TCCTCAGCATCCTTTCCGCATGATATGAGAGGGGAGAACCATGAAGGCGAGCTT
CAACTTCAGATTTGAACCTTTGTGAGTACACTCCAAGGCTCGAGTTACTCTGGT
GCGACCCTCTGAGATTGCTATGCTGAAGGCTTGACCTCATAATGTCTACCTTCA
GAGTTCAAAATGGTAAGAAGATAGAGC
>index82_0
TTTCTGTTGGTGCTGATATTGCTAATACCAGACCGATCTATGGCCGTCTTCGAT
GTCTCAGTGTGGAACCTAACACCCCATTGATGGAGGTCGACACAAGGCATGTTG
CTGGATGGGGCAGGAATATGTACAGCTATGCCCCAAGGCAGATTCGGAGCCCCA
CTCATTGAAAGCCGTGCCAGACCGCAAGGCCATGCAGTTTAAATACGGGGTCTT
ATTGAATGCGATGTGTGAAGATAGAGC
>index83_0
TTTCTGTTGGTGCTGATATTGCTAATACCAGACCGATCTAAGGCGAAGAAAGAA
AGGTCCAAGGTAGGCTTCATCGCGACAGTAGCATTCTGGCTGCAAGGCACACAC
ATTTCCACTACGAAGGAGGGTTCTTACGGTACAGAAGGCCGATGATGAGCGGTG
ACGATGTCGTTCTTGTCAACAGGTCAAGGCTCTACCTTAAGGCACACCACCTCG
GGGCACGTAGTTTGACGAAGATAGAGC
>index84_0
TTTCTGTTGGTGCTGATATTGCTAATACCAGACCGATCTAATCGGACTTACCTC
ACCTCGCTTAGCACTGGTAGCAGTAACTGTGGCTAGGCCACTAAAGGCGATGTT
CGTACGGCTCCCTTGGAGGTCTGAACTAACGTGTAAGGCCATATGACGGGGCAA
GGCGTGAGTCCTCACGACACCTCTCAAGGCTGACGTTTAAGGCACACCACCTCG
ACGTGATTGGTCACTCGAAGATAGAGC
>index85_0
TTTCTGTTGGTGCTGATATTGCTAATCTTGTCCTCAGTACAAAGACAGACCCTC
CCATCGCTTAGCACTGGTAGCAGTAACTGTGGCTAGGCCACTAAAGGCGATGTT
CGTACGGCTCCCTTGGAGGTCTGAACTAACGTGTAAGGCCAGATAACCAGACTG
GGGTCATTTGGTTGGTAGTGTTAGAAAGGCTGTATTTTAGGCGTGAAGTGTACA
ACCTAGGTATGACCTAGAAGATAGAGC
>index86_0
TTTCTGTTGGTGCTGATATTGCTAATCTTGTCCTCAGTACAACATGTTACCGGA
GGATCGCTTAGCACTGGTAGCAGTAACTGTGGCTAGGCCACTAAAGGCGATGTT
CGTACGGCTCCCTTGGAGGTCTGAACTAACGTGTAAGGCCATATGACGGGGCAA
GGCGTGAGTCCTCACGACACCTCTCAAGGCTGACGTTTAAGGCACACCACCTCG
ACGTGATTGGTCACTCGAAGATAGAGC
>index87_0
TTTCTGTTGGTGCTGATATTGCTAATTTAACAGGCGAAAGAAAGACAGACCCTC
CCATCGCTTAGCACTGGTAGCAGTAACTGTGGCTAGGCCACTAAAGGCGATGTT
CGTACGGCTCCCTTGGAGGTCTGAACTAACGTGTAAGGCCATATGACGGGGCAA
GGCGTGAGTCCTCACGACACCTCTCAAGGCTGACGTTTTAAACTAACCCGATCG
CCGAGGAAACATATCGGAAGATAGAGC
>index88_0
TTTCTGTTGGTGCTGATATTGCTAATTTAACAGGCGAAAGCCGGCCTGAAAGAA
ACATCAACCACACAAGTGGGACGATGAGTGTCCACACTGATTGAAGGCTTACTC
GTTGCTGCTCCTCAACAGTACGAGGCCTGTATGTAAGGCTCTTCCACGTATGGT
CTCCATGCGAATACCACTCGCGCTTAAGGCAATATGGCCGTATAAGAGATGATG
TCAATAATCCCACACGGAAGATAGAGC
>index89_0
TTTCTGTTGGTGCTGATATTGCTAATTTAACAGGCGAAAGACCTCACCATCGGT
TGTTCGTTGACCTTACGGGAACGGACTACCAGTGTTGCGTAGCAAGGCCTCAGA
CACATGCTTGATCAAGGCCACCATCGAACGCCAAAAGGCTCACAAGTAAGTCGC
GCGCGAGAGTGTTCAGGCTCCTCTGAAGGCAGTGAACTACAAGGCGTGAGCCGT
GTAAACGTTCTTCCGCGAAGATAGAGC

>index90_0
TTTCTGTTGGTGCTGATATTGCTACCACCCATCCTACAACAATCTTGTCCTCAG
TACTCATGCCACACGTACAGGAGGTGACTAGAAAGACTGCCGGAAGGCTGGAGT
TATTTGTCGGTTCGTTACCAACCGCTCTAAAATGAAGGCCCTCTGTACGCTGGC
CGCACCAAGTTGGGGACCTCTTTGGAAGGCGAGCTGTCGATTGTTAAGTCCCGC
CCAAATTTGGAACGTTGAAGATAGAGC
>index91_0
TTTCTGTTGGTGCTGATATTGCTACCACCCATCCTACAACAGTTAAGGAAATCA
AACTCTACGGTACAGCCTGTCACTGCTCTAGTCGGATACAACAAAGGCTTATCG
TTGAGGGCGGTGCTCAGTACCGTTATCACCTTCAAAGGCTCACAGTGTTGGCGT
TGTGGTATCTTCGTTGCCAAACTCAAAGGCAGGCGTCGTACAACGCTTTTCTTA
TCCTTCGATGTGTTGTGAAGATAGAGC
>index92_0
TTTCTGTTGGTGCTGATATTGCTACCACCCATCCTACAACAAAGAAAGGAGGAA
AGATCGCTTAGCTAGCTGAAAGACGATGACTGAACGACCCCAGAAGGCTTGACT
GAGCGGTGTGCTGAGGTCGTGTGAAGATCAAATAAAGGCCTCTTAGTTAGTGGT
CCATATTCCGGGGTCATGTGTTCCAAAGGCCGGTGCCACCTCATAAATTAAGCC
ACTAGCAGCCATTTAAGAAGATAGAGC
>index93_0
TTTCTGTTGGTGCTGATATTGCTACCGACGAAACTTTGTTACTCAATAACCCGT
AGATCAGCTCATCACATCGCGAGTAGTGCGACAAACCGTCGTAAAGGCCACTAT
GCCGGCTTCTACCAGGAAGCAGTGCGTACCAAGAAAGGCGCAAAGAGCCTCAGA
ACCTACAGCTCCGAGCCAGATCTGCAAGGCCTTCTGCTGAATACAACTCAGACT
TAGGGGTCTTGGTCGAGAAGATAGAGC
>index94_0
TTTCTGTTGGTGCTGATATTGCTACCGACGAAACTTTGTTAATTCCGTTACTCG
GACTCGCTTAGCACTGGTAGCAGTAACTGTGGCTAGGCCACTAAAGGCGATGTT
CGTACGGCTCCCTTGGAGGTCTGAACTAACGTGTAAGGCCATATGACGGGGCAA
GGCGTGAGTCCTCACGACACCTCTCAAGGCTGACGTTTAAGGCACACCACCTCG
ACGTGATTGGTCACTCGAAGATAGAGC
>index95_0
TTTCTGTTGGTGCTGATATTGCTACCGACGAAACTTTGTTAGGCGAAGAATCTT
TACTCGCTTAGCACTGGTAGCAGTAACTGTGGCTAGGCCACTAAAGGCGATGTT
CGTACGGCTCCCTTGGAGGTCTGAACTAACGTGTAAGGCCATATGACGGGGCAA
GGCGTGAGTCCTCACGACACCTCTCAAGGCTGACGTTTTATAGGCGAAGACGGA
ATTACTGAGTTGAACAGAAGATAGAGC
>index96_0
TTTCTGTTGGTGCTGATATTGCTACCGACGAAACTTTGTTTCCCGTTTAAGGGC
GTCTCTACGTCGATCCATTCGATCGTCGACAATGTATCTGGTTAAGGCTCAGAC
CTAGGCCGTATGGCGATTCGCATGTGCATGTAGTAAGGCCATATGACGGGGCAA
GGCGTGAGTCCTCACGACACCTCTCAAGGCTGACGTTTAAGGCACACCACCTCG
AGCAGCTATACAAGCAGAAGATAGAGC
>index97_0
TTTCTGTTGGTGCTGATATTGCTACCGACGAAACTTTGTTACCACGGAAATCAA
TCATCGCTTAGCACTGGTAGCAGTAACTGTGGCTAGGCCACTAAAGGCGATGTT
CGTACGGCTCCCTTGGAGGTCTGAACTAACGTGTAAGGCCATATGACGGGGCAA
GGCGTGAGTCCTCACGACACCTCTCAAGGCTGACGTTTAAGGCACACCACCTCG
ACGTGATTGGTCACTCGAAGATAGAGC
>index98_0
TTTCTGTTGGTGCTGATATTGCTACCTAATCAATCAATCATCCGCCTCAATTTG
CTGTCGCTTAGCACTGGTAGCAAGTGCCAATATGGCAGCGACTAAGGCTAGGTC
ACGAGCATGGACTGCTGGGTCTCTTCTCCAAGTTAAGGCACGCGGCAGTCTGAC
CACTTGATTTTCGACTCGACGTGCTAAGGCTCCTTGGAACCCCGTTAAGGGGCC
CTTTACCTTGATCGCTGAAGATAGAGC
>index99_0
TTTCTGTTGGTGCTGATATTGCTACTCAATAACCCGTAGAACCGACGAAACTTT
GTTTCACCACTTCTAGACCCCAAGGCAGTTATTGCATATCCTGAAGGCCCGGAG
TAGAAGACATCGCCGAGCTCTCTACGTTGGACGTAAGGCGCACGCTCGACTACA
GCTCATAGAAGGAATATGCACGCCGAAGGCCATGCCGTTGGTAGGTCTGTTGGC
AATGCTGTAGTGACCAGAAGATAGAGC
>index100_0
TTTCTGTTGGTGCTGATATTGCTACTCAATAACCCGTAGAAAGTAGTCACGGAT
GACTCACACTCGACAACGGATTCGTGCAGCTCACTGCATATGCAAGGCACGGTG
GATCGCTTTAAGAATCTCGGACCCACCGAGTGCTAAGGCTAATGTCCACCGGGT
ATCGGACGTGGTTATGTGCGTGCCCAAGGCTCCATACGTTCAGGTGTGGAGCCA
CAAACCGACATTTTGCGAAGATAGAGC
>index101_0
TTTCTGTTGGTGCTGATATTGCTACTCAATAACCCGTAGAAACCCTAAAGGGAA
GATTCGTACTGACTGGGATTGGGGACTCACAGACTAAGTAAACAAGGCTACGCA
GTGAGTGAACACATCCTGAACTGTAGCTGGATGGAAGGCCTTTCGTGTGAGGTC
TTTCAGCGGACTAGACCAAATGGGCAAGGCGGAACGATGATGTGGAGAATCGAG
CGACAACGACTATGAGGAAGATAGAGC

```
>index102_0
TTTCTGTTGGTGCTGATATTGCTAGGCGAAGAAACCAGACAATCTTGTTCAGCT
GGATCTACCGACTTCCTAGAGTGGCTGTCGATTCTCATCTCGGAAGGCGTACGT
GCCTATTCACTGATTAAAGCGTATATTTGGACATAAGGCCTTGAGGTCATGGTA
ACACGGCGTCGTCTCAGAGAAAACGAAGGCGTGGTACCTGGTAGAAACACAGCG
AGATCCAGGGCAGAATGAAGATAGAGC
>index103_0
TTTCTGTTGGTGCTGATATTGCTAGGCGAAGAAACCAGACTCCTACAACAGGGA
GGATCGCTTAGCACTGGTAGCAGTAACTGTGGCTAGGCCACTAAAGGCGATGTT
CGTACGGCTCCCTTGGAGGTCTGAACTAACGTGTAAGGCCATATGACGGGGCAA
GGCGTGAGTCCTCACGACACCTCTCAAGGCTGACGTTTAAGGCACACCACCTCG
ACGTGATTGGTCACTCGAAGATAGAGC
>index104_0
TTTCTGTTGGTGCTGATATTGCTAGGCGAAGAAACCAGACTCCTACTACCCACC
TCCTCGCTTAGCACTGGTAGCAGTAACTGTGGCTAGGCCACTAAAGGCGATGTT
CGTACGGCTCCCTTGGAGGTCTGAACTAACGTGTAAGGCCCGCCAAGGACAATT
TCGTGTACTGCCCTACGGTTTGTTTAAGGCTCGTAAACCATTATTTCATAATAC
AAGGAGGCTTTTACAAGAAGATAGAGC
>index105_0
TTTCTGTTGGTGCTGATATTGCTAGGCGAAGAAACCAGACAAAGAAAGGACCGA
AAGTCGCTTAGCACTGGTAGCAGTAACTGTGGCTAGGCCACTAAAGGCGATGTT
CGTACGGCTCCCTTGGAGGTCTGAACTAACGTGTAAGGCCATATGACGGGGCAA
GGCGTGAGTCCTCACGACACCTCTCAAGGCTGACGTTTAAGGCACACCACCTCG
ACGTGATTGGTCACTCGAAGATAGAGC
>index106_0
TTTCTGTTGGTGCTGATATTGCTAGTCGCTCACGGATGACTGTCGACGGAAAGA
TCATCGCTTAGCACTGGTAGCAGTAACTGTGGCTAGGCCACTAAAGGCGATGTT
CGTACGGCTCCCTTGGAGGTCTGAACTAACGTGTAAGGCCATATGACGGGGCAA
GGCGTGAGTCCTCACGACGAGTATTAAGGCTTGATTGCCCATAAAATCCTTCAG
CAATATTGATGTAGCTGAAGATAGAGC
>index107_0
TTTCTGTTGGTGCTGATATTGCTATCGTCATGGGTCATTTATTGCATTACCCTC
CCATCACACGTCTCGTAGAACACAGGGCTTTCAGTGTGGGGACAAGGCGAAAGG
CACTATATTTATGTATTGACTGCAATGGATCTGAAAGGCTCAGCATGGCTGACG
TTGTACCCTGGTCGGGTAGAGAGCGAAGGCATTGAGCCACGTTCGAAGGCTGAG
GCTGGTGGATAACACGGAAGATAGAGC
```

```
>index108_0
TTTCTGTTGGTGCTGATATTGCTATCGTCATGGGTCATTTACCCTCCCAAAGTG
GACTCAGGACAGACCTGTCACATCGGAGAGTGTTGGCTTGAGGAAGGCTCGTCA
ACGTGAGTTATGTTATATTTTCACCAACGGATCGAAGGCGAAGGAACCTACTTA
GCCCGTCAGGCGGTAGTATCGTCGAAAGGCGCGTATTCATGACTAGTTCTTAAC
CGGCGGCGCACTCCAAGAAGATAGAGC
>index109_0
TTTCTGTTGGTGCTGATATTGCTATCGTCATGGGTCATTTAGGAAAGAACGGAT
GACTCAGCTGAAGTCCTTTCTGCAGTGACTCTGCGAAAATGCAAAGGCAGGGCG
AATCCACGGGGCTGATGTTCACTAACCGGTCTCCAAGGCTTAATGGCAAGCAAT
GTTTTCTTCACGGCTTCTGTGGTTTAAGGCTGGCGCGCCTAACACTGACAACAG
ACAGACAAGACCTTACGAAGATAGAGC
>index110_0
TTTCTGTTGGTGCTGATATTGCTATTGCATTACCCTCCCAGGACATTTACAATG
ATCTCAACCACACGGTCAACGATAACGTTTGGAAGAGTTCTGAAAGGCAGTGCC
TCGGAGCGCCCCTGATCTTCCCCTACTGCGAACGAAGGCCATAAGCCTGAGTAT
TCACGTACCAAAACGTAGTTGAGAAAAGGCCATCAGGTTGCGCAGTTTCATAGG
ATCCCCAAATATGAGAGAAGATAGAGC
>index111_0
TTTCTGTTGGTGCTGATATTGCTATTGCATTACCCTCCCATCCTACAACAATCA
ACATCGCTTAGCACTGGTAGCAGTAACTGTGGCTAGGCCACTAAAGGCGATGTT
CGTACGGCTCCCTTGGAGGTCTGAACTAACGTGTAAGGCCATATGACGGGGCAA
GGCGTGAGTCCTCACGACACCTCTCAAGGCTGACGTTTAAGGCACACCACCTCG
ACGTGATTGGTCACTCGAAGATAGAGC
>index112_0
TTTCTGTTGGTGCTGATATTGCTATTGCATTACCCTCCCACCCACGTTCACCAA
CTATCGCTTAGCACTGGTAGCAGTAACTGTGGCTAGGCCACTAAAGGCGATGTT
CGTACGGCTCCCTTGGAGGTCTGAACTAACGTGTAAGGCCATATGACGGGGCAA
GTTCACTGTAGATAGGCATGATAACAAGGCTGCACAGGAACCCGTTGAATTCCA
GGGACCCTCACAGGTGGAAGATAGAGC
>index113_0
TTTCTGTTGGTGCTGATATTGCTATTGCATTACCCTCCCAACCTCACCAACTTT
GTTTCACCACTGTTAGAGGACCACATCTAACCGACGTACGAATAAGGCAAGCTT
GGGCTTAGACTGTAAACACGATGATCATCCTCGTAAGGCAGCCAACGCTTCCTA
CCTCATAAAACGGCTTGGTAATTATAAGGCACACTCCGTCCTGATTACGCGTCA
GGGTGAGAAGCCCACCGAAGATAGAGC
```

>index114_0
TTTCTGTTGGTGCTGATATTGCTATTGCATTACCCTCCCACCCTAAAGAAAGTT
CAATCATGGCTTCACAGTTGTCATCAGACGAACGCTTGCCAGTAAGGCTTGCGT
CGGCCAGCACAACATTGGAAGGGCATTTACAATCAAGGCGGTCACGTAGCCGAA
ACACGCCTTCCAGAAAACGACAGGTAAGGCTACAATTGTCCAACCTGAGATCGT
GGCCGGACCTCGCAAGGAAGATAGAGC
>index115_0
TTTCTGTTGGTGCTGATATTGCTATTGCATTACCCTCCCAACAATGATCGTTCT
ATCTCCCTTAGCTCGATTAGTCACCGAGAGGACTCGAATGAGTAAGGCTGGCCA
ACACTGCATGTTTTCCCGTGTAAAAGTGTATTTTAAGGCCTGGACTTTCAGGAT
TGCTAAGGAGATGACACCCATACGAAAGGCTGCTACCACGTCTAGATGGAAGCC
GCAGCTTTGACCGATCGAAGATAGAGC
>index116_0
TTTCTGTTGGTGCTGATATTGCTCAGGCGTTGGGTCTGAAACCGACGAAAGGAC
TACTCACCTGAGTTGTACACTCGGCCATCGTAAACACGGGGAGAAGGCGTCAGT
ATCAATCACTCACCTGTCGGTCATTGGCCCACGGAAGGCAAACGGCATGGAGGT
ATCATAGTTCTGAAGCTCAGATCGCAAGGCTCTTCTCAGGAGCGAGTCACGTGC
TTTGCTCTGTCCTTCTGAAGATAGAGC
>index117_0
TTTCTGTTGGTGCTGATATTGCTCAGGCGTTGGGTCTGAATCCTACAACTCCGC
CTCTCAGTCAGGATTCTCTGTGTCCTCGGGAGAGTATTACGTCAAGGCGCGGTA
CCATCCCATCAATCTCCGTCCCGAAACTAAACACAAGGCTACATCTCACGGTTA
GTTAATCGGGTGGATGGCTATCGCAAAGGCCAAGTCGCCCAATTGTGACCGCGG
GTTACTTCGTCCTTTCGAAGATAGAGC
>index118_0
TTTCTGTTGGTGCTGATATTGCTCAGGCGTTGGGTCTGAAAGGAAAGAAATCAA
TCATCAACGTCTGGCTTCTTGAGCTGCCACTCAAGTATCTCATAAGGCATCTTG
TGTGTAGGTAGGGTGCTGGACGACGGCAAGAATTAAGGCCTCTAGATTAGAGGT
TCCGAGATAGAGTAAGCTTTGCTGCAAGGCAGTCTACTCACAGAGAAGTGCGCA
GTGTAGAATGTCCGGGGAAGATAGAGC
>index119_0
TTTCTGTTGGTGCTGATATTGCTGGACATTTACAATGATCATTGCATTACCCTC
CCATCGCATGAAGACTTCTGGTAACTCGCGTACGGGTGGAGTCAAGGCATCCCA
GTGTTGCAGAATGGGCCATCCACTTTGACGAAGGAAGGCACCCTCGAGTCCACT
TCTGATGTTCGCTGGCAGTGAGGAGAAGGCGTGACATTGAGCTAGTCCTCGTGG
GTTGCACGTCAACATGGAAGATAGAGC
>index120_0
TTTCTGTTGGTGCTGATATTGCTGGACATTTACAATGATCACTCAATAAGGTGC
GTCTCCCAACATGGTGGATATCTACGCTACAAGCGGCGTTGAGAAGGCATGCTC
TCGCAAGCGCAACGCAATAACTTCGTCATCGGTAAAGGCCAGCGGACGACGCGT
TCCAACGGTAAAGCGTTGATGGGGCAAGGCCTCTGTTTATAGGTGGGCCGACAG
TTATAGGCGGTCAGAAGAAGATAGAGC
>index121_0
TTTCTGTTGGTGCTGATATTGCTTCAATAACATTGCGGATAGGCGAAGAAACCA
GACTCTGCAACACTACTGTGGGTCGAACGTAACATGGGGCGTGAAGGCGAGTGT
TAAGTGGTGAATTATTGTACCTCCAGGAGTGTTCAAGGCGTAGTCATCTTTTAA
CGTTACTTCGCTGAGAGTTCCGAGAAAGGCAAGGTCCAATTTGTAGGTAGTCCT
AGGAAGCACGTATACGGAAGATAGAGC
>index122_0
TTTCTGTTGGTGCTGATATTGCTTCAATAACATTGCGGATGGACATTTACGGGT
TTCTCTGCACAGTTCCAGGTCATAATCTGTCAGCATACGTGTTAAGGCAGCGCG
CTTTTAAGGCCAAATCGCTCATTGGTCACAACCGAAGGCTCGAACATGTCAGCA
GAGTTCACTGCGCGCCAGAGGCTGCAAGGCCTTCAGAACTACGCATGTCCCTGC
TAGGAAGCTCGCCACTGAAGATAGAGC
>index123_0
TTTCTGTTGGTGCTGATATTGCTTCAATAACATTGCGGATCCCACGTTCACCAA
CTATCATGCCATGAGAACACGGACTAGTAAGGTACGTAAGAGCAAGGCCCACAA
CATATTGTGAGACAACGAGGAGAGAGGGCACCCCAAGGCTCTTCTACACTCCTC
GTGACACAATTCTGCGATAATTTAGAAGGCTTCGTTCCAGCTGCACGAAGAGAG
GTGCGCACTTGAAGGAGAAGATAGAGC
>index124_0
TTTCTGTTGGTGCTGATATTGCTTCAATAACATTGCGGATAACGTATCATTGCC
GTCTCAGGACAAGCACGAAATCTTCCACCTTTGCTGAACTCTTAAGGCTGGTGG
CGTCTTGTGCCAGATGACTTCGGCACGGGGCCTTAAGGCCAAGAAAATTGTTCA
TTGCCCACTAGATCACGAACCGCTTAAGGCCGTCACGTCTGTCTTCTCACTATC
CCATCCCGTGAGCATCGAAGATAGAGC
>index125_0
TTTCTGTTGGTGCTGATATTGCTTCAATAACATTGCGGATACTCCGTGAAAGAA
AGGTCAGCTGATCTTCCTAGGTAACACGGGCCTAGATCTGGCTAAGGCGCTTAT
ACAACTAGTCGTTGCGCCGTATTCCGACCTCAGTAAGGCTCGAGGCCTGAGCTG
TAGCGTCATTCCATAGATCGGCTTAAAGGCTTGACACTTGGCTTGGGGAAAGCA
ATATCGTTTTAATTATGAAGATAGAGC

```
>index126_0
TTTCTGTTGGTGCTGATATTGCTTCCCGTTTAAATCAAACAAACCACCACCACG
ACTTCTGCAGTACGGTGATTAGCTCGCTCGAGAGATCGGATCGAAGGCCTGAAC
CAGTTCCGATTAGCAAAATATGGCTAGGCCTGAAAAGGCCTTTTAGTTACGCCG
GTTATTTAACTAACCAGCCCGGAGGAAGGCTACCCTTCATTGCTCTTATGCTAG
TACGCTCATTACACGCGAAGATAGAGC
>index127_0
TTTCTGTTGGTGCTGATATTGCTTCCCGTTTAAATCAAACGGGTAGGTTTCCCA
CTTTCATGGGAGAGCAGATCGTCCTTCTATCCTGAAGTCCCAGAAGGCTTAAAG
GCGATACAGGGGTACCGCTCAACCATATAACTTTAAGGCACTATTCAACCGATT
TGAGCGTCGCCAATTCATGAAGCTGAAGGCTACTAAACACACTTGTGCACCAGG
CCCATTTACCTAGACTGAAGATAGAGC
>index128_0
TTTCTGTTGGTGCTGATATTGCTTCCTACTAACCGTTCTTAATGCCTTGACCAT
CAATCACCATGTCTTCCCTGATAGTGGGGCTCAAAGATCAGTAAAGGCAGAGGT
GCGACGAAAATTACAGTCCATTTGGGAAGCACTTAAGGCCAGATCTGACCGCCT
GCTATAGCTGATAATGCCTATATGGAAGGCGCAAATTAACGCTGTCGATTCATC
CCGTAACACACACGGCGAAGATAGAGC
>index129_0
TTTCTGTTGGTGCTGATATTGCTTCCTACTAACCGTTCTTAAGTAGTCACCAGC
TTTTCATGCCACAAGGGTTTGTTTCTGACCACCTATTGAGCGGAAGGCTTGCGT
GGCGATCGCGACTCTTAGTGGAGAATAGGAGTCAAAGGCTAGCCTAATGAAAAC
CTGATATTACACGGTGGCGACCTTGAAGGCATCGAAAACGCCAGGTGCAGCCCT
TTAGGTGCCGAGCGATGAAGATAGAGC
>index130_0
TTTCTGTTGGTGCTGATATTGCTTCCTACTAACCGTTCTTACCGACGAAAGGAC
TACTCTGCACTTGCGATACTTCTCATGGCAGCTGTTGTCGTAAAAGGCTTCTAG
TCCGTTGATAGCGGCACCCCACACAATTAATCCCAAGGCCTACGCCGTCTCAAC
CTTTCGGACCGCTCGTCTGTCGGGCAAGGCCGTACAGCCACGATCTTCATTTCT
TCCACGATTTCAATTCGAAGATAGAGC
>index131_0
TTTCTGTTGGTGCTGATATTGCTTCCTACTAACCGTTCTTCCCTAAAGAATCAT
GTCTCGCTTGACATGTAGGGAGGAGATTACTAGTGCTTTCCCCAAGGCAGATAT
GTTCAGTCCAATCTACTGGTACCCGTGTCCGTTTAAGGCGGCCGAACCACAATT
CTCACTGTCCAGACCTCTTAAGAGCAAGGCACACAGTTCTGAATTTTGGTCGAT
GTTAGAGACAGGGTGTGAAGATAGAGC
```

```
>index132_0
TTTCTGTTGGTGCTGATATTGCTTGGCCGTCTTCGATGTCAACATGTTACCATC
TTTTCTGCACAGTCGAACACAATGACGGATGCTGGAGTGTCCCAAGGCAACATA
TCCACCTGCTGCTAGACCGACTTAAGCGTAACGGAAGGCCATAAGAGTCATCAT
AGAAACTATTGTCGGGATATACTACAAGGCGGAGATACTGTTAGCGCCCTGCTA
AGGGATACTAGGAGTGGAAGATAGAGC
>index133_0
TTTCTGTTGGTGCTGATATTGCTAAACGTATTAGGTTCCATCCCGTTTAACAAA
GGGTCACAGGAGAACCGTACCTCATCATACAGTATCGAGGTAAAAGGCCAGTGG
TGTGAGCCTAGGTCTTTCTTACTATTCACTCTGGAAGGCTTGGCCGCCGGCAAC
TTCACCGGGGTCCGAAGAAAGTACAAAGGCAGCTATGTGCACGGCTTTCGGAAG
GTGTACACTCATGATAGAAGATAGAGC
>index134_0
TTTCTGTTGGTGCTGATATTGCTAAAGAAACAACGGATGAAAGACCCGACAATG
ATCTCATGGTGTGAAGTGCGTCACGACATAGAATTGCTCAACCAAGGCTAACAG
GCCTATGCAATTGCCAGGGTACGCTATGGAACCTAAGGCCCCCTATACCAAGTT
TGATTGGAGATAGTTCTTGTGACTCAAGGCGGTTCAATATTCTGGTCTATTCCG
ATCAAGACAAGCGCCAGAAGATAGAGC
>index135_0
TTTCTGTTGGTGCTGATATTGCTAAAGAAACAACGGATGAACGGATGACATTGC
ACTTCAGCTGACTCTGCTCAATTGATGGAACAGAACTGCCAGGAAGGCACACTC
TACAGGCAACGCTTGCAATTTATGAGTAAGGTGCAAGGCCTACCTGCACGAGGG
CATTGCGAGGCGTATCCGGTATGCTAAGGCGACAGTGCAGCAATCGTGTAGCAG
AGTGTTGCGAGGCTCGGAAGATAGAGC
>index136_0
TTTCTGTTGGTGCTGATATTGCTAAAGAAACAACGGATGAACCCTCCCACCCTA
AAGTCAACCCATCGCACTTTCCTTTTCTGAGACGGTCGTCAGAAAGGCATGCAG
GCAAAGCATAGTAGACTAAAGATGCTGCCTTCCCAAGGCCAGGCCCGCATGAGG
ATATGCCAGAAGTACTCCTAAGGTAAAGGCTAAAGCCCCACTATACGTTCAGTC
GATTAGTGTAGTCGAGGAAGATAGAGC
>index137_0
TTTCTGTTGGTGCTGATATTGCTAAAGAAACAACGGATGATCCGGAGGATTTCA
AAGTCACCTGAGTTGCGTACTATTCGTACCTCTTCCGCCTGATAAGGCGTAGGG
GTAGAACACTGCAGCCGGGACAAGGGTTTGGCTTAAGGCAGTAGCGCGTCACCA
ACCATCTCAACTATCGCTCATCTCGAAGGCCGGTGAATAAGACACGCGGCGGGA
TGCTGAGCACTGACATGAAGATAGAGC
```

>index138_0
TTTCTGTTGGTGCTGATATTGCTAAAGATACATCCCACTTAGGCGAAGAAAGAA
AGGTCGTTGTGTGGTTAGTGGCTTAACAGGATGGGGTTAAAGCAAGGCGGAGTT
ATTGTCGCTGGAAAAGGTCGATGCTATGCGCCCCAAGGCTCCAGACAGGCCGAG
ATCACGGCAGCCTCTATGTAATAGAAAGGCTCCAATTAATGAATCATTTGGCGT
GTAATCGAGAACCTGAGAAGATAGAGC
>index139_0
TTTCTGTTGGTGCTGATATTGCTAAAGATACATCCCACTTTCGATGGATCACCC
ACGTCCACTTCGTTTCGTTGTGCGAACGACCCAGAAATACATAAAGGCCATGAG
ACCAGAATCCGAGCATTCCTGAGGGTATCTATGGAAGGCTGGGCTGGACGCGTT
TAGACGATATCGAGACTCCATTAGAAAGGCATGATCAATTTCAGGACTTCATGT
TAACCTCATTGTCACCGAAGATAGAGC
>index140_0
TTTCTGTTGGTGCTGATATTGCTAAAGATACATCCCACTTACCCTCCCACCCTA
AAGTCTGCAGTACCGTAACGAATTCGGGCGAATCACGAATATAAAGGCACCGGA
CTATCTCAACGTACGCGGACCATCGTATACTTGTAAGGCTGCCTACTTGTCTAA
CTCCCCTGGTCGTGCGTATACGTGCAAGGCTGGTCACTTGGTCAGCGACCATGC
CGTGACTCGCCGATGAGAAGATAGAGC
>index141_0
TTTCTGTTGGTGCTGATATTGCTAAAGTTGCAAGGAAAGAACCACCCATCCTAC
AACTCTACCCAAGTGTCCATTCCCCGAGAATGAGTATGAATGCAAGGCTCGGAA
GCGCGGTCCTCGTTAGTGCTGTAGACGCCCACAGAAGGCCTATCTGTGTGCTCA
TGGTAATGGGGTTAGACGCTCCAATAAGGCCCCTAGGTGTCGTGGCAACCCTCT
TTCAAGAGATACGGTTGAAGATAGAGC
>index142_0
TTTCTGTTGGTGCTGATATTGCTAAAGTTGCAAGGAAAGAACCGTTCTTAGGGA
TCCTCAGCATGTCCTTACGTGTCATTGTGAGCTCTGCTCTAGTAAGGCTGCGTC
ACCCTTCTCGCCAAGGTCGGGTGTGAACCTCTTCAAGGCCAGAAACGACGGACC
CTACGGATGTGAGTTTCTACGACCTAAGGCGGTTATCAGTCGACGAACTGCTCG
AGTTTTCACTGGGACAGAAGATAGAGC
>index143_0
TTTCTGTTGGTGCTGATATTGCTAAAGTTGCAAGGAAAGAAGGCGAAGAAAGAA
AGGTCATGCCATCAGGACATTCATCTGACGGTAATCTGCGTGTAAGGCGATTGA
TGCGCAATTAGTCTGGGTCTATTCCCACGGACACAAGGCAGGACCCTCGTTCGG
TCAGATTGCTCTGTCCCCTTCCGCTAAGGCAAAGGTGTACAGACTACGAGATCG
CGGGATTGTCTGGAAAGAAGATAGAGC
>index144_0
TTTCTGTTGGTGCTGATATTGCTAAAGTTGCAAGGAAAGAAGGGAGGAAAGTCC
CACTCACAGTCCTCCTGTGAGTCGAACCTCACATTGTTACTTCAAGGCTGGTTG
ATCTGCTGTTGCTTTGTCATACGCGCACGCCTGGAAGGCGAGGGACATTTCCGA
CGTCGACGTCTGTCCTCCCTCTTTCAAGGCGGTTGATTCGTTAAGTTGAAGCTA
TCAATCCAAAATCGTAGAAGATAGAGC
>index145_0
TTTCTGTTGGTGCTGATATTGCTAAAGTTGCAAGGAAAGAAAGGACGTTGTCGA
CGGTCACCTCTTGCACGTACTATACGATGCGTGATTACAGCGTAAGGCCCACCA
GACTTCACGCGAAAAGAGGATTCGAGACAGAACGAAGGCATGCCTTGTGCTAGT
TGATGTCACACATCCCGAATATAATAAGGCGGAAGAAGTGTCACCGAAGGCATA
AGAATATCAGTCATCCGAAGATAGAGC
>index146_0
TTTCTGTTGGTGCTGATATTGCTAAAGTTGCAAGGAAAGAATGGGTTTAACTCA
CCCTCATGCCAGATCGGATATGACTGTAGTGGCGTATCTTACAAAGGCTGGTGT
CTGAACTCTACTGTTGTCGTGTCTACCAGGATTAAAGGCGCCTTCAGACGCTCA
CTAGTAACCAATTGCACAATCCATCAAGGCTAGCTCGTAGCATAACACATTGGA
GAAGGCATGATGTCAGGAAGATAGAGC
>index147_0
TTTCTGTTGGTGCTGATATTGCTAAATTAAAGAAACCACCAATTCCTCATCCGC
CTCTCCAAGGACAAACGTAGACCAGCGGATCACGCAAGCAAAGAAGGCACATCC
TGGGCTATTGAGCAAAAGGAGCTATAAAGCTATAAAGGCATACTGACCAACTAC
AGACATCGAAGACCGCAATCTTCCTAAGGCTAGGTCAAGTTTGGGGTCTTGCAA
CTAAGAATGTCGTACCGAAGATAGAGC
>index148_0
TTTCTGTTGGTGCTGATATTGCTAAATTAAAGAAACCACCAGGGAAGATTCGTG
CTCTCGCAACTCTGTCGTGTGAGAGTGGTTGCAGCACTAGAAAAAGGCAGTTCT
ATAGAAGGAGGATTCGACCTCTATCGTACCCGTTAAGGCAACAGGGGTGCTCCT
ACTCCGAACAAGGGGCTGGCACCCTAAGGCATTCCACGTTTGCACACTCCGGAC
TGAAACTTGGCAAAGGGAAGATAGAGC
>index149_0
TTTCTGTTGGTGCTGATATTGCTAACAGATCAACTCATCAACCACTTGCGTTCT
ATCTCGTTGTGTGCGTGATGCTGACCAACTCTGTCTGCTCCTTAAGGCCAACTC
GTCCTAGTTCGGATGCAGCGGACTAGTGTATCGGAAGGCGAGTGCGTTAGTCCT
CGTGCGTCTTCCTCATCCATTGTATAAGGCACTGAGTCCGCCCAGCCCTAGGAC
CTCACATAGGGCTGTGGAAGATAGAGC

```
>index150_0
TTTCTGTTGGTGCTGATATTGCTAACCTTTCATGGCCGTCAACTCGTTAATCAT
GTCTCAGCATCCACGTGTCAAAGAACAGTACGCCAACTAGGATAAGGCAAAGGT
CCCCGCCTAGTACGGGATTCAGCTCCTCCACGATAAGGCCATTATGTCCACTTC
ATCTGAGATCTGTGGATCCCCATTAAAGGCCATTGGTGTAGTGTGCGATTGTGC
GCCGGTGAGGTTCCTAGAAGATAGAGC
>index151_0
TTTCTGTTGGTGCTGATATTGCTAACCTTTCATGGCCGTCAAAGAAACAACGGA
TGATCAGCTACACCATTCCCTTAGAACTGCCCTGTCGTAGTCGAAGGCTTTCCC
CTCGTTTAGCACATCTTCGCCGCCGCTATGTACCAAGGCCTCGGTACTGTCAAA
ATCTCACACACAAAGTTGGATCAGAAAGGCGTGACTATACGAGCCCACCTGTCT
TATACCAACCGCCATTGAAGATAGAGC
>index152_0
TTTCTGTTGGTGCTGATATTGCTAACCTTTCATGGCCGTCAATCAACAAACAAA
GGGTCAACCAGACCCACATGCTCCATGCAAGTCATCCACACCCAAGGCTACCCT
GAGTATCCCCTGAATACTATGTACAATTTGGTAAAAGGCCCCTTGTAAACGCGA
TCAATTACTCTACCATGTTCCTTCCAAGGCACGCCATATTACGTGGACTGCACC
CGACCCGGATTACGCGGAAGATAGAGC
>index153_0
TTTCTGTTGGTGCTGATATTGCTAACCTTTCATGGCCGTCCACCCACGATTGCG
GATTCGTTGACTCTTCCTCAGGCCTATATAGGTGGGACCACGAAAGGCACGATA
CTGGTAGGATTCTATAACATTCTCGTCCAGCGCAAAGGCCTAAGCCAGTGATCC
GATACCGAAGGTCGGAAAAGGGCCAAAGGCAGAGGGTTCGTGCAGGTCACGACT
CTGGCTCATCACTCCTGAAGATAGAGC
>index154_0
TTTCTGTTGGTGCTGATATTGCTAAGGGCGTCAATAATTTAAGGGCGTCGGACA
TTTTCATGGGTTGTCACCGAACGGTACTGCAGTTTTATCATCTAAGGCCTCGAT
ACGAACGGTGTGGATGGCAAATACGGGCTTATTTAAGGCTGGCAAGGATTGACG
CGTCTAGAAACTGCTCGGGGATGTGAAGGCTGTATCGGAATGGGCTAATAATCC
CGGTACAGCCCTGCGCGAAGATAGAGC
>index155_0
TTTCTGTTGGTGCTGATATTGCTAAGGGCGTCAATAATTTCCCTAAAGCCCTCG
TTTTCACGATCAGTAGTGCCCTTCCCATCTCCTGGTATGTGGAAAGGCCTCCGA
AGTGTCAAATATAGCCCTGGACTAGTACGCGGGTAAGGCGACCCCAGATTAACC
TTATGGACGTTTATGTATGGTATAAAAGGCATTCTCATTCTTCATGTGTATCGC
ATGTTCGACGCAGTGTGAAGATAGAGC
```

```
>index156_0
TTTCTGTTGGTGCTGATATTGCTAAGGGCGTCAATAATTTATCGGTTGTTCCGC
CTCTCAGCATCCAGTGGTTCGTGCTGTACAAGCACCATCGCACAAGGCCAACAC
GACAATATTGGTAGCAGGATCGCTCATGTAAAGCAAGGCTGCAGACGTTCGTGG
AATCTGGTTCCTCCAATGTAACTCCAAGGCGCGCTTGACCCCAAAGGAAGAGGA
TCATTCTCAGTCTCTCGAAGATAGAGC
>index157_0
TTTCTGTTGGTGCTGATATTGCTAATCAATCAATTCCGGGAACCTTTCATGGCC
GTCTCATGGGTTGTCACCGAACGGTCTGACTCTTTAAGAGGGCAAGGCCGCTAA
ATTGGACTTATCGGAATTCGTTGCATGTATTAGTAAGGCAGACCTCTCGTCTAC
TCCCTGGGGCCAACAGAGTTTGTATAAGGCTCGCATTTCTTAGACTGGGCCCCT
GACTTACGTGCGCCTGGAAGATAGAGC
>index158_0
TTTCTGTTGGTGCTGATATTGCTAATCAATCAATTCCGGGAGGGAGGAACGTTT
CAGTCACCTTGCTGGAGAGTTGGGTCTGTATCAGACGAGACGGAAGGCCTGTAG
TCTCCAAGCGTACCTTTAAGCAACACCTCATTTTAAGGCCTGTATTCAGTTGAA
AGTCAAATCCCAACATCAGAACCCAAAGGCCCGTTACCCGCTAACAGCTGTGGA
ACGGATGATGGGAGCCGAAGATAGAGC
>index159_0
TTTCTGTTGGTGCTGATATTGCTAATTCCGTTACTCGGACAAAGTTTCTTCGAT
GTCTCACGATCTGTTCCGCACACAATGCAAGAATCGGTTTACGAAGGCAAGCTC
AGCTCTGCTAGGTTTTCTTCTAAGCGTACTCGAAAAGGCGCGGGCTGATTTGAA
AGCATACAATTCATCCTGTAAGACTAAGGCTCGGTCTGAACTGACCCCACCAAC
CCGCTTGTGAGCCTGCGAAGATAGAGC
>index160_0
TTTCTGTTGGTGCTGATATTGCTACCATCTTTACTGTTTCATCGTCATTCCCAA
ACCTCTACCACTGCCTGAACCAGCTACCTTGTACTGCAATTCTAAGGCATGTCG
TTAGGGTATCGACTCTCACCCTCCGGTGTTTCAAAAGGCAAGCTCTCGTCCCGC
AAGGGGTCGGAAGTCTCCCACATCAAAGGCTCCCGGGCACTGCGTCAAGACTTT
CCGCAAAATTAAAGGAGAAGATAGAGC
>index161_0
TTTCTGTTGGTGCTGATATTGCTACCATCTTTACTGTTTCAAGGGCGTCACCAA
TCGTCGTTGACACTGACAGATGGATGAACTCGATCCTTCAGACAAGGCAACTAG
GAAGCAATGCAGCCTAACGACCAGCATTTCAAAGAAGGCCACGTAGCGGATGGT
TCGACGATGCCTTCTAGGAATTGGCAAGGCCCACCGCTGCCAAAAGAGAGTGGT
TGTGGACATACGCACAGAAGATAGAGC
```

```
>index162_0
TTTCTGTTGGTGCTGATATTGCTACCATCTTTACTGTTTCGTTTGCTTAAGTGC
CGCTCGTACCTTCGGTCTCACATCACCCTGTAACAAGGACTCGAAGGCACGTTG
GCAAGTACATGTGGGCGTGGTCGATACTTTATGAAAGGCATAACTACCCCACCG
TCTTTGGGACTATCTAATGACCGAAAAGGCAGCGTGCGATTGATGCACGATTTT
GCGGCTCACTGTTTGAGAAGATAGAGC
>index163_0
TTTCTGTTGGTGCTGATATTGCTACCATCTTTACTGTTTCACCCTCAGAAAGAA
AGGTCAGCTAGCTCGTTGGAGATACTAACACGTAACGTGAAGGAAGGCCCGCTG
TCCTATTGGGTACCTAAGATGACTTGTTTGGGCTAAGGCGGGAGTGCGCAAGAT
GCTTCGTATCGTACTGGTGGTCTGGAAGGCGCTGCTGTTCTCATTTTCGGGGTG
CTTTAAGTGTAGGTTGGAAGATAGAGC
>index164_0
TTTCTGTTGGTGCTGATATTGCTACCGTTCTTAGGGATCCAAAGCTATCATCGT
TCCTCAGCTCAACCTGTATGTCATCACCCTGAGGACCATCGAGAAGGCACGAAC
CTAACGTATGCGTGCGAACCGATACGACCTAGCCAAGGCCACCATTTTCGTCCA
GAGTTTTCTGGTTCAAAGCTCGCGCAAGGCCTAAACGGGGAACCTCCTGTCCTA
GTGTCTTCCATCTCACGAAGATAGAGC
>index165_0
TTTCTGTTGGTGCTGATATTGCTACCGTTCTTAGGGATCCAAAGAAAGGACCGA
AAGTCACCACTGAAGACACCCAATTAGACGATGTCGGGAGATAAAGGCGACGGG
GTAACAACTCATATTGCAAATCACACTTCCCTTGAAGGCATAGCCGCGTTCAGA
AGGTACCTCGGCCGCTGATCGCTGCAAGGCGTGCGCCCTTCCAATCACGCTCGG
CACGCAACCGTCTTCAGAAGATAGAGC
>index166_0
TTTCTGTTGGTGCTGATATTGCTACTGTTACCAACTACTCAAAGTTTCCACCCA
CTCTCACCTTCCATCTGTACTGACTCTCCGTCAAGGGCAGAGTAAGGCTTGGGT
GACTAAGTGACCCGTTGTGATATTTATGCAGGGGAAGGCTGTGACCAGCCCTCT
GCGGCTGCGGTCATGTTCTTGTTGCAAGGCCGTTGTGTCAAACCCTATAATAAT
AACACGCGTCTTAATGGAAGATAGAGC
>index167_0
TTTCTGTTGGTGCTGATATTGCTACTGTTACCAACTACTCAACTCGTTCCCACG
TTCTCAGCACACTATGGGTTGATCATGGGCATATTCGAGCTCCAAGGCTGAATA
GCTACGCTACATCACAGAATGAGCATGTCTAGCGAAGGCCACCTTGAAACGTTC
GAGCATTGCTTCGAAATGTGTACCGAAGGCCGGCTAGATCTCACGGACCGACAT
TCCGTGGAGGAGTAAAGAAGATAGAGC
```

```
>index168_0
TTTCTGTTGGTGCTGATATTGCTACTGTTACCAACTACTCAAACTCAACGTTCC
CACTCCAAGGTTCGTACACCAATTGAGAGGAACTGGTCGGGGTAAGGCTAGGCA
ATGTGGGCAGGAGTCACTCCGATCGCATGATATGAAGGCAACATCGTCGCAATA
GCCCGTGCAATTGAACGACCCTCCTAAGGCACCGATGTTTTGGGCTGTCCTGTA
GAGCCTTGCGGGTATAGAAGATAGAGC
>index169_0
TTTCTGTTGGTGCTGATATTGCTAGGTGCGTCAGGAAAGAAACATGTTACCATC
TTTTCAGTGAGTCTACGTAGGCGTTGGAATCCCGTTCATTGTCAAGGCTGAGTG
AGGTCGTATCTACGCCATCCCTCAAGAACAACGAAAGGCAAGCTCAGCCGAAAA
GGCAAGTATCGGTAGTAAATGGCAGAAGGCTGTCGGCGCGAGTAACCATTTTGC
ATCTGTTACCCGATAAGAAGATAGAGC
>index170_0
TTTCTGTTGGTGCTGATATTGCTAGGTGCGTCAGGAAAGAAGGCGAAGAAACCA
GACTCATGGTGTGAAGTGTCAGGCAAAAGTCAGACGATCCCGCAAGGCCCGATA
ATGTAACTTTTGGGGCGTACGACCTGTAAGAGTGAAGGCAATACCTTAGGTTAG
TTTCCTATTAGGCAGCACTCATAGAAAGGCCGATATGTTCACTGTGCAACAGTA
TGAGAGTTGTGCGTCGGAAGATAGAGC
>index171_0
TTTCTGTTGGTGCTGATATTGCTAGGTGCGTCAGGAAAGAAGGTGCGTCAGGAA
AGATCGTACTGCACGTCTATACATGTCCTTCGCAACCGTCCCTAAGGCCTCCCT
AGAAGTCTCCAGTAATGTAGACTCTTTATGAGCCAAGGCCTACTTGTGCAATTG
TACCGTATTCATAGAGGCCCGCTCCAAGGCTGGGATAGTGAAGAAAAGCTTCAA
CGGGGTGTAGGCATTAGAAGATAGAGC
>index172_0
TTTCTGTTGGTGCTGATATTGCTATTTCAAAGATTGCACTACCACCCATCCTAC
AACTCAGCTTCGTTCAGCTTACGCTCTCCTCTGCACACGTAAGAAGGCAGTGAT
CGGTTCGTTAATCTGCATACATTTTCTGGAAGCAAAGGCAACTTCTGGTGGTCA
TCTATCACCACCTATGTAAGTCGCCAAGGCCGGGCACTATTAGGTTGTAGCCCC
GCGGGTGGTGCATACTGAAGATAGAGC
>index173_0
TTTCTGTTGGTGCTGATATTGCTATTTCAAAGATTGCACTACAGACTCAACCGT
CCATCCCTTACAGGCTTGTTTCCTATCAGGGTGCTTAATGTGTAAGGCCGCGAT
GCACGACGTATTATTGATGCAGTGCTCTCAAGGGAAGGCGTTAGTTGCATAAGT
GGCACTATGTAAGTCGTAGGAGGCTAAGGCAAGCTCCATATACATTCCAGCCGA
TCCTACTTGGGAAAAGGAAGATAGAGC
```

```
>index174_0
TTTCTGTTGGTGCTGATATTGCTCCCACGTTCAAATTTCAAAGTCCCACGTTTG
GTCTCGTACTGAGGGACTTGAACACTTTGGACTATGCGCCCCAAAGGCACCTGC
TGATGGAGACCATTAGCTAAGGTGCTTATAACCTAAGGCTTCCATCGGCTTCCC
AGGTCGTATGCTACGCTGCGGCGTCAAGGCAATTACACGACGAATTAGCGAGAA
TGTCCCGGATATTGTCGAAGATAGAGC
>index175_0
TTTCTGTTGGTGCTGATATTGCTCCTCAGTACGTTTGGTCAGTTAAGGAACCTT
TCATCCCTAACGAGGAATGTACAGGACGACATACCGTCAGTTGAAGGCATGCGT
GGTGTTTTCCCAATCCCCTGCGCAAGGTCAGTTAAAGGCAGCAAAGGTTGCGGC
CTATTTGGGACGAAAACAAGAAGTTAAGGCATTGATCCGTTAGACTAACTGTGC
CCTTTCTCCGTATGACGAAGATAGAGC
>index176_0
TTTCTGTTGGTGCTGATATTGCTCCTCAGTACGTTTGGTCAAAGAAAGGAGGAA
AGATCAGCTAGTGCTCACACACAGTACTAGTGTCACGACGCGAAAGGCGATAGC
ATTGCAGAATGGTCTTTCGACCGCGTCGTTCCCAAAGGCATCTGTGAAACCCTT
GGACAAGTCTCGCTACCGGTACTCCAAGGCTTGCTGATCCAGTGATCCTGTTGG
ATCTAGTACTTGATATGAAGATAGAGC
>index177_0
TTTCTGTTGGTGCTGATATTGCTTCCAAATTTAGTTTACGCCCACCTCCCGGAC
TTTTCACAGTCACGTATCGGAAATGCGTTCTCCGCGCTTTCGCAAGGCAACCAA
TGATGAGATTTAGTTTTGCCCCGTCCTCTCATCTAAGGCCTGCTACAAGTGAGA
AAGCCGCGGAAGGCAGGGCCCTTGAAAGGCATGGCACGTTGGCATTGTCTCCGT
GGGTGACCTGGCGCAGGAAGATAGAGC
>index178_0
TTTCTGTTGGTGCTGATATTGCTTCCTACAACAATCAACAACTGTTACCTCAGA
CTCTCCACTGTCTCTCTATGCGAGCACTTCAACAACACACAAGAAGGCCGGCGA
GGAGCACAGATCTCCCTAGTACGCGAGCAGTGAGAAGGCCAAAACGGGAATATT
CAGGGTTCTTGTAAACCCCGTTTGTAAGGCCTGACCGACGGGTGGTCGCCAAAC
CAAAATTGTTGTGGTAGAAGATAGAGC
>index179_0
TTTCTGTTGGTGCTGATATTGCTCCGGCCTGAAAGAAACAACCATCTTTACTGT
TTCTCATGCTGCATCCTGTCGAAGTTCAATGGCGTAGCGAGATAAGGCCTAAGT
AAAGAGCACTCGAATTTCTGACTCCGCTTAGTCGAAGGCCAGCGCCGTTCAATG
GCGCAGGTCTAGCCAGCAAATCACGAAGGCTTGAGAGCTTCTCGGTGAATATGA
TGCTATGGATATTCGGGAAGATAGAGC
>index180_0
TTTCTGTTGGTGCTGATATTGCTCCGGCCTGAAAGAAACAGGGTAGGTTAAACG
TTCTCCAAGGATGTACAGCTCTCAGCCAAGCGATAGTAGACGAAAGGCCCTAAA
GGTTGTACTGGGGAGTAAGCGCCTTCCCCTTAATAAGGCTGACAGGCCTGAAAC
CTCCTGATGTACTGGCGCATTCGAAAAGGCCTTCGATAGCATTAACTGTCATAC
CAGCACCACCAGCAGGGAAGATAGAGC
>index181_0
TTTCTGTTGGTGCTGATATTGCTCCGGCCTGAAAGAAACAACCACGGAAAGGAC
TACTCCAAGGTGAACCCTCTACTAATCCTGCTCCAAGCGCTCTAAGGCTTTGAG
TCTGGCGGGTTTTGATCCCCTAACTAGTGGCACAAAGGCTTTCGCGTTGGCAAT
TTGTTATCTGCTCAGCTCAATGACTAAGGCCTTGGTGCAGTCCAAACGACAGTA
GATTCATGGCCTCTACGAAGATAGAGC
>index182_0
TTTCTGTTGGTGCTGATATTGCTAAACCACCAATGTCGTTTCCCGTTTAACAAA
GGGTCAACCCATCCGTTGATTCAGAGACCACGACAGGTGTGACAAGGCCCGAGT
CATTGTGTGTGGCCCATGCTGTACGGTGTGACAGAAGGCGATTTCATGGGTCCA
CGGATAATTTAATGTCCTAGCCCGCAAGGCCCACGTCCTATCATGATTTGGGAA
TCAATAGCCGCTCAATGAAGATAGAGC
>index183_0
TTTCTGTTGGTGCTGATATTGCTAAACCACCAATGTCGTTGGCGGAGTAAAGAA
ACATCATGGAGGTGTAGTTGCGTGTTTCCTGACTTCCTTTAGGAAGGCCATCTG
TGGATCGTATCGCGTCTAATACCCGGGGAAGAATAAGGCGCAAAACAAGAGTTC
GCCTGTGTGGTGGGTGGCACTCCTCAAGGCACAAGCCGAATAAAGCCGCAACAA
TCATACTGATCCCATAGAAGATAGAGC
>index184_0
TTTCTGTTGGTGCTGATATTGCTAAACCACCAATGTCGTTAGGTGCGTCATCGT
CATTCATGGAGTGATCTCACTGTGACTGCTTCACGTTGGGATGAAGGCGGGAGG
ATCTATTTTAATGTGTCCTTACTGGAGTGCTCTAAAGGCTACTCGACCGATATG
GAGAATTACCCTGAGTGGGCGGCGTAAGGCTGCAGCTCTCCACAGCTATCGAAG
ATAGTTTAGGTTGTTCGAAGATAGAGC
>index185_0
TTTCTGTTGGTGCTGATATTGCTAAAGACAGACGTTTCAGGTTTGCTTCCCACG
TTCTCGCTACTCTGTAACTGCGACTTACGATTGTGCGATGTACAAGGCTTTCAC
GCGGGGATTATAGGGATTACAGGCCACCCCGGTCAAGGCTCGCCTCCGTGCAGA
AGAGAAGTCAGCCGAAGTACGTGGAAAGGCTCATCGCAATCTTACTACCCTGCG
ACAATACGATTAGTGCGAAGATAGAGC
```

>index186_0
TTTCTGTTGGTGCTGATATTGCTAACATGTTACCGGAGGAAACTCGTTAATCAT
GTCTCACCTCTGAGTCAGTTGCACGGACAAGAGTAACCTTGCTAAGGCGGGAAT
CTACGGCACACTTAGTGTCGCCAACCATAACGAAAAGGCCGGGCCCTCCGGTCA
TGCTGGAGAAGTCGAACGATCGTGAAAGGCAGTTAACTGTCGCCTTTCGCGCAT
TCTCGCTAGGGCCCAGGAAGATAGAGC
>index187_0
TTTCTGTTGGTGCTGATATTGCTAACATGTTACCGGAGGAAATCAATCAGCTTT
CCATCGTACCTTGGAATCTCAGGTGATGACTGGACAGCCACTCAAGGCCTCGAT
GCACTCGCTTAATACCAGTTCACCGACGCGAGTTAAGGCCGCTCTCCCTAGATG
CAAGACGAATTACACCAAGACAGCCAAGGCTAGTAAGTCCCAGACCATGGTTAC
GGTCCGACACGTAATAGAAGATAGAGC
>index188_0
TTTCTGTTGGTGCTGATATTGCTAACATGTTACCGGAGGAAATGGAGGAACTAG
TTCTCATGGCTGTGTAAGTGGCGGACACAGTGACAGTACATAAAAGGCATACGA
AGAGTGTTGCAGGAACACTACCAATACTGTGGTTAAGGCCTAGTCACACGGCAC
CCTAGAACCTTGTGCCAACCAGATTAAGGCAGAGCACCAATAACTTTGAAACCA
GCTGCATGTACCATTCGAAGATAGAGC
>index189_0
TTTCTGTTGGTGCTGATATTGCTAACATGTTACCGGAGGAGTTCCCACTCGATG
GATTCATGCTGCATACTGGGAAGTTGGAGGGTATCAAGCAAGTAAGGCTATGAC
CTACTTGTCCACCCTTGAACGCTAGAGAGCGTGAAAGGCGATGGGGATGGGTAT
CGCCTATGGTGGCTCACAGGTCTCTAAGGCTGGCTAGTTAATTAATTAAACTGG
ATTACTCGGTTGCAACGAAGATAGAGC
>index190_0
TTTCTGTTGGTGCTGATATTGCTAACATGTTACCGGAGGAAGGGAAGATAATCC
GTTTCACCTTCAGCAAGCTGGTCTTCTGTGATTCCGTTGCGACAAGGCGATCGA
TAGAATGGGATTTAAAACAGGAACATCACGAAGGAAGGCGCGGACAGTGGCTAC
TACGTATGGTGCATTCCTCTGCCGGAAGGCGTTAGACCGTGGCAAACAGACAGG
AGACATAGTTTTGAATGAAGATAGAGC
>index191_0
TTTCTGTTGGTGCTGATATTGCTAACTCGTTAATTCCGTTAAACCACGTCCGGC
CTGTCAGCTCTCTTCCAAACAGGATCCTCAACTACGAGTCCCAAAGGCCAGAGC
TTAAGTGACATTAGGTGCAAACGCGGGATCAGTCAAGGCTGAGACTCCAGTATC
TCGCTCAAATGCCAATACGCAGGTGAAGGCGCTACAATCCACAGTTGGGTCAGG
AGAGGGCGGATTTAGCGAAGATAGAGC
>index192_0
TTTCTGTTGGTGCTGATATTGCTAATACCAGACCGGAGGAAGGCGAAGAAAGAA
AGGTCAGCTGATGCTAGCAGCAAAACGTCCGTACGCAACTGTTAAGGCTTATGC
GGACGCCGTCTCGCGAGTACCTTAATAAGGGGAGAAGGCGGCTTGACAGTTCTC
AGTCGGCACGAGACGAGATGAAGCAAAGGCTTTTAGGGATATCTAAACACTATC
GCCTCGACAGGCTCGTGAAGATAGAGC
>index193_0
TTTCTGTTGGTGCTGATATTGCTAATCTTGTTCAGCTGGAAAGACGCCTCCTAC
AACTCAGCTGACTGCAAAGATGCACACACTGCATGATTTGTGCAAGGCACTACT
AGTGCTCGCATCTTGGATTCAACCCTATCTGCCAAAGGCGCGATCGGACGCGTT
CGTTGAGAATCTTCCCCGTGGTGTAAAGGCGTAGGGGTCAACCTCGCTGCGCTG
CACGATCACCCTAACGGAAGATAGAGC
>index194_0
TTTCTGTTGGTGCTGATATTGCTAATTTAACATCGGGATTAACTCACCCACCAG
ATCTCACAGTCGTTTGTCCTCACGAACAAGCGATGGTATTACAAAGGCCCAACT
TCACTGCCGCAGACTATACTTTCAGAACTAGAGGAAGGCTACGGCCGTTGCTCA
TGGCTGCTGAAGGCAATCTAGAATGAAGGCGCCCGTTGCGCGTGTATAGAGCGC
ACCGTCTCTTATCCATGAAGATAGAGC
>index195_0
TTTCTGTTGGTGCTGATATTGCTACCGACGAAAGGACTACAAACGTATTATCCA
CTCTCTGCACAACAGCGTGACAGGAGAGTTGCTGCTTGACCACAAGGCATAGGT
GTATCAAGGACACAGGGTTACACGAAGGCGCCGGAAGGCCTCTTCGTGAGTGGA
GATACAGGTGTGTGGGGTCAGATTGAAGGCGCAGTACGGAAAGATCGGGCATAG
TCCTCCTAGCCTTCGCGAAGATAGAGC
>index196_0
TTTCTGTTGGTGCTGATATTGCTACCGACGAAAGGACTACTCCCACTTACAATG
ATCTCACCTTGCTAACTGAGTCCGCATGCCTACCGTAAGCTTAAAGGCAAACTT
CCACGAGGACCCTAGCTATCAGGATGGGCGTGGCAAGGCGATGTTGCAGTACAA
AACAGTTGCTGAAAGTGTAACCACCAAGGCTTAAACATAAAACAAGATGTCCTA
TTCGTATAGTGACCTTGAAGATAGAGC
>index197_0
TTTCTGTTGGTGCTGATATTGCTAGGCGAAGAAAGAAAGGAAATTTCATCCGCA
AAGTCACCTTCCTGCTTGGAGATGTGCACCATAATACTAGAGCAAGGCCTGTTC
CGAGTGGCATTCGCAAGTGCCTTGGAGTCTCCTGAAGGCCCCGTTTGTTCAGCC
CAAACGAGCGGTACTTGTTAGAACCAAGGCAACTGCCAATCGGACGAGGCCGCC
GTGAGTGTAGGACCTCGAAGATAGAGC

>index198_0
TTTCTGTTGGTGCTGATATTGCTAGGCGAAGAAAGAAAGGACCGACGAAACTTT
GTTTCCCTTACTGTTCCGGAGAAACTTCGGCCTAAGTCGCACCAAGGCGATGGA
AAGCCATACATCGCTCGCATATCCGCTCATAGCAAAGGCAGAAGGTAGACGTAT
GAGCAGCCGCCCGTTCTCACCATCTAAGGCGCGTTGAGTCCGGCTTACAAGCCC
AACCTATCCCCACCAAGAAGATAGAGC
>index199_0
TTTCTGTTGGTGCTGATATTGCTAGGCGAAGAAAGAAAGGCAGGCGTTGGGTCT
GAATCTACCACGACTCTTCAGCGAGACCTATTCAAGTGACCGTAAGGCACTATA
GCTAGCGCGTCTGGAATTCCGGTTCCCTCTTACAAAGGCCAGCGCCAAAGGTAG
GACCCTCAGTAAGTAGATAATCACAAAGGCACAGTGGGAGAGCAGATGTTTGGG
CCACTGACGCGCACATGAAGATAGAGC
>index200_0
TTTCTGTTGGTGCTGATATTGCTAGGCGAAGAAAGAAAGGAAAGTTGCAAGGAA
AGATCTGCTTGGAAAGTCCCGAATACTGAACACATCGGTTACCAAGGCTTCTTG
GTGGGTAGTCTATGGAACTATCACACGTCTAAGGAAGGCCATCCGGTATCAGTT
ACTCTCGCGGACACTAGATATATATAAGGCGCCTGCGGGCTTATCCTTGAGCCA
AGAGCTGCCACATGGAGAAGATAGAGC
>index201_0
TTTCTGTTGGTGCTGATATTGCTAGGCGAAGAAAGAAAGGATCCTCAGTTGGGT
ATCTCACCACTGTGGTGAATTGCGCTGATCGGTGGAAGCAAGAAAGGCAACCAC
TTGCAGAGGAGAAAATCCTGCCTGACATGGTGATAAGGCGCCGTGGCAGAAAAG
TTTAATCTTGACTGCCCGGGACGTGAAGGCGACAGAGTCGTGATCAAGTATGGG
GAACGACCGCTGGTACGAAGATAGAGC
>index202_0
TTTCTGTTGGTGCTGATATTGCTAGGCGAAGAAAGAAAGGCAGGCGTTTCCGCA
AAGTCTACGTGGTACGTAACCTACTACGTCTCAAGTTGACGACAAGGCGGTTTT
CATGTTCTCTGGCAAAGTTCTGTGTCTACTGACAAAGGCTGACCTAGACCATTC
GGGCTGGGTGGGGCGCGGAATGATAAAGGCCCTGACCCTGGTCAGCTTTGTCTA
GTGCGACGCCCAGTTTGAAGATAGAGC
>index203_0
TTTCTGTTGGTGCTGATATTGCTAGGCGAAGAAAGAAAGGAGGGAGGAAACAGA
TCATCTACCAGGTAACACTGTCGAAGACGAATGTCCGTGAATGAAGGCCCCGGC
CTCTACACATACCTATGAGAAACTGAAGGAATGTAAGGCGGTGGTTCTCAGTCG
GGCGTTTTCGAGCGCGTATTAGTGGAAGGCTTTCCACTTATCTAAGCGGCTACG
CCAGGTCCTTCAAGACGAAGATAGAGC

>index204_0
TTTCTGTTGGTGCTGATATTGCTAGGCGAAGAAAGAAAGGTCGATGGATCACCC
ACGTCTGCACTGTAAGCTAGACATGTGTGACGTGATCCAGAACAAGGCACAGTG
TAACCGTGGCTTAGGCTCATGGCCGGGGTTGGCCAAGGCTCTCTAAGCGCCAGT
AAATTGCACTAAGGCATAGACTGGCAAGGCCCACCCAGAGGGAATACGCTTCTA
GAGTATGTACGTTATCGAAGATAGAGC
>index205_0
TTTCTGTTGGTGCTGATATTGCTAGGCGAAGAAAGAAAGGCCCTAAAGAAAGTT
CAATCAGCTACTCCATAGACTCCGCACTGGTGAAATAGAGAGAAAGGCCGATCA
TCCTCGATGATTTTACGAGTGAGGTTCTGGCTTTAAGGCCGGGATCGCAAACGT
CCGTCTCTAAAGTTTGGCACGAGGCAAGGCGAGCATATACCAAAGTTACTTGTT
CATCAGGATGGGTACAGAAGATAGAGC
>index206_0
TTTCTGTTGGTGCTGATATTGCTAGGCGAAGAAAGAAAGGAAAGAAAGGAGGAA
AGATCATGGAGTGATCAGCGGTGTTTCTGCCCAGGAAGACCAAAAGGCCAGCTC
TCGAAGGGATAAGCGAATCCCGGAGCATACTCCGAAGGCGATGTATGCCAGGTT
AGTATAATAATCGGTAAGTACCAGAAAGGCTCAGCACACCATAAGAACAACATA
CAAATCGAACGGGGCCGAAGATAGAGC
>index207_0
TTTCTGTTGGTGCTGATATTGCTAGTCGCTCACTCCATCGACTGTTACCAACTA
CTCTCATGGTGTGAAGAGTCCAAGAGAAGGTTGTTCCCGACGAAAGGCACTTCG
CGAATATAACCCTCTATAGTAGCATAGAGACCGTAAGGCTGGTACTGTAGCGAC
TTTATACTGTCTCTGTACAGGCACAAAGGCCGTGTAGGATTGCATCTACATACC
AGACGCACCAGAGCGGGAAGATAGAGC
>index208_0
TTTCTGTTGGTGCTGATATTGCTAGTCGCTCACTCCATCGATTTCAAAGGTTTG
GTCTCACACTCGTTCAAGGGACGTCTGACAAGATAGTGGTAGTAAGGCTACCCC
TAGGGTCACCCAGGAGGTTAGCGGTATCTCTTGTAAGGCGCCCCGATCTAATAG
GTCCGGACGCCGACTACGGTGTCTGAAGGCAGCAGTCTTGTTCGAGACTCGTGA
TGGCGAAAATCGTCGCGAAGATAGAGC
>index209_0
TTTCTGTTGGTGCTGATATTGCTATCGTCATTCCCAAACCACTGTTACCAATTT
GTCTCTACGGATCGATAGGGGTTTTGCTCCTTGTGGTTAAGGGAAGGCCACAAG
ATGTACTGTATGCTCGTTATCTCGCTGCAAACCCAAGGCGGAGGGTCGACTGTA
CAGAACAAAGAGACCTTTCTTTTATAAGGCCCGTGGGAACTGTTTGGGATCAAG
GTAGGATGTTCTGTACGAAGATAGAGC

>index210_0
TTTCTGTTGGTGCTGATATTGCTATCGTCATTCCCAAACCACCATCTTTCCCTA
AAGTCATGGGATCGACGTTGTGTCACGCTAGCTCATTTTATGAAAGGCTCACTG
CGGCCAGACGCATTGTCAAATCCGTACTCTGAAAAAGGCCTGCAATGTGAATTG
CCATCCAACGCCTTAAGGTGCGCGAAAGGCTCGCGTGTGTCGCGAACTACCTCG
AGTTACATCCTTTGCTGAAGATAGAGC
>index211_0
TTTCTGTTGGTGCTGATATTGCTCAGGCGTTTCCGCAAAGACCATCTTTACTGT
TTCTCAGCTGATCCATTGTGACGGCGATTTTCAGATCGGTTTCAAGGCGGCTTA
CTTTTACCAACTGGAAACTCGACAGACCCAAACCAAGGCTCTAGACTACTCCTC
CCGCAAGTCCATTTCACGACCGCAGAAGGCACATTGGGGAGCCCCGGGTGACAC
CTTATGTTATATTTAAGAAGATAGAGC
>index212_0
TTTCTGTTGGTGCTGATATTGCTCAGGCGTTTCCGCAAAGAAAGAAACAACTCA
AAGTCTGCATCGTAGTTCAGCCTATCACAACCCCTTTTGATCCAAGGCCCACCC
GCGAATCCTCACTTTGTTGCCGCGGACATGTACAAAGGCTTTTATGCGCTGGCT
AATCTTATAGGTACAGTTTCCTGAGAAGGCTGCACGGGAACTATCAACGTAGAG
AATACGAACACGCGGGGAAGATAGAGC
>index213_0
TTTCTGTTGGTGCTGATATTGCTCAGGCGTTTCCGCAAAGAAAGAAAGGAAAGT
TATTCAACGGAAGATCTCACGTGACAAAGCTGAAGAGATCACCAAGGCGCGCCG
GTATTTTCAAGCAGCCAACCGTTGCAGAATCCATAAGGCAATTGTTGATGCGCC
TATCCAGACGAAAAGCTGTGCAACAAAGGCAATATGTGCCATGGTCTCAAACAT
GATTTGCCGTTGTATCGAAGATAGAGC
>index214_0
TTTCTGTTGGTGCTGATATTGCTCAGGCGTTTCCGCAAAGAAAGGATCACGGAC
TTTTCTGCTTCACATCCAGAGCAAACAGTAGACCCTACCACGAAAGGCCTTACA
CAAAGTTTCGGTCACCACGCGGGTATCTTGGTAGAAGGCTGTCTTGTTATATAT
TCCATACCAGATGCTAACCCAGTTAAAGGCACCATGGGAAACCGGTTGGCGCGG
TGACTCCTCTGTTCTAGAAGATAGAGC
>index215_0
TTTCTGTTGGTGCTGATATTGCTCAGGCGTTTCCGCAAAGCACCCACGATTGCG
GATTCTGCACAAGGACGTAGAGGCAAGATTCTGGCTACATATAAAGGCGGATAG
ACGTGCGCCGTATGTGTAGTGTGCGATACCGATGAAGGCGCTTTGACCTAGGGA
TTGTACCTCGACCGTGTGCAAGCCGAAGGCTAAAACTTGGCATACTGCCCAGTC
CCCTTTTAGCTATCTCGAAGATAGAGC
>index216_0
TTTCTGTTGGTGCTGATATTGCTGGACATTTACGGGTTTCAATGGAGGAACTAG
TTCTCGTACCATGAGGACCAAGTGTGACTTGCTGTGCTCTCGTAAGGCACACGC
AGGAGAGGAACAGCCTAATAATGGGGCGTCTTTTAAGGCGCGACGATATGATGG
ACTAGGAGTGTAAAAGAGCCACGAAAAGGCTTCGAATGATGCCTGGCTAGAGTA
CTAGTTTTAACCTCATGAAGATAGAGC
>index217_0
TTTCTGTTGGTGCTGATATTGCTGGACATTTACGGGTTTCAACCCTAAAGGGAA
GATTCATGGTCGTCTGTGAACTCGTCAACCCACTTCTACAGAAAAGGCAACCCA
CTGTGAAAATGTGCGGCATCGGTGCACGCCCGCTAAGGCAGGGGTACTGAAGCA
CTGACTTTGTTTACAATCAGTGCCAAAGGCCGGACAAACAGTCTACATTAAACC
GGACGGCCGCCCTACAGAAGATAGAGC
>index218_0
TTTCTGTTGGTGCTGATATTGCTGGACATTTACGGGTTTCACTCAGTCAAATCA
AACTCCCTTACTCTACTCAGACAAGACACACTCCCATCATCCAAAGGCGGGCTG
GTCGCATAGAAGTTGGCTACAGGGATTCTGCAACAAGGCCCTTATCAACGGCAG
TCGGCTTGGTGGCCGCCAGTGTTATAAGGCTATGGATACCACGATGAGTGAATT
TGCGTAAATATTAATCGAAGATAGAGC
>index219_0
TTTCTGTTGGTGCTGATATTGCTGGACATTTACGGGTTTCAAACTTCCCGCTTT
CCATCAGCTGTGTTGGTCTCCAAACAATGACAGTTGTGGATGGAAGGCTTGTCC
CGGATATACCGGTAGGCTATCTGCCTACGAATCTAAGGCGCGCCGCACAGTCAG
ACCGGGAATGCGTTTACCTACTAGCAAGGCGGAACCAATAACTGCTAGGTAGAC
ATTCTCTGGGTAGTTTGAAGATAGAGC
>index220_0
TTTCTGTTGGTGCTGATATTGCTTCAATAACCACCCACTCAATGCCTTGACGGA
TGATCAGCTAGGAGTACCGTTGGCAAAAGGTAAATGCGATGCGAAGGCTAGGTG
GCATCTTACGTAGTGGGACTACCGCGTAAATAGTAAGGCAGCTGCTACTCCAAC
TGCACCAGATCCGTCCGAAGTAACAAAGGCACGATGTTAAAGTCAATTCGCTTA
CCTTATTAAGATAAGCGAAGATAGAGC
>index221_0
TTTCTGTTGGTGCTGATATTGCTTCAATAACCACCCACTCACGGATGACATTGC
ACTTCAGCAGTACGATAGAGCGGCATCATCATGAAGGCCAGGAAAGGCGGAGCA
CACTAGTCTGCAAGCTGCCAGTATTGGTACCTTAAAGGCTGGTTGCCGCGTCGT
AGAGATTATTCTACGTTGAGTTGAGAAGGCTGTTCGCAGCAGTTATTGATGAAG
GACGATAACCCTTTCCGAAGATAGAGC

```
>index222_0
TTTCTGTTGGTGCTGATATTGCTTCCTACTAAGGCGAAAGGGACATTTACAATG
ATCTCAGCACTACGTCCAACAACGGTCTTGATGTATGCATGACAAGGCAGCTAC
GTAACCGTTTAGGCTAACCTCAGGGCTTGTCTCAAAGGCACGTTACTGCGACAG
AACATGTGAAGCTGGGATCTTACAGAAGGCAATAATCTCACTGGCAGCCGGCCG
TAATTAAAATTCAATCGAAGATAGAGC
>index223_0
TTTCTGTTGGTGCTGATATTGCTTCCTACTAAGGCGAAAGAAACATAACAAGGA
CGTTCCAAGGTGACCTGTCATCTACACGGGTGACAGCTTTTCCAAGGCGTAGGG
GTTTCATTCGAAAATCCAGTTAGTCCTGCATGGAAAGGCCCTTACGAGTGCTGT
TCGTTATTTGCAGCCCTAAACAATTAAGGCGCTTCGCGTGCCAGATTATAGTAC
GTCTTGTAGTGCCCTGGAAGATAGAGC
>index224_0
TTTCTGTTGGTGCTGATATTGCTTCCTACTAAGGCGAAAGAAAGAAAGGACCGA
AAGTCAGTGAGTGGAGGAAGTTTCGCTTGGAGACTGATCCCTCAAGGCCCAAAA
GCACACGAGTATAGCGCCCGGTCTTGAGTAGTTCAAGGCATACACAGGGAGAGC
TTCCTATCATTACGGGCCTGATAACAAGGCTCAGACCTGGGCGTATTGGCGGTC
CCCGGCCGTGAGTAATGAAGATAGAGC
>index225_0
TTTCTGTTGGTGCTGATATTGCTAAACATAACAAGGACGTTCCGCCTCAAACGT
TTATCACCTCAAGTTCTCTCCGATAGGCCTTGCAGACTACGTGAAGGCGGGTGT
TATAAACGCCTTATACACCTAACGCCCCAGACCTAAGGCACTTGCAATACTGAG
CTTGGTCTGGCCGGCGTGAGTATTAAAGGCACTAGGCCGGGGACGTTATTTTAC
AGTAAAAGCGCCCATGGAAGATAGAGC
>index226_0
TTTCTGTTGGTGCTGATATTGCTAAAGAAACAACTCAAAGAAGGGCGTCACCTC
ACCTCTGCTTGGTCACAGAGGAATAGAAGCTCACTGTGGCCTTAAGGCCATGAT
TGATTAGGGCGTGGTACGGCGTGAATTGGAATTTAAGGCGAACCTTATACGGTT
ACGCTTTGTAAGCATTGGCACTCGAAAGGCTCATCATCTAAAGGACATGGATGA
GTCGGGAAGGGATTACGAAGATAGAGC
>index227_0
TTTCTGTTGGTGCTGATATTGCTAAATTAAAGAAAGAAACTGTCGACGGACCGT
CCATCAACCAGAGGCAATCCACCCTTACGCTAGTTCACGTTCTAAGGCGTGGGG
AAGTGGATACACGCGCCAGACTGAGGACACACATAAGGCGAGCTGAGAGAGGTC
CTGTATCAACTAGTGTGGGTCCCTCAAGGCGCACAGTGTCTTTGCACAGACCGC
TTATCGGTTCTGTTGCGAAGATAGAGC
>index228_0
TTTCTGTTGGTGCTGATATTGCTAACAGATCAAGCGACTTACCTCACCAAAGAA
ACATCCACTTCCTATCGCGTCAACCTAATGAACGAACAAGACGAAGGCGATCCT
GGACGACTACGCAGGTGGATATCTTAATTGAATAAAGGCCTAGCTAGTGACAAG
AGTCGGAAACCGGGGCTGAGGCGTTAAGGCACCTGGTGAGTTTAGTAGCACGCG
ACCGGGAACAGACTGCGAAGATAGAGC
>index229_0
TTTCTGTTGGTGCTGATATTGCTAACAGATCAAGCGACTTTCCGGAGGAAACTC
AACTCTACCCAAGTCTTGAGGGCAGGACAGTTCCCAAGCTACCAAGGCGTTTTA
AAGATTTGAGTTGTGCCTCAGGTTTAACAGAACCAAGGCCGCCACCACTGGCCC
CTGAACCCGATGAAACGGACGGTAGAAGGCATGTTCCCAAGAATATAGCGTCAA
AAGGTACGGCCTCCCTGAAGATAGAGC
>index230_0
TTTCTGTTGGTGCTGATATTGCTAACTCACCCAAACCACCTGGGTATCAAACAA
ACATCAGGTGAGACTATGTCTCAACTCTCTGTCGGCGACTCTGAAGGCTAAAGG
ACGATGCACTCAGTCATTGCCGGCTGAGGTATTTAAGGCTGACAGGCAGCGTCG
TAATACGCCGACCTAATTTGCCACGAAGGCATCGGCCTGTCCCGCGCTGTTCGT
GATAAGAGTGAACAGTGAAGATAGAGC
>index231_0
TTTCTGTTGGTGCTGATATTGCTAACTCACCCAAACCACCACCATCTTTATGGA
TTGTCACCTTCCTGCTCACAAAAGAAGCCCCTCCATTTATTGCAAGGCGTCATT
TCCCTGAATCACGACTTTACGGATACTTGCATGAAAGGCTCTGGAACTGGGTTT
TCGCTAGGCGGGTAGAACCGCCGGAAAGGCATTAATGCCCGCCTGTTAGTCTCC
CTGTGAGGCTCCCCTTGAAGATAGAGC
>index232_0
TTTCTGTTGGTGCTGATATTGCTAACTCACCCAAACCACCACCACTTGCGTTCT
ATCTCCCTTAGCATCATCCATGTCTGTCGCCCAGTGTGCTGTTAAGGCCGGTAA
GCAGCAGCACACTCGAGTCATGGATCATAACCCCAAGGCAGTCTTCGTGGTGTA
CTACTTCAGGGTGTGCAACAACAAGAAGGCATTGAATATAAGGATTCGACAGAA
ATCGCGTTAATAACATGAAGATAGAGC
>index233_0
TTTCTGTTGGTGCTGATATTGCTAAGGGCGTCACCAATCGAAAGTTTCATTTCA
AAGTCCCTAACGAGATTCGTAGGCTGAGCTCCTAGAGCGACACAAGGCAACGGT
ATTGTGAAGCATCCTGCCCTACCATCTGATACTTAAGGCTCGTGCTTTACGAAG
AACCCAGCCACAAAGTCTCCGCAAGAAGGCCGACCATGTTAATAGACCAGCTGA
TCCTGCATTGGCGCGTGAAGATAGAGC
```

>index234_0
TTTCTGTTGGTGCTGATATTGCTAAGGGCGTCACCAATCGACCTCACCAAAGAA
ACATCACACGTGTCCTCTTGTCTCAATCTCACGGCGGACAAAAAGGCCACCCT
AAGGACAAAGTTACCAGCACTAAGGATACCCCAAAAGGCGTCATTAGTGACTAG
CTCGGTGTGCATGCTGTCCTAATACAAGGCGCCATTGCACTCTCTGCTGCCGAT
TAGTGGATGTACGTCTGAAGATAGAGC
>index235_0
TTTCTGTTGGTGCTGATATTGCTAATCAATCAGCTTTCCACCCGGCATCACCAC
GTCTCACCTCTGACCTAAGCACTATCTACGGACTTGGCGCAATAAGGCCAGGGT
ATCGGGGATGCCAATAACGCGAGCCGTCATACAGAAGGCGACTCAGCATAGAGA
TCGCAGAGAAGTAATGGGGCAACCAAAGGCAGCTATAGCCAGGTGTATCAAAGC
GACGGGGAGCGTCGCGGAAGATAGAGC
>index236_0
TTTCTGTTGGTGCTGATATTGCTACAGACTCAAATGGAGGAGGCGAAAGCGGAC
TTTTCAGGAACAGGCAGAGATCCCTAAATAGGAGAGTCTATCGAAGGCTCAATC
GGTTCGGCGCCAGACTGACTGAGTCGGGGTCACTAAGGCAAATATATAAGTTGA
TCACGTATGGTCCCGACCCATCCGTAAGGCTCACGGTCACGGAGTTTGGATCCG
TTAGGTGGGGCCATCGGAAGATAGAGC
>index237_0
TTTCTGTTGGTGCTGATATTGCTAGGTGCGTCAGTTTACGACTGTTACCTCAGA
CTCTCAGCTAGGAGTCATCCCAACGCTAAACCAGGATGCCGTCAAGGCATCCCA
AAGGTTGTGGCTTCTAAATCAACAAAAGATTGGCAAGGCTAAGGCAAAGCTCCC
GTCGGAGCGATATAGTACTGCCCCGAAGGCCCAATCATACTACGGGCCAAATGG
TGAGAAGAGCTGCTTTGAAGATAGAGC
>index238_0
TTTCTGTTGGTGCTGATATTGCTAGGTGCGTCAGTTTACGAAAGAAAGGAGGAA
AGATCAACCCATGAGGCTGAACGGACTTCTGTTAAGTGCACTTAAGGCAAGGCG
GTCGTCACAATATGTCAAAGTGCTCATAAACCAGAAGGCAACACATGTGGGTTT
CAGGATCACAGTGTTCGCGCAACGAAAGGCGAGTCAGTGCTTCACGTGTTATTG
CGGCAAACCTTTTGTTGAAGATAGAGC
>index239_0
TTTCTGTTGGTGCTGATATTGCTATCGGACTTCCGGCCTGGGACATTTACGGGT
TTCTCATGGGAGATTGACTCTCGTGAGTTACTCTTGGCACTTCAAGGCCGCTCT
AAATTCGCATGCGAGCACATATACCCATCTAGCAAAGGCATTCAGGTGTAGGTG
CCGGCTTCTTTACGCCTGGCGTTATAAGGCCCTCTCTGTGTTTTAAGTTTGATT
ATCGGTGCATCCCTCTGAAGATAGAGC
>index240_0
TTTCTGTTGGTGCTGATATTGCTATTTCAAAGCACCCAAACAGGCGTTTCCGCA
AAGTCTGCAACAGGCTAGCTGTCGACTTCATCATAGATCATCTAAGGCTGCACC
GATACCGATGTCGCAAGTAAATTGCAAACTCGTCAAGGCAAAGTCGCAATGGGT
TAGGCCTTCTTTCACCTAAGTACGCAAGGCCTGCTCGTTAGCCCGTAGCACAAC
TCCGTAACGGTTGTAAGAAGATAGAGC
>index241_0
TTTCTGTTGGTGCTGATATTGCTATTTCAAAGCACCCAAATCCGCCTCAAACGT
TTATCACCTGTGTTCCAAACACCTGCGTATGCTAGGCGTGCATAAGGCGTTAAT
AGGGGTATGTTATCTGTATCTGCTCAAACCGTGAAAGGCGAGGGTTTCCGGCTT
TTAACATGTTACCAGGGGCCGTATTAAGGCTATGCTCAACAACCCATAAGCTAA
CGCATGTACCGGGTGGGAAGATAGAGC
>index242_0
TTTCTGTTGGTGCTGATATTGCTCCCACGTTCACCAACTAACTCAATAACGGGT
TTCTCACCTGTGACGATAACGGGTTACCTTGGGAACTATACCAAAGGCCCTTTT
GGAGAGGATTTAGGGTCCCTGTACAGTATCCACAAAGGCACGCAATCCAGATGG
TAGTTCCTCTGAACACGGGGTCAACAAGGCGATTAGTTTTCCATCAGAGGTTCA
CGCTTATCTGTCTAAAGAAGATAGAGC
>index243_0
TTTCTGTTGGTGCTGATATTGCTCCCACGTTCACCAACTAATTTCAAAGCACCC
AAATCTGCACAGTTCCAGACTGGGAACGTCTTGGGTTTTGCTGAAGGCCTTCGG
AGCATGGAACCCTCCTCTCTGTCAGGACACTAGAAAGGCAAATGTTTAAGTTCA
CGATATAGCGCCCCTACTAAGATGGAAGGCTCAGATTGCCTAGTCCAACCGATG
CGCTGCGGATCTACACGAAGATAGAGC
>index244_0
TTTCTGTTGGTGCTGATATTGCTCCCACGTTCACCAACTAAGGGAGGAACGTTT
CAGTCACCAGAGTGAGTACACGTTGGGGTCACACCAGTTAAGCAAGGCTGTTCT
GCTAAGGGGAACTAGGGGAGTATGTAGCCCGGAAAAGGCTCTCAGTTTCTTTCA
CGACGTCGTTGTACTGGGGTGTGTGAAGGCAGGCAGATATAAGCCGAAGTTTAG
CTACCGCTCTCGACGGGAAGATAGAGC
>index245_0
TTTCTGTTGGTGCTGATATTGCTCCCACGTTCACCAACTAAATGTCGTTGATGT
TATTCATGGCTACCAGGACTTGGATTAGATGCGTGTAGTCACTAAGGCCGCCCG
TGCATCGCTGTGTTTTGTACAATACGTGCACAATAAGGCTCAGTCAGGCGGGCT
TCCCGTTACATTGCATGGACCGGCGAAGGCTAGAGTCTATTCCTCCACGCTGAA
ATCTCCGGAGGCTGGAGAAGATAGAGC

>index246_0
TTTCTGTTGGTGCTGATATTGCTCCTCAGTACTCACCAACAAATCACCGAATCT
TCATCAGCTACGTGCATGGTCTAGTACGCAGACTTGCAGTTCCAAGGCCTATTG
ATCATACTCAAGTCTATGGTTGCACACCGCTACCAAGGCTGCCGGGGTTCTAGT
CCAACCATCCGATAGCCAACACTACAAGGCCTTCAAATTGGGTGCTGATGCCAC
AATCCACACACTGACGGAAGATAGAGC
>index247_0
TTTCTGTTGGTGCTGATATTGCTTCCAAATTTATCGTTCCACCACCCATTCGAT
GTCTCAACCACGTTACCAGTGTAGAAGTCACGCCGGCTCATTCAAGGCGCATGA
GTTTCCTTTGGAATTGGCTCATTCGGAAAATCGGAAGGCCTCGCCACCGTTTTC
AAGCTAACACTATATATTCGCTAGAAAGGCTGCCTTCAGCATTGCGCGATTCTG
AACACGCTTAGGAACCGAAGATAGAGC
>index248_0
TTTCTGTTGGTGCTGATATTGCTTCCTACAACAGGGAGGAATTGCATTACCCTC
CCATCTACCAGCTAGTTCTCTCGGTAGCAACACACTTCTGAGAAAGGCGTGCTT
GTGGAGAGGGACAAGTTAGCAACCTCCTCGGGTAAAGGCACAGCTCGGCTGGAT
ACCTTGAGATTCCAGCACAATGCGGAAGGCTTAGATGAATTGAGCCCACTTATC
CTTTGAACATAACTTAGAAGATAGAGC
>index249_0
TTTCTGTTGGTGCTGATATTGCTTCCTACAACAGGGAGGAATTTCAAAGATTGC
ACTTCAGCAGTCATCGACGATTAACCACTGTTCATGCCGGCTGAAGGCTTAGGG
CTACGTCGTTCCCTCCCATAAGTTTGCTATATCGAAGGCATCCATGTGCGAACA
ACCGACCAAGTGACTCAATTTTATCAAGGCCTGAAGTTGCGGACCAGCACAGCT
TATAGATACCCCGATCGAAGATAGAGC
>index250_0
TTTCTGTTGGTGCTGATATTGCTTCCTACAACAGGGAGGAACTCAGTCAAAGAA
GTCTCCACTGTTCCAAGCCTGTGGACTCATAGCTGCTGGACTTAAGGCTAGGGG
CACACGTGACAGTGCAATGTGAGAACCATTCGCTAAGGCGCTAGGTTGCCACTC
GCCTTAATCACTAGACTTATAACGAAAGGCTGGAACTCTATCAGCATTGTGGAG
GGACCTTCTGCATTTGGAAGATAGAGC
>index251_0
TTTCTGTTGGTGCTGATATTGCTCCGGCCTGAACTCACCCACCATCTTTGCTTT
CCATCACGATCAGCCACTAATCCGGTCAAAGCTGAGCCTATGCAAGGCTAAACG
CCTGGGCTATCCCCAGCTCTTTGCGCTTGCAAGAAAGGCTAACACGTCGTTCGC
CGAGCACTCGTAAAGATCGAAATCTAAGGCTGAGGGGTTTGACTATGCACCAGG
TAATAGACATTAGTGAGAAGATAGAGC
>index252_0
TTTCTGTTGGTGCTGATATTGCTCCGGCCTGAACTCACCCATTTCATTGACATT
TGATCGTTGCATGCTGGAGACACAATCTTGTCGCACGTTTGGCAAGGCGGAGGA
GTTCCTCGTCTTATAAAACCCCTGTAGTCAATTGAAGGCTTGGCGAGATGTAAC
ACAAATCTGTTGCGGACGGAATTGTAAGGCCAGACTAAAATACTAACTGATCCC
GTATGTGCACCCTTCTGAAGATAGAGC
>index253_0
TTTCTGTTGGTGCTGATATTGCTAAACCACCACCACGACTACCTAATCAATCAA
TCATCATGGAGCATCCAGCATGTACGTGTTACAACATGTGTCAAAGGCCAGTTG
AGCCCTGCCTACGCAGACAAGTGATAAAGTTGAGAAGGCGTAAGTTCTCAGGTG
CCGATAAGGAGAACTAATGTTAACGAAGGCCGCATCAGCTGAAATATCTGCCGG
AAGGGTACGGGTTAGGGAAGATAGAGC
>index254_0
TTTCTGTTGGTGCTGATATTGCTAAACCACCACCACGACTAAACCACCACCACG
ACTTCATGGTCAGGCATGTTGCCCATAATGTCGCCCCTACCTAAAGGCCAGCTC
GATGGGTCATGGAGGACTCTGTCTTCTGTGTAGCAAGGCAGCCTGGTTGGTAGT
CTGAGTCCTTTCGCGGTGGTTGGTTAAGGCTCATTTATTTAGCGCAAGATCGCC
TAGTATAGTTCGCCTAGAAGATAGAGC
>index255_0
TTTCTGTTGGTGCTGATATTGCTAAACCACCACCACGACTAATTCCGTTGTTCC
CACTCTGCATCTCCCTACATGCGCTGTAGCAGTACTCGGTTAAAAGGCCGGGAC
CAATAGGTTCCTTTCGCCTTCTCGCCTTGCTCTGAAGGCTTGGAGTGAAGTGGT
AGGGGCGGTTCGATCATTCGAAAACAAGGCCATCCCGTTGAGGAAGTTAATCAA
AAGGGTATAAGCATCGGAAGATAGAGC
>index256_0
TTTCTGTTGGTGCTGATATTGCTAAACCACCACCACGACTCCCTCGTTTAAACT
TTATCTGCACATGCGTAACCGAGCATATGATCAACAAGTCCGCAAGGCGATACT
CGTAGGATGCACCCGTGAGACCGAAGTAAACCACAAGGCTGTCCTGAAGGAACT
TTAGCCTGGGTATTGCGAGACGGTGAAGGCGATTCTTCAGCTTAAGTCATTAGT
CGAAGGCGACATTCTTGAAGATAGAGC
>index257_0
TTTCTGTTGGTGCTGATATTGCTAAAGTTTCCACCCACTCATCGGACTTCCGGC
CTGTCAGCATGACACCGTGACAATTCGAATGTCACTTGAGTCGAAGGCATTCTC
ACGTCGTTGAGTGAATAGTCCCCGCAACGGCCACAAGGCCGCAAATATAGGTCT
ATAAGTCCGGTCGGCCCCTAAACGCAAGGCGCACCCCACTTTACCCGATTGAGA
GCCCCATTCGGTTTACGAAGATAGAGC

```
>index258_0
TTTCTGTTGGTGCTGATATTGCTAACTCGTTCCCACGTTCATTGCATTACCCTC
CCATCACGTACAGCCTCAGAGTCTTACTGCCCTAACCTTATCGAAGGCACTAGC
TGCTACATTCTAGCAAAACGGCTGGGTAATGCTAAAGGCTGATAGTCAGGTAAC
CCGATGGTGAATGCACATAAGCTCGAAGGCTCGTGTATATCCAAACGCGGGACT
ATAGGCCTCGCAGGTAGAAGATAGAGC
>index259_0
TTTCTGTTGGTGCTGATATTGCTAACTCGTTCCCACGTTCATTTCAAAGATTGC
ACTTCAGCTGATCGAATGCGATTCGCGAATCGGGCTGTAGGTCAAGGCCAGTCA
AGACTTAACAAGTACCCCAGAGATACAGTATGGGAAGGCGTGAACTACGGCGAA
CTACGAGCAAGTTGTGCAGCGTTAGAAGGCGCGTAAGCATATATTGGAGGAGTC
GGACAGGAACGTCGTTGAAGATAGAGC
>index260_0
TTTCTGTTGGTGCTGATATTGCTAATGGAGGAACTAGTTCCCGGCCTGAAAGAA
ACATCAGCTTGGTGGAAAGAGGGCTTGGATCTCTAACGTTATGAAGGCCGTGCC
TACGGCGAAAACTTCGATAAGCATCGTTGTTCTCAAGGCTGATAGGCATTCCTT
ACCGATCTGTCTGGTTCTGGTAAGTAAGGCCATTGACTGTACTTGTGAGATACA
GCTTATCCACGGGCCAGAAGATAGAGC
>index261_0
TTTCTGTTGGTGCTGATATTGCTAATTTAACATTTCATTGATCCCACGGACCTC
ACCTCACAGGACTCTAGGCTTCTATAGTGCGGGCGAGATATTAAAGGCTCAGCA
CTAGCATACCGGGACACATTCAAAGTTGGGGAGGAAGGCCGTGTTGATGACGTC
GAGAGAAACATAGCTCGTATCTGAGAAGGCGCACTCGAGTCTCTAGAAGCCCAA
AATCCTCTCGCGTCTAGAAGATAGAGC
>index262_0
TTTCTGTTGGTGCTGATATTGCTACCACGGAAACAACGTTAAACAGCCACGGAC
TTTTCGCTTGACAGCTGAGAGAAGCTATGTCGAATCTTGGCGGAAGGCCCACAA
TTTTGCACTGAACATGGGCCTTTGATAAATTACGAAGGCGATATGACGCCGCCT
TAGATCTTCTTAGTGGACTCTATAGAAGGCTGATGGTCTAGCGTTAACAGCTTA
ACAGGTGCCACACTTAGAAGATAGAGC
>index263_0
TTTCTGTTGGTGCTGATATTGCTACCTCACCAAAGAAACAAACCTTTCATGGCC
GTCTCCAACGTGACTGGAAGCACTTTAGCCTCTTAACACGTTCAAGGCATAGCA
TCACCGTCGCACCTTTCGGGCCGTGAACGATATAAAGGCACACGCGCCGAAATG
GGCACGCCCCTCATGCCATATCGCCAAGGCCTAACTCGTGCAAATGCACAGTCT
TCAGGCAGCTCCAGGCGAAGATAGAGC
```

```
>index264_0
TTTCTGTTGGTGCTGATATTGCTACCTCACCAAAGAAACAACTCCGTGAAAGAA
AGGTCTGCACACACCACTGACAATTAGACAGACCCTGGACCCGAAGGCAAGGCA
GTGTGGTCACTAAACTGTGTTCAGGTCATAATTAAAGGCGAGCAATAGCGTCAA
CGCCACACTAAGCGCAGAAGATAGTAAGGCAAAGAGTCTTGCTAATGCATTATG
CTATGATTGGTACAGGGAAGATAGAGC
>index265_0
TTTCTGTTGGTGCTGATATTGCTAGGCGAAGAATCTTTACAACCTTTCATGGCC
GTCTCCACAAGAGCAGATACCACTGTACCATGTGCCCTCAAATAAGGCCACCGT
CACTACTCGTATAACGCATAAACTAAGACGGGGTAAGGCACCATCACGCTTTTC
TTCCTGCACGATGTCTGTCGCCCAGAAGGCGCGACGCCTCAGAGTTTTAGGAAA
TGTGAAAGTAACTAGGGAAGATAGAGC
>index266_0
TTTCTGTTGGTGCTGATATTGCTAGGCGAAGAATCTTTACAAAGATCACCCACG
TTCTCATGCGTACCCTGTTAGTCGTAAACTGCACTAAGTCCAGAAGGCCGAAGT
TCTACTTAGACCCACCCGAGAGAGGCTGACTAAAAAGGCTTCTTTAGATATCTG
GTTCTGGTTGCTTCTGATAGTCCCAAAGGCGCTGCGTTACGACTCGGCCTCTAC
CAGTCTGACGCGCAGTGAAGATAGAGC
>index267_0
TTTCTGTTGGTGCTGATATTGCTAGTCGCTCAGGGAAGATAAAGAAAGGAGGAA
AGATCCCTTAGGACGTCTCAGTAAGTGGCGTTTTAAGTGGTCAAAGGCCGCTAA
TCAGCAATTAGAGTTGACCTTCTGGCTATACAGAAAGGCCGTCCCAGTGGGATG
CCCTCACTGCCCAAAAGCAAGTGGAAAGGCGTCATTCATATGCACCAATGGCAG
ATTTGACAAGTCAAATGAAGATAGAGC
>index268_0
TTTCTGTTGGTGCTGATATTGCTAGTCGCTCAGGGAAGATAAAGAAACATTAGA
CGTTCATGGCTTGCGAGTTGAACACGTCTTAGCTGTCGATACTAAGGCTACTGG
TGCGTGTAAGTGTGCTGCTATCTCGGGGCCCTCCAAGGCGGACGTTCAATCCCC
ACAGCATCTCTCGCATAAAACTAACAAGGCTAAATGAGCCCGGACAAATACGGT
TGACGAAAGTCAGGGAGAAGATAGAGC
>index269_0
TTTCTGTTGGTGCTGATATTGCTATCGTTCCAAACAGCCACCCATTCCCACCTA
ATCTCTGCACAGATCGAACCAAACCGAAGCCAAGAGTTGCCGGAAGGCCAACGT
GAAGTACTATTCATTGAGGACGGGGACTCACATCAAGGCAGTATGCCGTACAAA
GTGACATAAACGTTTCCATGAGGCCAAGGCACTTTTGTGTCGTCTTCTTGGCCA
ATTTCACGCTATAACCGAAGATAGAGC
```

>index270_0
TTTCTGTTGGTGCTGATATTGCTCAGGCGTTTCGATGGATAGGCGAAGAAAGAA
AGGTCACAGCTGACACGTCAGTCAACCTTAACGTGCCAGTTCTAAGGCGTGGAC
CTACTACAACCACCCCAAAATCTAATCGTGCCGGAAGGCTTACTTATAGAATTC
TACGACTAACGATAGTGGTCGAACTAAGGCCCTGACCCGGGTGAATTAGCTGTC
AGTCTGGCGGGACTTGGAAGATAGAGC
>index271_0
TTTCTGTTGGTGCTGATATTGCTCAGGCGTTTCGATGGATAAAGAAAGGATCGT
CATTCACACTCTGCTGGTACCTGAACAAGTTACCAGCAGATAAAAGGCGCTTTC
TCGTGCTGCGCGGTAGACTATAGCCTGCCGGGCAAAGGCGAGAATCCGTGGCAA
CAGGACCCTGAATCCCTACAGCCTAAAGGCGCCTGTGTTCCCATCAGGAAGTAT
CCTGAAAAGCACCGTGGAAGATAGAGC
>index272_0
TTTCTGTTGGTGCTGATATTGCTGGCGGAGTAAAGAAACAAATGTCGTTAACTA
CTCTCTGCTTGGAAGTCGCAGTATAAGTCAGTGCCCCAGGTGCAAGGCTCTACT
ATCATCCTTGGTATCCATCAGGACTCGGATAACTAAGGCGGGCTACCAAGTATG
GATCAGGTAATCGAGGTCGAAGGACAAGGCTTTTATATGAGCAGACTTACAGCG
ACAGCCGGCCCCTCACGAAGATAGAGC
>index273_0
TTTCTGTTGGTGCTGATATTGCTTCACCAACATTAGACGTTCAATAACCACCCA
CTCTCAGGACAGACGAGTCTTCCTCCATAGGTAGAATCTTACTAAGGCAGTCTG
GCAACGGTGGCGTGCTTCGACCACAAAATCTTGAAAGGCGTTTCTTAAAGCACT
AAAGGGGTGTTTGTATTCGGAATCCAAGGCTGATTGTGCAATGGCGCGGGTTCT
AGCCTGTTCTAGGGAGGAAGATAGAGC
>index274_0
TTTCTGTTGGTGCTGATATTGCTTCCCGTTTAAGGGCGTCAACATGTTATTTCA
AAGTCTACCTCAGGACGACTCACTATCCTACGGTTGTGCAAGCAAGGCGTTTTG
GTGACCCACCCACACTAAAACGCTGAGTGTATATAAGGCCGGCTAGGACCCGTA
CCGACCTTTCATAGGCCAACGTCCGAAGGCTTATAAGTCTCCTACACGGGGTCA
GGGCTCTGGCCGACCCGAAGATAGAGC
>index275_0
TTTCTGTTGGTGCTGATATTGCTTCCTACTAATTATGTTTACCCTCCCACCCTA
AAGTCATGGAGGTGAAAGTCTGGACAACCGAGGTCCTACGGTAAAGGCGAGGCT
ATGCTATTCAACGGAGACATCGTTGGTCTTGATAAAGGCACATTGGCGTTTTAA
AGCTATTGCCATAGGTCTTAGGTGAAAGGCGGGTGGCGCGAATGGCAAAACTAG
ACCGTTCGCTAGGGTCGAAGATAGAGC
>index276_0
TTTCTGTTGGTGCTGATATTGCTAAACGTCCAAAGTCTTTCCCACGTTCACCAA
CTATCAGCAGATCACGCTACAGCTCTCAAGTGACTGTCACACTAAGGCTTGCGA
CCTCGAGAATCCAACTGGATGCGAGTCAAAAGTAAAGGCGGTTAGACTCTTACG
GGCAACCGAGGGGAAGAACTGAGATAAGGCGGATGCTGGAGGTGAAAACTTCAC
CCATCTTCAGTTAAAAGAAGATAGAGC
>index277_0
TTTCTGTTGGTGCTGATATTGCTAAAGAAACAAGGTCCGCTCCTACAACAATCA
ACATCTACCAGCAGAAACACGAGTACCAGACGTACGCGGTTTGAAGGCGACGGT
GTTATAATTTCAGACCCAATTGAAGTGGGGCACCAAGGCCCAGTCTGAGCTGTC
TCTAGCCCAGCAACTATTAGAGAGGAAGGCCCTTATATTGAACTCCTGCGATTA
GGGGCTTGCAAAGCGCGAAGATAGAGC
>index278_0
TTTCTGTTGGTGCTGATATTGCTAAAGAAACAAGGTCCGCAACCTGGATGTTTG
GTCTCAGCTCTCTATCGAAGTCTCAGCATTGGCGTCGATGATCAAGGCTATCTT
TCTGAAGACCAAGTTCTCACCCTCAAGCGCTCTGAAGGCAGAGTGCTTGGACTG
GTAGTCATCCCACTGATTTGTCTGTAAGGCCCACCACCCAACATAAAATGCAGG
GGATTAGGGGCACGTAGAAGATAGAGC
>index279_0
TTTCTGTTGGTGCTGATATTGCTAACTCACCCAATCCGTTATTTCAAAGCACCC
AAATCCCATACTCGGAGTGTAGGATCGTAACGCCGTCTCTTAAAAGGCTTTCGG
CGTACATGGTCGCTTCATGTAATACGGCCTGTATAAGGCAAGCTGCTGCTAGGG
CCGAGTCCACTGGTGAACCTTAATAAAGGCTACATATGGTGATATGCGTTAGAT
CGACTGACCACCCGAGGAAGATAGAGC
>index280_0
TTTCTGTTGGTGCTGATATTGCTAAGGGCGTCACCTCACCAAAGAAAGGAACAT
ACCTCAACCAGGACTGGATTTCTAGGGACGAAACCTGGGCGATAAGGCCTACAT
ATCATGGGCAACCCCGACCCGTACGGATGTAATTAAGGCCGCAAAACCGTTTGA
AATGCGGATTCCGCCTCGTAGCGGCAAGGCCAGACGAGGGGAACACCGACGAGA
GTGTTCACATCCAAAGGAAGATAGAGC
>index281_0
TTTCTGTTGGTGCTGATATTGCTAAGGGCGTCACCTCACCTCCGCAAAGATGGG
TTTTCAGCTCTAGTTCCGTGGTAAATCCAGGTGGACACTAACGAAGGCGCCGAG
TCTTCCGGAAGGTATCTGTTAATGAATCCGCACGAAGGCCTGAGGAACTCTACC
CCATCCTTCCGTTAAAAGGCACTGTAAGGCAACGATTAGGGAGGCCGGTCGGTA
CATAAAGCGATATCACGAAGATAGAGC

>index282_0
TTTCTGTTGGTGCTGATATTGCTAATTCCGTTCACCCACGAGGGAGGAATTCTG
ATCTCATGGGACTGCAATTGACTAGGACCCCGTCAAGATTTGGAAGGCCGGATT
GTAAAGTGCAGATCCATCAATGAGGATAGAGCCGAAGGCCACAGAAATCGAGGT
GCGTCTATTCCCGCGTTGTACGTAGAAGGCTTTGACCACTAAGGAGTGTTTGGG
ATTAGTAAACGCGCCAGAAGATAGAGC
>index283_0
TTTCTGTTGGTGCTGATATTGCTAATTCCGTTCACCCACGCCCACCTCCCGGAC
TTTTCCCAACTTGTCGAGCACAGCAGTATGTCAGACGGCTCATAAGGCAGACAC
TTGGGGCGTCTCATACTAAGAATATCTAAAGCCCAAGGCCTTCCACGTTGAGAG
CATCCTTTTGTTAAGGGGTATTGCAAAGGCTGGATAGTCTCCATTCCGGGTTGG
TTAAGACGGGAACATCGAAGATAGAGC
>index284_0
TTTCTGTTGGTGCTGATATTGCTATTATGTTTACCACGGATCCTACTACCCACC
TCCTCAGGAAGCTATCTCCCACAAGGAGTGTGTCTGCAGAAGTAAGGCACACCT
TGGGAATAGTACGCGAGGGCCTAGTTTCATCTTGAAGGCAACGAAACGCCGCCG
AGTTGCCTCTGGACGCTTAAACACTAAGGCGGTCGACGGTGCCCGCTGTACTGA
GCTAAAATTAGACCTCGAAGATAGAGC
>index285_0
TTTCTGTTGGTGCTGATATTGCTATTTCAAAGCAGGCGTTAAACCACCAATGTC
GTTTCTACCTCGTCCAGTAACGAGAGTCGCAGTGAGTTTCCGCAAGGCGTAGGA
GTGTTATCTTCAAGCGTTTTGCCATTTTCCCCAGAAGGCAAATGGTCCGTGCAA
ACGTGAACTAGACACTAGGCCCACCAAGGCACATATCGTGCACATGATTCGTAA
ATTAGGCCTTTGTTGTGAAGATAGAGC
>index286_0
TTTCTGTTGGTGCTGATATTGCTCCCACGTTCATCGTCATGTTTGCTTAAGTGC
CGCTCCACTGTGAACTACTCTGGACCGTACCCTCTGACTCGAGAAGGCACATGA
GGGTCGTCAACCTCCTGAGTCTGATGTAATCCTAAAGGCGACATAGTGTTTTGA
CCGCATTCCCCGGGGCCCCGATCATAAGGCGTCCTGACGAAACCACGGATATCG
GCTAGGGGCAAACCGTGAAGATAGAGC
>index287_0
TTTCTGTTGGTGCTGATATTGCTGGGTAGGTTAAACGTTCAAAGGTAATACCAC
GGATCACCACTAGTTGAGAGACACGGCCAGTTGCCAAACAACGAAGGCGATGTC
ACCTTCTATCAGGTGATGCTCACTTTGTGGGAGCAAGGCCTTCGCAGCGTTGGT
GCGCCCTGTTTGCCACACATTCGCCAAGGCTATCCGTCTAGTTTTCCTAAGGCA
CCCAAAGGCAGCCTTTGAAGATAGAGC
>index288_0
TTTCTGTTGGTGCTGATATTGCTTCCAAATTTCCAGCGACATTATGTTTACCAC
GGATCTGCAGTTGTCCGAACCAGAAAACTGGATTCTGATCACTAAGGCTGACCA
CGTTAGCGTATCGTCAGCGCTGCACCTGCAGACAAAGGCTCACGTATTTGTAAT
TGAAGGAAATGTGTCTTGGTGGTTCAAGGCCCCTGGTTGCTTACCGCTAACCTT
CACAAAAGTCGTGCCGGAAGATAGAGC
>index289_0
TTTCTGTTGGTGCTGATATTGCTTCCTACAACTCCGCCTCACCATCTTTCCCTA
AAGTCCAAGGTTCGTCACAAACTTCCAGTCTACAGATCAGCGCAAGGCTGCGTG
GCCTACCAACAGCCGAAAGGAGTGGAAAAGGATAAAGGCTTATGATATGCTTAT
CCCGTAGTAAGTAGACCCACGCCTAAAGGCTAGGGACCCGGTGTCTAACCGAAA
AGGTCCCCACACGTTTGAAGATAGAGC
>index290_0
TTTCTGTTGGTGCTGATATTGCTTCCTACAACTCCGCCTCATTATGTTTGTTTG
CTTTCACAGCACTTGGTACCGACCACTTGGTCAGAGGTCAGAGAAGGCACTAAT
ATATACAAGACCTTGATGGATCTCTGCGAATAGCAAGGCGAATAGACACGTTTC
ACAGTACCCCTCCGTTCGGGCAAAAAAGGCCGTTTTGTCGTAACTCATCGAGAT
ATCCCGAGCTATAAATGAAGATAGAGC
>index291_0
TTTCTGTTGGTGCTGATATTGCTAATGGAGGAAACCACGTAACAGATCAAGCGA
CTTTCAGCTCACTTTCGAGCAGTGGCAGATCACAAAATAGCAGAAGGCCTAAAG
ACCTGTTCACGTCATATGGACTCCCAAGTCGGGTAAGGCGAAACCATGGGGTCA
AGATACGGGTGATATACCCGAGTGAAAGGCTTATGCACACTGCTACCACAACCA
CTCCCCACGCTGCACAGAAGATAGAGC
>index292_0
TTTCTGTTGGTGCTGATATTGCTAATGGAGGAAACCACGTAATGGAGGAACTAG
TTCTCACCATGTCTTCACTGGCACGGAGAGAGTCTAAACTGCTAAGGCATTAGG
TACTGTGCCGTCAATTCCCTGGCGGTAGATGTGGAAGGCCGGGTTGTGCATCGA
GATCGTCCCCGCAAATTACGGTTACAAGGCGACAGCGAAGCATGGCGGCGCTGA
CTGTCTACACATAGCGGAAGATAGAGC
>index293_0
TTTCTGTTGGTGCTGATATTGCTAATGGAGGAAACCACGTCCCACGTTCATCGT
CATTCCAACCTCACTGTTTGTGTTCGTGATCAGTTGCAGTGGTAAGGCTGGCCT
ATAGTCAAGGTACCGTGAACCGTCAAACCTTCACAAGGCTACTCACTCACTCGC
TTAAAGCGAGGCAAAACACCTAACGAAGGCCTTATTAGGATATATGCGGCCCCA
GGTATATCTGAATGGCGAAGATAGAGC

```
>index294_0
TTTCTGTTGGTGCTGATATTGCTAATGGAGGAAACCACGTACCACTTGCGTTCT
ATCTCCCAACTTCCTCGACAGAGATTGACCGCTGCTTCTTCGAAAGGCAGTGCT
AGATGTCAGCCACAGGCGGAACTACACCCGATATAAGGCCGCCAGACTAGAGAC
AGCATCGATTACATAACCGCCGAACAAGGCATGCGTCGGTACAGGAGGTTCTGA
TTTTAAACACTACTGCGAAGATAGAGC
>index295_0
TTTCTGTTGGTGCTGATATTGCTAATTTAACTCCGCAATCGTTTGCTTCCCACG
TTCTCTGCACTCTTTGAGGGGTTTCACACATCATTCGTAGCCTAAGGCTCCTAC
TCTCTCATGTTCGCTGAGATTCGATCGTCCCACAAAGGCTAGCATCCTCAGTTC
GTCAAGCTGTACGTGACCAGGTATGAAGGCCCTCGAATGAACTTGTTTCTTGAA
GACACGAGTTTGTGACGAAGATAGAGC
>index296_0
TTTCTGTTGGTGCTGATATTGCTACCACGGAAAGGACTACAAAGACAGACGTTT
CAGTCACCTGAAGTCTTGAGAGCGCAGACAATTCCGTATCAATAAGGCACGCAA
GGTTCAGGAGGTGAGAGCTTTACGTACAACCTTAAAGGCAACAAGGATGACCGG
ATGGTTTATTGACTGTCTAGAGCTAAAGGCTCCGGCGCCTCCTCTATTACCCTC
GAGGGAACTCCCTGTCGAAGATAGAGC
>index297_0
TTTCTGTTGGTGCTGATATTGCTACCACGGAAAGGACTACATTATGTTTACCAC
GGATCAGCTCATCATCGAAGTCCCGAAGGATCTTTGGGGCGTTAAGGCATTCTT
ACTTAGATGAGGCCTTAGATTCTCAAGCGTGTTTAAGGCCACTAGAGTGGAGAG
TACTAAGCGGCTTATCAATGTCTGTAAGGCACTCAGGAGTCTGTGCGTGGAACT
TCGCTCACGGGCAGATGAAGATAGAGC
>index298_0
TTTCTGTTGGTGCTGATATTGCTACCACGGAAAGGACTACAGTTAAGGAAATCA
AACTCAACGTCTGCGACAAGTTCAATGTCACGTGAGGATTAGGAAGGCTACCCT
ATCACACCGCGGGTTAAACTACAAAATGATGTCAAAGGCTGACGATCGCATAAC
CATGGCGTTCATCAAAGCTCCTGGGAAGGCCCCATGCCCATCGACGTCCTCAGC
GGTGAACCGCCCAAAAGAAGATAGAGC
>index299_0
TTTCTGTTGGTGCTGATATTGCTACCGACGAACAAACAACAATTCCGTTGTTCC
CACTCCCAACTTCCTTAGGCATTGCCATTAGTGGTCCCGAGCGAAGGCAGGTAT
GAGGTAGTCGGTGGGATCGTACCTATCCTCGGAAAAGGCACGTCGCACACCATC
TTCCAGCTTATCACAGCGTGACCGGAAGGCGTAGTGAACGAAGCTCGAGTTTTC
TGGCACTTAGCATGACGAAGATAGAGC
```

```
>index300_0
TTTCTGTTGGTGCTGATATTGCTACCTCACCAACTTTGTTTCACCAACATTAGA
CGTTCAGCTAGGTACATCTCACGCTTCGATCGTACCGGCTTATAAGGCTAAAGG
AGAAACTGCCCCATCGCCTAATGCAAACACATCGAAGGCCCGAGTACGGTTCTC
GTCCAGACTTATCGGCGTGAAAGTCAAGGCTATTACAACTCACTGAGGCCACGG
CTGCTACAGACCACTCGAAGATAGAGC
>index301_0
TTTCTGTTGGTGCTGATATTGCTACCTCACCAACTTTGTTTCCTACTACCCACC
TCCTCCCAACATGATCTGAGGTTACAGACGCCGATGATGCAAAAAGGCTACCTA
AGCGCATTCCGCCTCATCTCAACCCGTATTTGTAAAGGCAAGGACCGCTGTCCA
GAGAAATTGGATATCGCTGACAGTCAAGGCAATATTGATTTGAACCGGTAATAC
GATCTGGACACGCGGGGAAGATAGAGC
>index302_0
TTTCTGTTGGTGCTGATATTGCTACCTCACCAACTTTGTTACCGAGTTAACCTT
TCATCAACCCATCGCACATTCGGCTTATGCGCTGAACGCACAAAAGGCGTTCTC
TACAACGCACATCTCCGTACCTAACTTCCGTCTTAAGGCCTTCGGATCTTCTGC
AAGTCGCACCGAGAGAAGGTCGGTAAAGGCCCCTAAGCGAACACTGCAGTTTTG
GGAAGAACACTACAGAGAAGATAGAGC
>index303_0
TTTCTGTTGGTGCTGATATTGCTATCGTTCCAAAGAAGTCACGGATGACATTGC
ACTTCTGCAACACCTATCGAAGCGATTGATCTCGTTAGTGCTAAAGGCAGGGCG
GTAGTCTCAGACCCGTAAGGTCTCATCATTGGCTAAGGCCTTGTGAGTGCAGCT
TGGGTGCTCCAACTAAGCTAGGGCGAAGGCCGGCCATTGGTGTTCTTGATCAGG
TGAAGTAGCAAGCGCCGAAGATAGAGC
>index304_0
TTTCTGTTGGTGCTGATATTGCTGGCGGAGTAATGTCGTTAATCTACTAATGTC
ACCTCAGGTGTTCCCAACAGAGGGTTTTGCCAATGATCCCTATAAGGCGGACGC
ACTAATTGCTCGGCACAACAAAATACTACGTTAAAAGGCTGCCTGTCACAAATG
TCGGGAGATATACTGCGCGCTTCACAAGGCGAGATCATTACCGAAATCAGAGTT
GGATCTTTAGCCTGATGAAGATAGAGC
>index305_0
TTTCTGTTGGTGCTGATATTGCTTCCTACTACCCACCTCCTCAATAACATTGCG
GATTCGTACCTTGGCATAGTACAACAGCGTGGCTACTGCGAATAAGGCTCGTTT
GCAACGAAACCCTGCAGCTGGCCACGTGGGATTGAAGGCTAGTTACGTCGGACT
GAATGGGGCCTCTAGCCCTTTAAAGAAGGCTAGATAAGTACTCCTTCTTACCGG
GGTCGCTTTTGATCGAGAAGATAGAGC
```

>index306_0
TTTCTGTTGGTGCTGATATTGCTTCCTACTACCCACCTCCAATGTCGTTAAATC
CTATCAGCTTCTGCCACTTCGTATTTCACTGGCGCGTATCTCTAAGGCTAAGGC
ACTTATGACCCTTTTGCTGACCATTGATTGGTCGAAGGCGTATAAACACCTAGA
ACACAACACCAGTAGGCCGAGACGTAAGGCACATTAGCCACCCATTTAATCCCG
TCAGTGTATTCTTCTCGAAGATAGAGC
>index307_0
TTTCTGTTGGTGCTGATATTGCTTCCTACTACCCACCTCCATTTCACGAATCTT
TACTCAGCATGCACCACAAATGCATAGAGGAGCCTGCCTTATGAAGGCGTAGAG
AGTAGAGTCTGATAACGGGCGGCAGAGCAGATCGAAGGCTAGAACTGGTCTGCA
AGCGTCGGGACAGATTCTTTTCTAAAAGGCAACCATACTATGCGCCTAAGGAGA
GGAAGACCCATACCCTGAAGATAGAGC
>index308_0
TTTCTGTTGGTGCTGATATTGCTTCCTACTACCCACCTCCAAAGAAGTCACTCA
AAGTCATGGGACTACTGTCCGAAAGTGAACATGTGTGCTCGGAAAGGCTTACCC
CGATGGGAACTTTAAGCGGCAAACGTTTTGCAGGAAGGCTGATGGTTCCAACAT
TTCCACCGTTGAAGAGCACGGTTGAAAGGCCTCTTCACATATATCTTGGTAAGT
CCGACTTCTCAGCGTAGAAGATAGAGC
>index309_0
TTTCTGTTGGTGCTGATATTGCTAAACCACGTCCGGCCTGACCAGCTTTAATCT
ACTTCTGCAACAGAAGTCCGAAACGGCGTACTTAAAAGTGCAAAAGGCCCTTGG
GGTTGATCAGGGTGACAGGACTTTATACGTACACAAGGCTGTTTTCGTAGCGGG
GATCGATCATAAAACTCTCCATTCTAAGGCCTACATAGACGTTCGTACCCCGGA
TGCTTTACAATACTGAGAAGATAGAGC
>index310_0
TTTCTGTTGGTGCTGATATTGCTAAAGAAACAATCGTCATAAGTCCCACCAGGC
GTTTCACACTCGTAACCTGTGTGTGTGTTTGGCTGGACTACGAAAGGCTTATTG
GGGCTTGCCGGCCCATCATACGTAAGGACAAAGTAAGGCAATCTGAGACTTCTT
AACCTCAATGTACCTGTCCGTGGTCAAGGCACGACTAGATCGGGAGAACCCGAT
CCGGCTAGACCCATGTGAAGATAGAGC
>index311_0
TTTCTGTTGGTGCTGATATTGCTAACTCACCCACCAGATCAATGTCGTTAACTA
CTCTCATGGAGGATGCACGAACAGTAAGGGACCCGTCATTTCCAAGGCCACGGA
GTAGCGACATGGAACGAAAATCTTAGGAAAGTGAAAGGCAATATCTTGCAATCT
CTCTCACTGCGAGTATAGGAGTTTTAAGGCGTAGTCTCACTTCTCTCAGCTGCC
CAGCTCTATCAGTGACGAAGATAGAGC
>index312_0
TTTCTGTTGGTGCTGATATTGCTAATTCCGTTCGGACTTTACTCATCAACTGTT
ACCTCTGCATCGTGTTGTTCCCTGCTACTTACTATGGCAAGGTAAGGCGGCCAG
CGACAAGCCGTACACCGAGTTAATCGAGGCTCTAAAGGCCCGACTGCGTAACTC
AAAAGTACTTGTAGTTTGCGTCGGTAAGGCCAAGTTATCAATGATTGTGATTGT
TTTGTGCCCGCCGACGGAAGATAGAGC
>index313_0
TTTCTGTTGGTGCTGATATTGCTACCATCTTTCCCTAAAGAAATCCGTTACGCA
CGTTCACCTGATGGCATAAGGGTCGAGGTGTAGGAAACACTTGAAGGCATTATG
GGATTTCTCGTAGAGTCCGGGATCGCCGTTTCCCAAGGCAGTGAGCGTCCTGCG
CGACATTCTGACGCGATCGCGAACAAAGGCACTGGCGAACATTTCCTAGTCTCT
GTAGATATTAATGTGCGAAGATAGAGC
>index314_0
TTTCTGTTGGTGCTGATATTGCTACCGTTCTTCAGGCGTTACCAGCTTTAATCT
ACTTCAGCTACTCCATGCAGAAGGACTGCTGTTTACCTAATCAAAGGCAGCCTG
AGGTAGGTTAAAGTTCGCTTGTGACATAGCTACAAAGGCGGAGGGACACATAGC
GCAGTGAGTGTAACGCAGCCTTACAAAGGCGCACGTCTGGATACCGCCTCTGAG
AGGCACTTAGCGCATAGAAGATAGAGC
>index315_0
TTTCTGTTGGTGCTGATATTGCTAGGCGAAAGCGGACTTTAAAGAAACAATCGT
CATTCAGGACAGTTCCGAGTAGAAACTTGTGGGGATTATCCTAAAGGCGAAGTC
AGGAGACTCTCTTGGGCTCGTGCATGCGTGTTGCAAGGCATTCGAGTCAGGAGT
GCGCACAAGTATTTGGTTCCTTCGGAAGGCCGGTTTTAAGCGGCAGGTGTATGC
TGATGGGCAATGTTCCGAAGATAGAGC
>index316_0
TTTCTGTTGGTGCTGATATTGCTATTTCAAAGGTTTGGTCAAAGTTTCCACCCA
CTCTCCCTTACACCATGTTCCGTTCCATGCAGTACATGATACTAAGGCGCTAGG
ACTACGTTTTCGCACCTAAAACCCCTGGCACGACAAGGCGAGGCGCTATTAGGC
AACGTCAGCCTTCGCTATGGCAACAAAGGCTCACGAATTTATTTATTTCGAGTT
ATCATTAGTTAGATGAGAAGATAGAGC
>index317_0
TTTCTGTTGGTGCTGATATTGCTCCCACGTTCATTGGACTACCTCACCAACTTT
GTTTCCACATGTCGTCTTGATCACCTGCCTTCTTCAAGCTCTAAAGGCCATCAT
TCTCTTGAGCCTGTCTCGCCTAGTCAATATTAGTAAGGCACTGGACCTTTCACG
TCGAGCTGGGGTCCGAGTAATGTATAAGGCCCTGAATGTTTGTGCCACTGTGGC
TTACGGAGGTGTCTAAGAAGATAGAGC

>index318_0
TTTCTGTTGGTGCTGATATTGCTAAACAAACAAAGAAAGGAAACCACCACGGGT
TTCTCATGGACCTCAGCAGTAGACTTATCGGTTGCAATCAACCAAGGCCCAAGT
CATTTAGTCGCGTTTACCTTGAACTCATGTCGTGAAGGCGGTCGTCGCGGGCGC
TTAGGGATGCGCAATGGTGGCTAGGAAGGCCGTAGATAAGAGCCAACATTATTC
TACTGGGAATATACTAGAAGATAGAGC
>index319_0
TTTCTGTTGGTGCTGATATTGCTAACCCTAAAGGGAAGATAATGGAGGAAAGAA
GTCTCAGCTCAACATCCTGGTTGGTATCAACTCCTCCCTAGTGAAGGCAATATC
AGGATGCAGCTTTTGACAACTGGATTCCAAGGCCAAGGCCGCACCGTATAACCA
ACACGTTTGACCGGGGAACTGGCTTAAGGCTAAGTGCGAAGGAACTCTATGGTA
ACCCACGGAGCGTGCGGAAGATAGAGC
>index320_0
TTTCTGTTGGTGCTGATATTGCTAACCCTAAAGGGAAGATGTTTGCTTCCCACG
TTCTCGTAGGAACGCTGTGTGTACCTCATGCAGCGAACAACCAAAGGCAATGGG
GACTGAGGGACCTGACGTTTGCTCAGTTGCTAGCAAGGCCCGTTAAAAGTCCAG
CAACATCATAACCTGTGGACGGTCTAAGGCACCTCATTGCTAGAGGCTCAGATG
GCACAAGTACTCAGACGAAGATAGAGC
>index321_0
TTTCTGTTGGTGCTGATATTGCTAAGACGCCTCCTACAACATTTCGTAAACATC
GACTCAGTGAGTGATCAGACCTGGTCAGCCATGCTGGTCTTAAAAGGCATTTAC
TGGTCTACAGAGAAGACGGTCGACTTATTGAACTAAGGCTCTCCTACGTGGATG
TAGTATGACAGCTCTGGCTCCAACAAAGGCATGCGTCGTAATGATCTTTTAGGG
GTAGGGCAGAACAAACGAAGATAGAGC
>index322_0
TTTCTGTTGGTGCTGATATTGCTAAGTCGGAAAGGGCGTCAAATCCTAAACCTG
GATTCACCTGATGAAGCATCAGCTCCTAATGAGTTGAGGTACCAAGGCTAGGTC
TGGGGACTCGAATCCTTTGGCAACGTGAACTCCCAAGGCGCGTTGCTGAAGCGC
TTGGTTGCGTACTTTGCAGACTTCGAAGGCGGACTAGTGTACTTAATAGGGCAG
AGAGGAAAGTGGCTGCGAAGATAGAGC
>index323_0
TTTCTGTTGGTGCTGATATTGCTAAGTCGGAAAGGGCGTCAAAGAAGTCACTCA
AAGTCATGCCAGAGGTGTACTTTCGCATGCATGGTATGCCCAAAAGGCATAATG
GACAGTCGATCAGATCTTAGTGAGTTGCTGCCCTAAGGCAGTCTCAATTCCTAA
CTTCTGCCCCTCACGTCACCGCACCAAGGCTCCTGACATTCTGTAGTAGCTGAT
TCGCGGTTAAACGGGGGAAGATAGAGC
>index324_0
TTTCTGTTGGTGCTGATATTGCTAATGGAGGAAAGAAGTCACTGTTACCAACTA
CTCTCCACATCCTAGGGTTTAGTACTGGCACGAATACTTCGCGAAGGCTCCTCG
CGATGAATCACGGCCATCTAGATACTGAGTCCATAAGGCTCTCCCACACCGGCT
TTAACTCGCTTGATTGGTATAGGGGAAGGCTCGGGCAGCAGATATTACCTCGTT
CGTGGCCTCCCACGCGGAAGATAGAGC
>index325_0
TTTCTGTTGGTGCTGATATTGCTAATGGAGGAAAGAAGTCAGTCGCTCAGGGAA
GATTCAGTCCATCTGAGGTGGAATGGTAGATACATGAGTGAGAAAGGCGGGTGC
AAGGAGGCAAATATCCAAAACTACCTGAACTAGCAAGGCTCCCGTCGCGGCGTT
GCTCATGATGTTTATTAAAGTTAGTAAGGCATCTTCCGTAAGACGATGCCCCAG
TCTTGTACGCAACTACGAAGATAGAGC
>index326_0
TTTCTGTTGGTGCTGATATTGCTAATGGAGGAAAGAAGTCAGGGAGGAACGTTT
CAGTCCCTAAGAGCCTATTCGGACGTCGACATCGGATGGTCTCAAGGCTCAAAT
CGGCGACCTCTTTCCGCTTTTGCCGTTGGAGCTTAAGGCATAATGTTCTCATGG
ATGTCCACGTATACAATGGACAGAGAAGGCCCGAGGGTCGATTTGTGTTTCAGG
GGAAATCAGTTCCCCGGAAGATAGAGC
>index327_0
TTTCTGTTGGTGCTGATATTGCTACCTCACCATCGGTTGTAAACGTATTAGGTT
CCATCAGCTGAAGTAGGCTGTTGTCATTCCTTCAGTTCGATGGAAGGCGTACCC
GTATATGCGCTTGCAGATGATGTCCGCCTCCAATAAGGCCGACAGATAAGCATA
CAGAACATTGCAGTAACCCAGCCCCAAGGCTCACTCTGAACGCCCCAATGATGT
GCCCCGACAAGATTTTGAAGATAGAGC
>index328_0
TTTCTGTTGGTGCTGATATTGCTACTCAGTCAAAGAAGTCAAACTTCCCAACAG
GACTCACCAGTCAACTCGACAAGAGTTTCTGGTCACTGAATGTAAGGCCTTGGA
CTGCTCGAAACATGCATTGATGACCTACACGGTGAAGGCATATGATCACCCTCG
TGTCAGGTTCTAAGGTAATTTCCCCAAGGCACACGGTAAACTGTAGGACCCTTT
ACCTCATCCTAAGCTTGAAGATAGAGC
>index329_0
TTTCTGTTGGTGCTGATATTGCTAGGGAGGAAACAGATCAAGGCGAAGAAACCA
GACTCATGCTGCTTTGTCTGTCGACCGGTGTACCTCTACAGGAAAGGCGATCCA
GTCACTCTGCACCAGGTCCTCACTGGTGAACCAAAAGGCATGAAAATGCCCCGG
CCAGGAGCTAGCAATGGTCTCATTGAAGGCCTCCTAGATCAAAGAATACGCCTT
TGGCTCAGACACTTCGGAAGATAGAGC

>index330_0
TTTCTGTTGGTGCTGATATTGCTTCCGCCTCAAACCACGTAATCCATACCCTCG
TTTTCACGATCAGCCTTATGCGGCAAATGCAGATGCAGGACACAAGGCGAGCGT
GAAGTTTCCGACCGCTGAGAGAACAACATTTGTGAAGGCAAACCATCTGTCCGG
GCGCTTGAGTCGTACCCCGTTCTGCAAGGCCAGTATTCTAATTGTGTAAACAAT
TCAACGGCTTGCCCTTGAAGATAGAGC
>index331_0
TTTCTGTTGGTGCTGATATTGCTAAACCAGACAGTTAAGGAGGCGAAGAATCTT
TACTCCAACCTGAATGACGCGTTCACCTTGCGACTCGAATCCCAAGGCAGTCAG
GAAAATCTACCCCTACGCGCCTCTCCATCAGAAAAAGGCGCCCTCAAACTAAAG
AGATGGCGACACCATATTGTACGTTAAGGCGTAGATTACCCTGCGGCAACAATA
CTGTGCGTAAGTTCTTGAAGATAGAGC
>index332_0
TTTCTGTTGGTGCTGATATTGCTAAAGAAAGGAAAGTTATTCCGCCTCAAACCA
CGTTCACACGTCAGTCCTATGGGCTGAGTCACTTAAGTATCCGAAGGCAATCCT
GGACGATACCCAATCACCAACAGATATCATAGACAAGGCTGGAGGAAGGGCGGT
CTTATTTCACTGTTTTGCACAAACCAAGGCTGCGTCACAGGATAATATAATGTC
AGACTCTGATCCTCTCGAAGATAGAGC
>index333_0
TTTCTGTTGGTGCTGATATTGCTAAAGAAAGGAAAGTTATTCGGACCACCCTCG
TTTTCACACTCCAGCACAGAGACAGCAGTCAAGGAAAGAACACAAGGCCGCACA
GAGAACTAATCTATCGACTCGTTGCCTTCAGGAAAAGGCTAGCACCCTTGGGAG
GTAGCCATTCGCACACTGACAAAGGAAGGCTCGCATTCGAGCTTCTTTGAGGTT
GGCGATGTAGAGCTAGGAAGATAGAGC
>index334_0
TTTCTGTTGGTGCTGATATTGCTAACTCACCCGGACATTTAAAGAAAGGAACAT
ACCTCCCTTACTCCCATAAGGAGCTTACGGGGATCTTGAGGACAAGGCTTCCCT
TTTACGTATAATCCTATGTCGATGCCTGATGTACAAGGCGGTTCTGTCTCTTAC
TGTTTGCCAGTAGCTTGGGCTGCTAAAGGCTAGCTTTACCATAGTTAGGAATTC
ACACGTGGGCGTTGACGAAGATAGAGC
>index335_0
TTTCTGTTGGTGCTGATATTGCTAAGTCCCACCAGGCGTTATTATGTTTGTTTG
CTTTCCCATCTCTGACGAGAGAATGATGCGTTCGACTACGATGAAGGCATTTTG
CCGGTCTAGCGAAGAGGAATGCGTCTTCCTATGCAAGGCCACTCAACCACTGTG
ACATTTTGCCGTACGCTGTCCAACCAAGGCCGATTTAGACCAACCAAAACTCTC
TGGTTTACGTCCACGAGAAGATAGAGC
>index336_0
TTTCTGTTGGTGCTGATATTGCTAAGTCCCACCAGGCGTTCCCTAAAGAAAGTT
CAATCATGCTGCATAGTCAGGTATCAGGCCCGACTCTAAATGTAAGGCACTGTG
CAAAGTATCTGAGACATCGCCGGATTACGAGTGTAAGGCCGTCACACGAGCTAA
CGCCAGCTGTCATTCGCTGTCTAATAAGGCCCGATTCGTCTGTCGTTAAGGTAT
CGGGGCAAAGTTCCGCGAAGATAGAGC
>index337_0
TTTCTGTTGGTGCTGATATTGCTAATCATGTCAACATGTTACCACGGAAAGGAC
TACTCCCTTACACCATGTTGACGGGACATCCAAAACAACCGCTAAGGCCGGGGA
GTCTAGCCTTACAGGTAGTGACCAGACGTTGCAAAAGGCCGAATGTGGACGATG
CGTACCTTTGTATTATTGCGCAAAAAAGGCTCACTACGCGGGAATTCCGCCAGT
AGAATATAACTTGGATGAAGATAGAGC
>index338_0
TTTCTGTTGGTGCTGATATTGCTACGGATGACAGTTGGTTAAACTCAACGTTCC
CACTCCCTTACAGGTCTCAGTCTCCGAAATCAAAAGCCTACTCAAGGCCTCCAA
TCCTGCCTCGGAAACTAAGCTGCATCTGTCCTGCAAGGCCGAAGTCATGGTGGC
CACTGTCGGGGCTAAGGTGGGACATAAGGCAACATTATACTTATCGTAAATGCC
TCATTACGTCTCTGGTGAAGATAGAGC
>index339_0
TTTCTGTTGGTGCTGATATTGCTATTATGTTTGTTTGCTTTCCTACAACTCCGC
CTCTCAGCAGTCAACGTTTCACTTCCCCTACACACTACGCTTCAAGGCTCCAAT
ATTCGGCATTTACTGAAGCGCTTCTCATACTGACAAGGCATGTAGAGTGGAGGC
AGTTACACGTGGTCATGCCCAGGTAAAGGCGGGCTAACGACTCAAAGGTGCAGT
GGCCTGTTCTCGAAAAGAAGATAGAGC
>index340_0
TTTCTGTTGGTGCTGATATTGCTATTATGTTTGTTTGCTTTCGGACCACCCTCG
TTTTCAGTGAGTGGAGTAACTCACAACAGCGGTCTCAAAGGTTAAGGCCCAACG
CAGTGTAGTTCTATGAGGGCACCGAACGCCGCAAAAGGCCGTGCCACTATCGAC
CCGAGAGAGATAACCTAAATATCGAAAGGCATGAACGCGCCCTGTCTTGGTGGA
TAGAATGTTGCTCCTCGAAGATAGAGC
>index341_0
TTTCTGTTGGTGCTGATATTGCTATTATGTTTGTTTGCTTTGTCGACGGATCCA
CTCTCACCTTCCTTGCGAGTATTCGTACGTCGATGGGTACGATAAGGCAGTCTT
CCGCCACTCTTGAGCGAGATCCATGGTATCTTGGAAGGCGTCCTACCACACTGT
AGCCTGTAACGCTTAGCCTCCGTCTAAGGCATTTGAACTCCGTTACTCCCGGCG
CGGCCAACATTTACGGGAAGATAGAGC

```
>index342_0
TTTCTGTTGGTGCTGATATTGCTGGGTAGGTTTCCCACTTAAATAAGGAGGGAA
GATTCACCTGAGTTTGTCCCTGTACGCTGGTAGGTGGTGAAAAAAGGCCGAGTG
GGAACATCTAGGTCTCTGTCTCAAGCCAATCTAGAAGGCACTCGGAAAACGGCA
CATTTGCCCCATCATGCTTGGGCGGAAGGCACCTGGGGAATCTATCTCATGTTA
TAAGGTCACCTCCGCTGAAGATAGAGC
>index343_0
TTTCTGTTGGTGCTGATATTGCTGGGTAGGTTTCCCACTTACCACTTGCAAAGA
TCATCAGCTGTCAATCTGAGCATTGACGGGATCTGTGACGACAAAGGCACTCAG
ACGGCGCCACAGACGATTCGACGATATGCGGAACAAGGCTAGGTTTCGTCACTC
TTACCATCCCGTCGGAGCACACCTCAAGGCTTGTGCAGCTAGATTTCGAATTCT
TCCGGCTATCGCCACGGAAGATAGAGC
>index344_0
TTTCTGTTGGTGCTGATATTGCTGGGTAGGTTTCCCACTTAAAGAAACATTAGA
CGTTCAGTCAGGATGGGAGAATCACAGCGCTCAGCTTTGTCCAAAGGCTCTCAG
ATTTCTATGGTACACCAGCCAGTCAGCTAGTGGTAAGGCCTCGCACTTATTGGT
AGAGGTAACTTGACGTATTAATCGGAAGGCACAATCAGTACGACCAATACGAGC
GGGACCAAACCGAGCTGAAGATAGAGC
>index345_0
TTTCTGTTGGTGCTGATATTGCTTCGATGGATAAGGACTCAAAGAAAGGAACAT
ACCTCACGTAGGTCCTTCAGGTTATCTGGCGCACGATCCAGTTAAGGCGCAGTG
GATTGGAATCCCTGTTGTAAAGTGGGGAGAGAAGAAGGCTCAGTCGCCATCCGT
TAAAGATGCGCCACTCGACCTTGGAAAGGCACGCTAGGGGCCGCTGAACAGACT
TTTCCGGGGCTATTTTGAAGATAGAGC
>index346_0
TTTCTGTTGGTGCTGATATTGCTAAAGATCACCCACGTTCAAAGATACATCCCA
CTTTCTGCTTCTGCACCAAGAACTTTGTCACCTCTCGCGTTTCAAGGCCGCGGA
TGCTGCAGACATACTTAAATACTCCTGGTTTAAGAAGGCCCTGAAGGAGATGTA
TAGAAACTCCGCACCAACTTCAGAAAAGGCGCCCCGTAAAATTAGCCTTAATTG
TCAAGCGTGAGTCAACGAAGATAGAGC
>index347_0
TTTCTGTTGGTGCTGATATTGCTAAATAAGGAGGGAAGATAGGTGCGTCATCGT
CATTCAGGAAGCTATCTCTCTGGCGCTGTAGTTATCCTATAGAAAGGCCCGGAC
TAGTTACCCACATACCAGCGGCGGTTAAAAGGATAAGGCATCATAAGAGTGTTG
TCCCTTTTGGGAGGTCTTCTATAACAAGGCCGGCCCGAAAGGTTAAGGCAGCAA
GTAGCGCCCCTGACTAGAAGATAGAGC
```

```
>index348_0
TTTCTGTTGGTGCTGATATTGCTAATCCATACCCTCGTTTCCCTAAAGCCCTCG
TTTTCACGATCTCACACCCTGAACAGAGACTCTTGAGCCCGATAAGGCTACTCG
ATAACCACTGCATTGTAGACGACCTGGTCTGAGAAAGGCGGCGACGCACCCTGA
CAACTAAACCTTCGGATAGCGGGGAAAGGCGTGGTATAGATTCTCGCCACTCCT
GCCGGTTGATTGGACCGAAGATAGAGC
>index349_0
TTTCTGTTGGTGCTGATATTGCTACCGCATAACCAGCTTTAGTTGGTTAATGTC
ACCTCACCACATCCTGATGGAAAAGTGGTAGCGCACATTGAACAAGGCATGTTC
TCACACCTCCATACTCATACCTGGATCAGATGAGAAGGCGTCGAGACGCTAAAT
AACGCCCGAACCGCCACTACTAAGGAAGGCCTCCGTTGTCTCCAGGATCCGAAC
TAGGCCGTCTGAGTACGAAGATAGAGC
>index350_0
TTTCTGTTGGTGCTGATATTGCTAGGGAGGAAAGTCCCACCAGGCGTTGGGTCT
GAATCAACCACACAAGCAACGGCTACAATCTGCAACTTCAACAAAGGCTCACGC
ACAGTGAGAAGCTGTTGCAACCTTCAAGAGAGGCAAGGCAGAAGCCGGTCCTTG
CCCCAGCACCGTCCTGGCAGGATCTAAGGCAGATGTCCGATAAGGGAAAGACCG
TCGCGAACGTTCTCGTGAAGATAGAGC
>index351_0
TTTCTGTTGGTGCTGATATTGCTAGGGAGGAAAGTCCCACTGGGTATCAAAGAA
GTCTCTGCTTCTCTTGACTGGTGCCATTTCCTGTAATGGACGAAAGGCTAGGCC
CAGATCTCTCCGACTTTTGACGCCCAGCTCACTCAAGGCCGGTTAGCTCGGGTC
TACGGACACGACAACCTGTGTCAATAAGGCAAGCCACTTAGTTCGGTAATCCCT
CTTCCGCGGCTATAGGGAAGATAGAGC
>index352_0
TTTCTGTTGGTGCTGATATTGCTAGGGAGGAAAGTCCCACTCCGGAGGAAACTC
AACTCACACGAAGGACATGCAAGCCCTCAACGTATCCGTAAGGAAGGCAGGACA
CTAAGCTTCAACGACATCCTCCTTGAGTAACTCCAAGGCTGTTTACTTTGCGAT
GCATATGGGGCGTCGGAAGGGGCCCAAGGCTTGTTTAAGACGTGACACGAGTAG
GTCCCCGTGTGGGCGAGAAGATAGAGC
>index353_0
TTTCTGTTGGTGCTGATATTGCTGTTCCCACTCGATGGATACCATCTTTCCCTA
AAGTCAGCTCTACGGATAGAGGTTGAATCAGGAAAGCGCTCCTAAGGCGTTCGG
GCCTTGGACCATCAAAATACGCGCGTAGGCGTGAAAGGCACGTTCATATGCTAG
GGGCCAAAACCCGACGACAAGGTGCAAGGCGAGTAACGGAGCAGACCAAAGCCG
GTGAGAAATGTCAGCTGAAGATAGAGC
```

>index354_0
TTTCTGTTGGTGCTGATATTGCTTCCGCCTCAAACGTTTAAATGTCGTTAACTA
CTCTCAGGACAACTTGCGGACATGGTTTAACGAATCCACGCCTAAGGCACGTCG
ATACACTGGCCCCTCCCATAATGGCTCGTATCCAAAGGCTGGAGCTTATATAAT
AATGTTCATGCGCCGGAGGGCCTAAAAGGCGCTATTAGGTTGGCGTTCCCGGCG
CCTCACAACGACGGGTGAAGATAGAGC
>index355_0
TTTCTGTTGGTGCTGATATTGCTTCGGACCACCCTCGTTTAAAGTTGCAAGGAA
AGATCAGCTAGGACCAACAGCAGGAAAGATTGAGCTAACAGCAAAGGCGCATTA
CAGGATGCTCTCCGCAACTGCAGCTGACGCAACAAAGGCTACCATTCTGAAAAC
AGAAAACCGCCTTTCATTCCCGTGCAAGGCTCAAGGGCAGGCGTCTAGCGAGTT
TATCATGCGACTGATCGAAGATAGAGC
>index356_0
TTTCTGTTGGTGCTGATATTGCTTCGGACCACCCTCGTTTAACATGTTATTTCA
AAGTCATGCTGCAAACCACCAGCATGTTTTGTGATGAGTCTGTAAGGCACTCCC
TAACTCCCTACATCACCGTGCACTTGCCACACATAAGGCCTAGTACATCTAATC
GGTACTAATGAACCGACCTCCGGGCAAGGCGCCCCATAGGGTTGTGGTTCGCCG
GACTACATGCAATACGGAAGATAGAGC
>index357_0
TTTCTGTTGGTGCTGATATTGCTAAAGAAAGGAACATACCAGTCGCTCACTCCA
TCGTCATGGTCCACTTCCCTAATGGATAGTGTGCCGCGAGGTGAAGGCAAATCC
TTCATGAGAAATTCGAAACCTAACGAATGCCTCAAAGGCGAATGGGAACCCCGG
CCAGACGTGCCCCGCCGTCTTCACAAAGGCGCTCTCTGTCAGTAGCTCGAAGTG
CGAGGAACGAGACCGTGAAGATAGAGC
>index358_0
TTTCTGTTGGTGCTGATATTGCTAAAGAAAGGAACATACCACTCAATAAGGTGC
GTCTCAGCTCTTCTAGGTCCAGTAGAGGAGAGGATCGTCTGCCAAGGCGATGCT
GAAGGCAGTGAGTATCCCCTACTTTGTAGACAGTAAGGCCAACGGTGCAGTTAA
TCGGCCAGGCGGCCTTATGCCACCTAAGGCGCCACTGTGTGACTAGTGCAGGAT
TCTCTGTAGACCGGAAGAAGATAGAGC
>index359_0
TTTCTGTTGGTGCTGATATTGCTAAGTCCCACGTTTGGTCCATCTGTCAACCCT
AAATCACCTGATGTTCTGCAACCGTTCTCATTCGCGCGTGAAAAAGGCCCCCTT
GGGTCTTACTGCATATCGTTTAAAATTGTAGAGTAAGGCACGTGCGTTATGGAA
CGGCAGGAGGGCCTGAGCTGAAGGCAAGGCACTCAATGTGGAGGACTCTCGATC
TGAAATTAGACGCGTGGAAGATAGAGC
>index360_0
TTTCTGTTGGTGCTGATATTGCTAAGTCCCACGTTTGGTCAAAGGATCACGGAC
TTTTCGTAGAGCAGTTAGTGACAGGAATCTGAGGAGAGGTAATAAGGCAGCCAG
TAGGATTACACCGCACCCAGCTCCCCGTTCGTGTAAGGCTGACCGTCTCATCAA
GCCAACGGGGCTCGAATTAACTGGAAAGGCGGACGAGTGCAATGAGACGACCAG
GACATAGCTGCTACTTGAAGATAGAGC
>index361_0
TTTCTGTTGGTGCTGATATTGCTACCCTCCCAAAGTGGACTCCTACTAATTATG
TTTTCACCTGAGTCCAGTCTGTTGAGTTCCATATACACCCCACAAGGCAACTTG
AGTAACAACTCCGGATAAGCAAGACCTGCTAGAGAAGGCAACCTCCCCATCGGC
CAGCGTATATAGAACCTGTCGCCCAAAGGCCTAGTTTGACGTACCAAGCAAGAT
AGCTAGTGCGCCCCAAGAAGATAGAGC
>index362_0
TTTCTGTTGGTGCTGATATTGCTACGGATGACATTGCACTACCGACGAAACTTT
GTTTCCACTTCGTTCCGAATAGGGTATGATGTGATGTGGACAAAAGGCTGACCC
TTGGCCCTCATTGGTGGAGTGTTAAGAACTAAGTAAGGCGGGCCTGGGACAACC
TCTTTATTGCCCGTTTTATTAAGTGAAGGCATTTAGCGGATGGTCTACACCCTA
TGTATTGAGGTTCCCTGAAGATAGAGC
>index363_0
TTTCTGTTGGTGCTGATATTGCTATCGGACTTACCTCACCTCAATAACCACCCA
CTCTCGCATGAAGAGCGTGTAGTGAGTATTGCATCGATCGACGAAGGCCTGAGA
ATGCTTGGCTGCTGCCATGCTAATTTTCCGTGTGAAGGCAAGGCATGGGAGCAT
TTGCCTGGGAATGGCTGGTTCGCTAAAGGCCAACTCGGGGACTTGGTAGGGTTC
GGATCGGGCAACCGCGGAAGATAGAGC
>index364_0
TTTCTGTTGGTGCTGATATTGCTATCGGACTTACCTCACCAGGTTTATCAATCA
ACATCCCTATCACATGGGCATCTGCCAGTGTCACATGTGGGTGAAGGCGTACCA
TTTGTTTGCAATGTGAGCTTCGTTAGACTCCACTAAGGCCTACATGGTACGTGA
GCCACAAAAGAGCACTACCACCATTAAGGCATCCACTTTTCGATGCCCCGAGGA
TGGCGCTGGGCAAGCAGAAGATAGAGC
>index365_0
TTTCTGTTGGTGCTGATATTGCTTCCCAAACCAGTTGGTTACCACTTGCAAAGA
TCATCAGCTCATGTACCGTGAAGCGTGGTCTAAGTCGGGTCTCAAGGCAGTTCT
TCTTAGAATGTGATCTTGACAATCCCGGACATAAAAGGCGAGCCCGACAGCGTG
TAATGCCGTATAGGTACCATAGAACAAGGCCTTCAATCATCTGGAATCGAGGCA
GGTGGCGGTTACCGGTGAAGATAGAGC

```
>index366_0
TTTCTGTTGGTGCTGATATTGCTTCGATGGATCACCCACGAAACTTCCCGCTTT
CCATCACCAGAGTCCACTTCGACCTCAGTTTTACGGGATGAGTAAGGCTTACTG
CACGCTGTGCGCTGCAATACCCCTCGATACGACCAAGGCTAACAACGATTCGCT
CTTGCGCCGACACGTGAGGATGTCGAAGGCGGAGAAAATAAGACTCCGTTAGAC
CACGGACCCGATGTGGGAAGATAGAGC
>index367_0
TTTCTGTTGGTGCTGATATTGCTAAGGACGTAAGGGCGTCAAATAAGGAGGGAA
GATTCCAACCTCAACAAGCGATCCCTTTGTTCAAACTCGCAATAAGGCCCCTCA
GTCACCATGATCAGGGCTTACGTTTGGCCATCTTAAGGCTGTAAACTTCATCTA
TGTCATGTCCCTGGCGATCGATAGGAAGGCGCCGGGGCTTAGGAGGTTCCCCTA
TGGTTAACACCATTAGGAAGATAGAGC
>index368_0
TTTCTGTTGGTGCTGATATTGCTACCGCTTTAAGTCGGACGGGTCATTTGTTTG
GTCTCTGCTCACAACTTCCATGTTGGTAGTCTGGAGCTCAGGGAAGGCTGGCGT
TCTTGGGTATACTCTAACCTCTATCCAAATTCACAAGGCACCTTCGGAAACAGA
GATGCCTAGACCAAGTCGTAACCAGAAGGCGCAGGCGAGCGACGTTCTGACGGT
GGTGCGTTAGCACGGGGAAGATAGAGC
>index369_0
TTTCTGTTGGTGCTGATATTGCTAGGGAGGAACGTTTCAGCAGGCGTTTCGATG
GATTCTGCTCAGTTCTTGGGCAGTCGAATTACGAATCACCAGCAAGGCGATTCT
ATCTCCCCGGCTCGAGGATAGGGGATAATTAGGTAAGGCAACTTTAACTATATC
TCGTAGAGCTGCATGCCAGGGTGGAAAGGCGACTTTGGGTGACGCTTTAATGCG
GTGACCACAAAATAGGGAAGATAGAGC
>index370_0
TTTCTGTTGGTGCTGATATTGCTAGGGAGGAACGTTTCAGACCGAGTTAACCTT
TCATCCCATCAACGTAACGGCAATTACGGAACCACAGTTGCTGAAGGCTGCGTC
AGAGGTAGGGTACCGACAGACCTAGACTGGAGGCAAGGCCTCTTGTGAGAGCCT
AGGTCCACAGATCCTCTTTGGCGTGAAGGCTCTGAAGGCTCGAGTGTCTCCTCA
TGTTCGGTGAATACCAGAAGATAGAGC
>index371_0
TTTCTGTTGGTGCTGATATTGCTATTTCACGAATCTTTACTCCGCCTCAATTTG
CTGTCACCTGTGTGCAGAACAACTCCAAGTCTCGTTCCGTTTTAAGGCGATTAC
GAAGTCAGTTCGGCAGGTAAGAATGACGCCACGAAAGGCTACTTCTGATCGAAG
TATCAGGTTTGACAAATCGCTGTATAAGGCCGGCCGTAACCTGTCTGATCGCAG
GATGCACCCGCGAATCGAAGATAGAGC
```

```
>index372_0
TTTCTGTTGGTGCTGATATTGCTCCCTAAAGAAAGTTCAAGGGTAGGTTTCCCA
CTTTCCCTATCGTTTCAGTGTGTAGCATGCGTATACTGTGCGTAAGGCGGTAGC
GCATAAGAGGGGAGAAAACCTATGGGACGTTAATAAGGCAATCGAATCTCGCTA
TGTCTAAAGCAAGTGGGTAGCTTAGAAGGCTAGTGGGCTAGCAGTCAAAGTGAT
AATCGTGATTGGGCTCGAAGATAGAGC
>index373_0
TTTCTGTTGGTGCTGATATTGCTGTTTGCTTAAGTGCCGCAACTTTGTTAAACA
GCATCTACCACGAGTTTCTCTGGGTACTATCGAGACGATACGAAAGGCATCAAG
TCACGACGGCGCAGCATTCAGCGTTATGGGGTCGAAGGCCGAGGGTAAAGGCCA
CACTAGCTTCTCGATAACATAACTCAAGGCAGCAGCCCAAGTGCTAAGCTTTGT
TTCTAGTAATGGATGGGAAGATAGAGC
>index374_0
TTTCTGTTGGTGCTGATATTGCTTCCGCCTCAATTTGCTGCCCTCGTTTAAACT
TTATCAACCCATGTAGACGAACATCTGCTCATGCAACCTATCCAAGGCACTACA
GCGAACGCGCAGAAACGAAAGGTTTGACAAGGCTAAGGCCAAGCGTGAAGAGGT
CAAGGTAATGGGATTCCCGTGTGGGAAGGCCATCATGCGAGGACGATGACAGGC
TTCTTATTTTCACCCTGAAGATAGAGC
>index375_0
TTTCTGTTGGTGCTGATATTGCTAAAGAAAGGACCGAAAGAATCTTGTCCTCAG
TACTCCAACGTGACTGGACTCTGCTAAGGGACTCGAGAAACGTAAGGCATACCG
GATGGTTTGATCTCTACTGGAGCTCCCCACCGATAAGGCAATTGCCAACCTGCC
TCCAGATCGAAGCCCTCGAGATACAAAGGCATTCTATGGTGCGTGTGACTGCGA
ACGGTCTATAATTGTAGAAGATAGAGC
>index376_0
TTTCTGTTGGTGCTGATATTGCTAAAGGAACTCCCTAAAGAATGGAGGAAACCA
CGTTCTACGTGACGCTTATCGCAGACCGACAAGACAGGTGGACAAGGCTGAGGA
AGTCGTGATTTTACTCCCCGAAATGACGATGGAAAAGGCTAAATGCTGGCTCTT
GGTAGGATATACCACTTATAATAGGAAGGCTGAGGCTCTATAGGACGCGGAATA
CGACCTCCTATAGGTGGAAGATAGAGC
>index377_0
TTTCTGTTGGTGCTGATATTGCTAAAGGAACTCCCTAAAGAACGTATCATTGCC
GTCTCACCTTGACATGTCTGAGATCGCGTTCGTTGTGCAATGGAAGGCTGCTCT
CAACAAGCCGGGATGCCCGTGTAGCCAGTCACTCAAGGCGAATGGGTGACAGCT
TCATTGAACGGGCGAGGATACTTATAAGGCCATAGCCGTATAAATGCAACTGAA
GGAGGCTCTACAGATGGAAGATAGAGC
```

>index378_0
TTTCTGTTGGTGCTGATATTGCTAAATCACCGACGTTCTTACGCACGTATCGGA
CTTTCATGCGTCAGGTTTCGAATTGTGGCTCTACACTAGCTGGAAGGCTTCTTC
CCCTACTTACTCCAACGCATAGCGTTAACTTGATAAGGCATCTCTAGCAGAGTC
ATAGGGACACTAATGGTTGCCGGGCAAGGCACGGCACCTTGTCTACGCATTCTG
TGGCATCTCCGATCCCGAAGATAGAGC
>index379_0
TTTCTGTTGGTGCTGATATTGCTAATCTGTTTAACGTATCACCGTTCTTCAGGC
GTTTCGTAGAGAGGTCAAGGTTAACCAAGGGTACGGCAGACCTAAGGCCGGATC
GTGTCGTTTCGTGGCATATCTGGTCCTGAGACACAAGGCGATATACTAACTCGC
TGGTCGCTAAGGCCTGCGCTACTACAAGGCCGGATAAGTAGTGTTCGACATTTC
GATTGGCTGTGGCTAGGAAGATAGAGC
>index380_0
TTTCTGTTGGTGCTGATATTGCTACCACTTGCGTTCTATCAACTCGTTAATTCC
GTTTCATGCCACAGGTCTAATCGCTTATGCTTCCACCTCAACCAAGGCTAGTAT
GGGGTTGCTTAAGGGATGGTAGGCACACACGTTGAAGGCGACCTGACGCGTGAC
CAGGAGCACTCGATTAAATTGGAGTAAGGCTTTCAGCTACACGTGGTTGCGCCA
CGCAAGATGACCCAGAGAAGATAGAGC
>index381_0
TTTCTGTTGGTGCTGATATTGCTACCACTTGCGTTCTATCAAGGACGTAAGGGC
GTCTCAGCTAGGATAGTGACTGCGTTAGGCATAAGTGACCAGTAAGGCCGGGAC
CCACACATCTGGAGGCTTCTAGGAACGCGCATCAAAGGCTGTGTTGTATATTGA
CGCTCCTTACGGTAGTGCGCAACTAAAGGCTATTGCCTCATAAAGCACCGGGAT
CTGTCCAGACGCTCTAGAAGATAGAGC
>index382_0
TTTCTGTTGGTGCTGATATTGCTACCCTCCCACCCTAAAGAAAGTTGCAAGGAA
AGATCAGCACATGTCTAGTGGACCAGTTGGGTAGGCACCATGGAAGGCGAGATG
GACTTAAAAGGACCATGGATTTTATCAGTATACGAAGGCAGCTAACAGTCGCAG
GCACGAAATACGAATGGATACCCGAAAGGCTGCAGAGGGGCATCGGTTGGCCTA
GACTGATTACCTGGTCGAAGATAGAGC
>index383_0
TTTCTGTTGGTGCTGATATTGCTACCCTCCCACCCTAAAGAAAGACAGACGTTT
CAGTCAGCTACTGGCTACCACTTAGTCGCCTGTATTGCAGTAGAAGGCATGCGG
CGTTTCTCGGGTGAGACATCCACTCTTTACCCCGAAGGCCGGAGATGTTCGCTG
CTCTGCTTAATTGGGAGCTACCTATAAGGCTACGTGGCTACACGGCGCTACGCC
GAACTCTTTATGACCCGAAGATAGAGC
>index384_0
TTTCTGTTGGTGCTGATATTGCTAGGGAAGATAATCCGTTGGCGGAGTAAAGAA
ACATCTGCATCCTTTGTGTGCACAGTGTGTAAGGGGAATATCAAAGGCCTGGTC
TATCGATCCACCCTGATCGAAGGACTAAGGGAGAAAGGCAAAGCCATATACAGC
CTCCTTCTCACCCGGGAGGCAACTTAAGGCGCTTCAGCATCCTAGCTTTTACAC
AGAGTATCCGAACAGTGAAGATAGAGC
>index385_0
TTTCTGTTGGTGCTGATATTGCTCCCTCGTTTAAACTTTAAGGCGAAAGCGGAC
TTTTCACCTTCCTTTGTGCCTTTGCCGCTCAACATATAGACCGAAGGCAGCTGG
CCACTATGCTGATGGAGGCTGAAGTTCAGCCTCAAAGGCCCGCGTATCAGAGGC
TCGTATATGACGCTAGAGCGTTACAAAGGCGAAATCAGCACCACCCTTGATGCA
CCCGAATACGCTTATTGAAGATAGAGC
>index386_0
TTTCTGTTGGTGCTGATATTGCTAAGGACGTAATGTCGTTAATGGAGGAAACCA
CGTTCAGCTTGGAGAGCAGTCTCAGCTGTTCGTATGTTCCTCAAAGGCCGAAAT
ATGTACTTGTGGATTCGAAGAGGGTGCACGTTATAAGGCGACGCGCAGTACTAA
CGCAGTCAAAACCTTTCGGCGGCTGAAGGCTAGAACGTTGCTATGGGATTCGGT
AAGCTGACTCAAGTTAGAAGATAGAGC
>index387_0
TTTCTGTTGGTGCTGATATTGCTAAGTCTTTAAAGAAACATCCTACTACCCACC
TCCTCAGGTGATGACGTTCTCGTAGGAGGTATGGGGCAGTTTAAAGGCGTCGAA
ACTCCCGTAGCCCGGACAGAGATACCAATTCATAAAGGCTTATAGTTTAGTGAT
GCGAAAACGCTCTAATGAGCTCACCAAGGCTTGACCCTCACGAAATCGCATCTG
AGTAGAGCTCCCATCAGAAGATAGAGC
>index388_0
TTTCTGTTGGTGCTGATATTGCTACCCTCAGAAAGAAAGGAAGGGCGTCAATAA
TTTTCGTTGACTCTTCGAAGGATCAAGACACACATGGTCGCCGAAGGCTTATTA
TCGCTCTTAACTGCAGAGGATGGTTCCAATATGAAAGGCGGAGCGACGGCACCA
CCGAACTGATTGCCCCTATTTCTTTAAGGCATGGGTAACCTTAGGTCCCATTTG
GACAGTTGAGCCTCTAGAAGATAGAGC
>index389_0
TTTCTGTTGGTGCTGATATTGCTACCCTCAGAAAGAAAGGAAGTCGGAAACATC
GACTCAGCTGATGGTTACCATCAACCTAGCGGAAGTCTTCGTTAAGGCACCGGC
CAGCCTTTGTGCGAATTAGGACTTCCGACGACTGAAGGCCAGTGTCCTTAGAAA
ACACAAGGCGTGCCAATCAACAGAAAAGGCCGGTCGTCCATCGGTAGTTTAATA
CGCCGGGTGTAGCTGCGAAGATAGAGC

```
>index390_0
TTTCTGTTGGTGCTGATATTGCTAGGGAGGAATTCTGATCAAAGAAAGGAGGAA
AGATCTACCAGCAGACCACTTGTGTACACAGCTAATAGACGACAAGGCTGGCCA
CCGCTCCTCACCTATGGGACAAGGGTCACGAGAAAAGGCAGTAAGTGACCCTTA
TTCCTTCGCCGTGCAAACCGTGTAAAAGGCGCGGGCGCGAACCACTGCACGACT
AGAGAAGTAGATATGCGAAGATAGAGC
>index391_0
TTTCTGTTGGTGCTGATATTGCTATGGGTTTAACTCACCCACCACCCATCCTAC
AACTCAACCACACGGTCAGTGACCTTCGTTAATTAGACCAACCAAGGCCTGGTT
GTTGAAAGGTGGTAATGAGTTCTGAGGTAGTGCTAAGGCTAACGACAGGCATAC
TAAGACTGTGGCGTCGGCTACCTTGAAGGCCGTCCGTTCTCCTCGCCGGTCGAG
AGTCTTCGAGGTACGCGAAGATAGAGC
>index392_0
TTTCTGTTGGTGCTGATATTGCTCCCTAAAGAATCATGTCATCGTTCCAAAGAA
GTCTCAGCAGTCTCCAGTTATCCAGGGTGTGTCCACCAGAGTGAAGGCGAATAG
GCCTCTCCTGTGTCGTACAAACTAGGATTCCAATAAGGCCCCTTAACCACCGAG
ATTACAGGATTTCATGATCCAGTACAAGGCAACATCGCAATTTGGGATTATATC
AGGTGATGAGACAGGGGAAGATAGAGC
>index393_0
TTTCTGTTGGTGCTGATATTGCTGTTTGCTTCCCACGTTCACCACCCATCCTAC
AACTCACCTGATGGCACTTGCTTCTTATGCTACCAGTCGCGTCAAGGCTGTAAG
GTAATCCCTCTCCAATCCATCCGTTTACTATCTGAAGGCGACCCAGCGACGTCT
AAGTGAGGTGTGGGTGTAGTAACAGAAGGCTATTATTCCCAGCTGTGCCGTTTT
GAGTCCAAGCTTAGGAGAAGATAGAGC
>index394_0
TTTCTGTTGGTGCTGATATTGCTGTTTGCTTCCCACGTTCTGGGTATCAAAGAA
GTCTCATGGAGTGAGTTGGTGTGAGGCCTGTTGCACATAGGCTAAGGCTCTAGT
TTAAACATCGGGGAAGGTCGGGAATGCCTGACACAAGGCGCACTAGGAGTTTCG
CTTTTAAGGGTGGAATTAGCACCCCAAGGCAATGAAGAATACACCGCAGGCACT
GGCCTACAATAGTCTTGAAGATAGAGC
>index395_0
TTTCTGTTGGTGCTGATATTGCTGTTTGCTTCCCACGTTCAAAGATCAAATTTG
CTGTCTGCTTCTGAAGGTCTAGACATGCCCTTTTACACAGGACAAGGCTTACCT
TCAGAAGCAGTACCATCAAGGTGTACTTGAGGCCAAGGCCTCGGAGGAACTTTA
CAGACTCTATCTGGCCTACACGCGGAAGGCGCCATGCGCGTGTACAATAAGTAC
GGGCTGTCCGCAAGAAGAAGATAGAGC
>index396_0
TTTCTGTTGGTGCTGATATTGCTGTTTGCTTCCCACGTTCAAAGAAAGGAACAT
ACCTCTGCAGTACCCTTGCAGTTCCTTTGTCCAGCAAGTGTCCAAGGCAACAGT
TCGGTCTGAGTTTTCCCAGGGGTCCTCCCCATTGAAGGCAGCGCTTACTTTCCA
AGAGCCGGCGAAGTCCCACTGATCAAAGGCTATGTTGGGAAGGTGAACTCTCGA
TCTGGGTCTAATGTGTGAAGATAGAGC
>index397_0
TTTCTGTTGGTGCTGATATTGCTAAACTTCCCGCTTTCCAACCACTTGCAAAGA
TCATCATGGAGGTATCACGACGAATGCTATGGACCCTCAAATCAAGGCATTCAG
ACGAGCCTCAATAAGGGAATTCACGCTAAAGTCGAAGGCAGTGGGATCACCAAC
GGTCCACATCTTGGCTTAGGAGGGAAAGGCCAATGAGCAGTGATGAAAGGGAAA
TCGTAACTAAGAGTCTGAAGATAGAGC
>index398_0
TTTCTGTTGGTGCTGATATTGCTAAACTTCCCGCTTTCCAAATGGAGGAGGTTT
ATCTCAGCTGTTGAAGCTTCACCGGCTCTGTATTCACGGGGATAAGGCTCTCCC
ATTTAGTACATGAATACGTCATTGGGGCGAGTTTAAGGCTCAATGACTGGACTG
TTTACAGCGTCGCTAGGTTGAACTTAAGGCGCACGGAGTAGTGGCCGAATAACC
GTGATTTAGTGTTGTAGAAGATAGAGC
>index399_0
TTTCTGTTGGTGCTGATATTGCTAAACTTCCCGCTTTCCACCCTCGTTTATTGT
CACTCACCTGAGTTAGGGATGGCACGTAACGCACACCAATGTCAAGGCGACGCA
GTTGGCTAGGTGACTGATAGGCCACTGCATGGACAAGGCTCGTACACTACTGAT
AATCTCTCTTCACACAGAGACGGGCAAGGCTATCGCCTTATTCCTAGGACGACG
TTACTCCCCTCGGCCAGAAGATAGAGC
>index400_0
TTTCTGTTGGTGCTGATATTGCTAAAGAAAGGAGGAAAGAAACCTTTCATGGCC
GTCTCAGCTACTGACATGAGGGAGTTCTCAGTCCCGCTCATGGAAGGCGTCATT
GATCCGTCCCACTCTTATTGTGACGTAAAACCCCAAGGCGTGCATCGTATTCGA
TGATCCAGTTAATATATTATCTTCTAAGGCCCAACCAAACCATTCCCTTGCAAT
ACACTCTTCTGCGGGCGAAGATAGAGC
>index401_0
TTTCTGTTGGTGCTGATATTGCTAAAGAAAGGAGGAAAGAAGGCGAAGAAAGAA
AGGTCTACCTCTGTGTCGCTGTGGTTGTACATCATGTCGTGATAAGGCTCCGCA
AAATGAAACGATGTACGCGGAGACGAAGAGGTTAAAGGCGAATTCGAACTGTAC
ATGAGTCCGCCTCCATACAAAACACAAGGCGCTGTAATGGGGCCTCTCGAATGG
AGACGAGGGTACTAGCGAAGATAGAGC
```

>index402_0
TTTCTGTTGGTGCTGATATTGCTAAAGAAAGGAGGAAAGAAAGTAGTCACGGAT
GACTCAGCTCACTCGACAGACTACAACCCGATTCAAGTGATCTAAGGCGGCATG
GTATTATAGTCTCATGTCGGAGAGTACGGGTCCCAAGGCTTGCACCGCGGTGAC
CACTTTCCTACATATTAGTGAGGGAAAGGCATGAATCGGATACATATTGAATTC
TTCAAACAAGCGGGGCGAAGATAGAGC
>index403_0
TTTCTGTTGGTGCTGATATTGCTAAAGAAAGGAGGAAAGAAAGGGCGTCACCTC
ACCTCAGCTAGCACGATCTCGTGGTGTACCTACTCGTGGACTCAAGGCGAATGT
ATGCCCTATTCTATTCTCGCTGGCATCTCTGTTTAAGGCCAAGCTTGAGGAGGA
CCAATAGAGAGCCGGGTATCGCGGCAAGGCGGGCAAGCCTGTGGGAATTGCTGG
AGTGTGTCAATGGCTCGAAGATAGAGC
>index404_0
TTTCTGTTGGTGCTGATATTGCTAAAGAAAGGAGGAAAGATGTCGACGGCGGAC
TTTTCACCAGAAGCGAGACTAATTCGTGTGCCCTCTACAGGGAAAGGCGGTGCA
GATATTAGCTGTCGGCCCTCCAAGAGTCCCTCATAAGGCGATCTGCCTTGAACG
GAACCACGACATGTCCCCTTTGAGCAAGGCTTTACGTTCGTCCCGAGGTTGAAG
CTGCTCGAGCGTCTGCGAAGATAGAGC
>index405_0
TTTCTGTTGGTGCTGATATTGCTAAAGAAAGGATCGTCATGTTTGCTTCCCACG
TTCTCACAGCAGACTGGACATCTTCGGGTTAAGTGGGGTGACTAAGGCATTATC
TCTCATCGGCGGCACAATTGAACCTACTAAACCGAAGGCCTCACCCCTGAATGT
TAAAACATTGCCTATATATCACTCGAAGGCCATGGTAACAGAGATACAGATCCG
TCTTAGGTCATCAGCTGAAGATAGAGC
>index406_0
TTTCTGTTGGTGCTGATATTGCTAAAGAAGTCAAACAGTTGTTTGCTTAAGTGC
CGCTCACAGCATGGCTAGTTACTGAGGTACTAGCCGGAAATTCAAGGCATTCAC
ACACTCCATCGTGGTAGTTCCTATTTGTGCCTCGAAGGCGAAAGTCCTGGCTAT
CGTAGAGTAGTCACAACGCAGCCGTAAGGCTTTGTAGAGCACAGGGGCCATTCC
AGATTGATGGACACTTGAAGATAGAGC
>index407_0
TTTCTGTTGGTGCTGATATTGCTAAGTCGGACATCGTCATCCCTCGTTTAAACT
TTATCTGCATCTCCATCGCAAATACTCCAAGTGGCCAAAGACAAAGGCAACGAC
TGTAGCTAGCGCATCCATATCCCGCACGTCACTTAAGGCAACTTGCTGGAAACA
ATTGCAGGCATATTGCTTATGCCGCAAGGCTTCCATCCTAAGTACACATTCCGC
GTCTCGAGTATTGCTGGAAGATAGAGC
>index408_0
TTTCTGTTGGTGCTGATATTGCTAAGTCGGACGGACATTTACGCACGTAATTCA
ATGTCAGCATGTGCAAGTAGGAAGCAGTACGGTAAAGCCAACTAAGGCTTACGG
CCCTTGATAATTGTTCACGACGGTATGACAGGGGAAGGCAAGATTTTGGTTCCT
CGTAAGCGCGGTCGGATTAGACAGCAAGGCCCCTGGCCTCATTACCCAAAATGG
GGATTCTCCGGCCCCGGAAGATAGAGC
>index409_0
TTTCTGTTGGTGCTGATATTGCTATCGGTTGTATTCCGGGACCCTCCCACCCTA
AAGTCCCTTACTCATCCTACACGGTACTGTCTAAAGCATCGACAAGGCGAATTG
GTAGATCACACCACTAGATCGTCTCGGAATGAACAAGGCGTCGATCATCGTTCG
AGGCTCGTGGCACCCGATTGCTGTAAAGGCAAGCATAGATTAGTTCCGTGGCAA
ATTCGCTAATTAGGTTGAAGATAGAGC
>index410_0
TTTCTGTTGGTGCTGATATTGCTCCCACCTCCCGGACTTTAAAGTTTCATTATG
TTTTCAACCAGTGACTGCTCGTAAGAAGTCCAATTGACCTACCAAGGCCCGTTT
CGGCGTGCCAGACTTTCTGTCTAGGTTGCGATGGAAGGCATGCCATAAAGATAT
CTTAAGCTGAGGGTACGAAAAGGACAAGGCGTCGTACTCTCCCGGTATGTATTC
TTAAGGAAGGATTTTCGAAGATAGAGC
>index411_0
TTTCTGTTGGTGCTGATATTGCTCCCACCTCCCGGACTTTACCACCCATTCGAT
GTCTCAGCTCAACCTTTCTCGGGGACTGTCAAGCAACCAATGCAAGGCCTGGGA
CAGATTTCAAAAGGATGACCTTACCTTCCCCACCAAGGCATTGACAACGTCTTT
GGGCTCGGACCTAACTAGACCTGAGAAGGCATGAATTTCAACATGCTCCGCTTC
GCTAGCGTTCGTACCGGAAGATAGAGC
>index412_0
TTTCTGTTGGTGCTGATATTGCTCCCACCTCCCGGACTTTAGGCGAAGAAAGAA
AGGTCCACATCCTTGATCGGGTTTTGCCCAGAACGGGTCATTAAAGGCCCAATC
CTGTTATTGGGACGGCCTCACTACCAAGTACGGTAAGGCTTATGATACGTCGTG
GAAATGATCCATCCACTAAGGCGCAAAGGCCCTTGCAGAGACTCAGGTTGGAAA
CTGAGTCGTGAGGCCGGAAGATAGAGC
>index413_0
TTTCTGTTGGTGCTGATATTGCTCCCACCTCCCGGACTTTAATCAATCAGCTTT
CCATCAACGTCTCGAAAGGTCTAAGACCGTTAGTTCAGGTTGAAAGGCTCTGCC
GTTCTGGCTAAGTCTGCCTGTATAAAGGGGAGGCAAGGCTTTAGCTGCGCTGTC
CAGCACACATTCATTATACTCTGAGAAGGCGTGTGGAGTTGGTGATTAGTTCTG
ACCCCGGCTTTAATACGAAGATAGAGC

>index414_0
TTTCTGTTGGTGCTGATATTGCTTCCGCAAAGATGGGTTTGGCGGAGTAATGTC
GTTTCAACCCAAGTGGGAACTTATGACGGCCAAATTCTAGAGCAAGGCGTCTGA
GATAGGTGTACTTCTCAAGTACAGCGCATTAGCGAAGGCCCTCACGAATCAGGA
AGTGCTGACAGACGCGCCACTGCAGAAGGCCGCAGCAGGCCAGTACTATGTGTC
GATCGTGCCAGTCTCAGAAGATAGAGC
>index415_0
TTTCTGTTGGTGCTGATATTGCTAAAGAAACTCCGCCTTTCCCACCTCCCGGAC
TTTTCAGCTAGCTCACATAGTGTTGGCATCCCCGCACTTAGAGAAGGCTGTCTG
ACAATATAAGAGAGTGCGTAGACCGATGCCTGCAAAGGCTTGCAGACCCGGTGC
AGCGCCGCATGCATAATAGTCATACAAGGCTTCGCTTGCAACCCGACTTTGGTC
GCATATGCACATGCCGGAAGATAGAGC
>index416_0
TTTCTGTTGGTGCTGATATTGCTAATCGACGTCCCAAACCAAAGTTGCACCAAA
TAATCTACCAGTCAGTTGAGCGTCAATTCCAACCCATGTACACAAGGCTGGATT
ACCGGTCGTGCCCTGTCGCTTCACTTTGAAGGTCAAGGCTTCCTCTTCTCTTGT
GACAGATGCACGGCGCCTCCCATACAAGGCGTTTTCCACGCTCAGGTTAGACTT
GGACTCCTAATCTCGCGAAGATAGAGC
>index417_0
TTTCTGTTGGTGCTGATATTGCTAATCGACGTCCCAAACCACCTCACCATTGCC
GTCTCATGCGAACATCTCAGACTACCGACCGTAAGTTAACTGGAAGGCCTCTTC
GTGCGATGTAGATGCGTCAAAATGTCAACATCCTAAGGCTGTAACGGAGCTGCG
TCCTACCTTTGGGGATACCTCGGAGAAGGCGCGTGTCACGGTGCAAAAGATCCT
TCTCGGACAGCCGCCTGAAGATAGAGC
>index418_0
TTTCTGTTGGTGCTGATATTGCTAATCTACTAACAACGTTAGGGAGGAACGTTT
CAGTCTACCTCTGTAGGGAATGTACCCCTTCATCTAGTCACAAAAGGCGTCATG
TGGGACTACTCGTTGATTTCGATCGGACTATTCGAAGGCGGCTGTCACCCATTA
CCGTTCACGTTAGACTGGCAGGGTTAAGGCTGCCCACGTCATCAGCAAACCCAT
TCTGATCCTATCTTTGGAAGATAGAGC
>index419_0
TTTCTGTTGGTGCTGATATTGCTAATCTACTAATGTCACCAACTCACCCAAACC
ACCTCTACCCAAGTGGAAGTGGTGGCACACTAAGTGCTGGCACAAGGCTCCAGT
CTTCGTGAGCTTTCTGAGGTATCCCTTTAGCCTGAAGGCATGTACCGTCGGTCT
CAGGGGTATGCAGTCGGTGCTGTCAAAGGCTGCCGACTACTATCTCACAATCCA
TAAAGGAAATTGGCCCGAAGATAGAGC
>index420_0
TTTCTGTTGGTGCTGATATTGCTAATCTACTACAAACAACAGGCGAAGAAACCA
GACTCAGGACAGACCTAGGAGACTGATTATGTCCTTACCCACGAAGGCAACCTA
GGAAATGAGACTGACGTCAGGGAATATCTTGAGTAAGGCTAACGAGTTGAGGAA
CGAACCGGAGCATGGGCTTACGCAAAAGGCAAGAAAGCCACCAGACAGTGGCCC
CGGTTCGTAGCCTAATGAAGATAGAGC
>index421_0
TTTCTGTTGGTGCTGATATTGCTAATCTACTACAAACAACAATTCCGTTCACCC
ACGTCTACCAGAGCTAGATGACTCGGTCATTGTGGGGAATCTCAAGGCGTAGCG
CAGCGTAAGGCATGGAGCACCACTACAATTGGAAAAGGCGCCTTCCGGCATACA
AGTTGTTCGAGTCGAGTATGAGGAAAAGGCCTGCGAATATTTGACTCGCGCATA
AGACGCCCAACCACCTGAAGATAGAGC
>index422_0
TTTCTGTTGGTGCTGATATTGCTACTCAAAGAAACGTTTAAATCGACGTCCCAA
ACCTCAGCTGATGCTCGAGAATCTCCGACGAACTTGGATATGTAAGGCGGGGCA
GTAGGTGAAGGCATTGCGACCAAGTGCTATTCAAAAGGCTGGTTTCTGTAGGCA
GGGGTGCTAATCCCCGGACGCTAGAAAGGCAAATAATGACGCTCGATTAAAGGA
CCCGGAACCGGCCCCTGAAGATAGAGC
>index423_0
TTTCTGTTGGTGCTGATATTGCTACTCCGTGAAAGAAAGGACAGACTCAAATCT
TCATCAGGACAGACTCTTGGCATCTCGCAGAATACCGCACCAAAAGGCTGCGCA
TCGATACGGATGTTGAGCATACAGGGTAGAATCAAAGGCCTGGAAACGGCTTCC
TTTTGCCTCAGAGGCGTTACCTTGTAAGGCCAGGTTAGGCTCATGAGCGCCATA
AACACAGGTAGGCCATGAAGATAGAGC
>index424_0
TTTCTGTTGGTGCTGATATTGCTCCCTAAAGCCCTCGTTTAAGGACGTAAACTC
AACTCCCTATCGTCCACATCATCATCCAGTCCATTATGCTAGTAAGGCGCTCGA
ATTCACGCTGCTGACTGACCAATGTATGATTCGAAAGGCCCCGGTAGGCGTCGT
CAATTGCCGTGAGACGTGTAACGGAAAGGCTATGACCTGGAAAATCACGAGTAA
CCGGGCCCGAGCGATCGAAGATAGAGC
>index425_0
TTTCTGTTGGTGCTGATATTGCTTCCCACTTACAATGATCACGGGTTTCCGGAC
TTTTCTGCAAGGTATGGAACCCCATTGGTCGAAATAACTGCAAAAGGCTGGAGA
CTTGTAGCGCGGTGGTGGCTCGCGCGTTATTGATAAGGCGACAGCAAATACACG
TTTCGCTCGTTGACTTTTAACGTGAAAGGCTGGCGTGTTGGTCCAACCGCTGCA
ACTCTCTCTGCATTGTGAAGATAGAGC

>index426_0
TTTCTGTTGGTGCTGATATTGCTTCCCACTTATTTCAAAGTGGCCGTCTTCGAT
GTCTCTACCACGTTCCTCTTTCGACGTGTATGGTCGTTAGGAGAAGGCTGATAT
CTCCGCTTTCAGCCGGCGTTTTGTACCAGTCTCCAAGGCGATGTGAGCCTCCCT
GGTTCTGGGTGGAAGGACTTTCCATAAGGCAATAGAACTAAGTGGCGGTTGATG
GCACCCATCTAAAGTAGAAGATAGAGC
>index427_0
TTTCTGTTGGTGCTGATATTGCTTCCGGAGGAAACTCAACAAAGAAACAACGGA
TGATCACAGGAGAAAGGTAGGGAGCTATCGTTTTACGATCCGCAAGGCCAAGGA
CGTCACTGTAGTTACGCGGCCCATTAGCCATACTAAGGCTTTCAAACACACCAA
AACACCCGAAATTTGTACCTTCCCGAAGGCTAGACCAAGCTTATGGCGATTATA
ATCGGCATCGTGTCGAGAAGATAGAGC
>index428_0
TTTCTGTTGGTGCTGATATTGCTTCCGGAGGAAACTCAACATTTCAAAGCACCC
AAATCACCTGAGACCAATAGTGGCTGGTCTTCAACTTAGTACTAAGGCTCTCAG
CGGAATCCACAACCTGGTCTCATCACAATTGATAAAGGCACGCCTAAACGAGCT
ACAATGCCAATTCGTGGGTGTCCTTAAGGCAGGTACGCTTATGCACGTCTAAAA
CCTTATCCGAAATGAGGAAGATAGAGC
>index429_0
TTTCTGTTGGTGCTGATATTGCTAAAGGATTTACCGAGTTACTCCGTGAAAGAA
AGGTCATGGTCAGGTTCCTCGTAGGGTCTAAAGATAGGTTGGCAAGGCTACCGA
GGCTGCATACTGTGTACCAGCGTGGAAGACTATGAAGGCATAGGCGTACGTACC
CACTGGTTGAGGCCGTCAGCGTCTCAAGGCCAGATGTACTGAATGGTGAATGAA
GAGCTTCGTCTTCCCCGAAGATAGAGC
>index430_0
TTTCTGTTGGTGCTGATATTGCTACAAACAACAACATGTTAAGACGCCTCCTAC
AACTCAGCTGTAGCATACAGCCCGATTTTACTGTCTTGGAGGAAAGGCTGTTCG
CGCGAAACGATGAGGCAGTACGTCGCCCAAACAGAAGGCGCTTATACCGACTAG
GAGACGCTGTATGTAGCGCATCAAAAAGGCCTACCATCGTCCTGTACCCAGATG
ACATTACCTATACCGTGAAGATAGAGC
>index431_0
TTTCTGTTGGTGCTGATATTGCTACTCCATCGACCGCTTTACTCAAAGAAACGT
TTATCAGCAGTAGTTCGCTTGCCCAACTATTAGAACCCCAAGGAAGGCGCCGGA
GTCACGTAACATCTAATCCTTGCATTACGCACGGAAGGCTATACGGTGCATTGT
CGATAAGGACGTGGTCGCGGCTAAGAAGGCGGATCAGCACGAGTGGATATGTGG
CATTTTCCTAAAGTTAGAAGATAGAGC
>index432_0
TTTCTGTTGGTGCTGATATTGCTATCGGTTGTTCCGCCTCACCATCTTTATTGT
CACTCACACGTCACGTACACCATTACGGCCAACGTACTTCCTCAAGGCCAAGCT
TAAAGGGAGACAAACATTTCAGGATAGCCGCAGTAAGGCGGCCTACGGTGCCTT
TGCACACCCCGTAGTCGAATCCGTGAAGGCACAGGAACCAGACCGCAACATCCC
AGTCCAGCAGTAATTAGAAGATAGAGC
>index433_0
TTTCTGTTGGTGCTGATATTGCTTCCGGAGGACCACTTGCAGGCGAAGAAAGAA
AGGTCCCAACAGTACATGGCGAAGCTTGCCAGTCTGTCAGTGCAAGGCGACTAG
AGCACGTTATTTATCCAATGAGGGGCCCCGTTTGAAGGCCATAGATCAGACTAA
CGGCTTCATATGTGAAGATAGGGCTAAGGCTTAACGCTTGACCGAGCAGATAGG
ATGGTGGATATTCGGGGAAGATAGAGC
>index434_0
TTTCTGTTGGTGCTGATATTGCTTCCGGAGGACCACTTGCATTATGTTTAATCG
ACGTCACGTACGTCCATTAGAGATCACACACCGACAGAGAACTAAGGCGTGAAT
TGCGCCACAACTGTCAGTTTCATTCCGATGGGTGAAGGCTTGGGTAAATGAGCA
CTCGGTCCACCGTTGGAAGGGTTAGAAGGCCGCATGCCGATGCAGCTCAACTTC
ACCACAAGCCTTCCTGGAAGATAGAGC
>index435_0
TTTCTGTTGGTGCTGATATTGCTTCCGGAGGACCACTTGCAAAGAAAGGATCGT
CATTCATGGTCCACCACTTAGTAAGTCGACATGATGCTCGTATAAGGCTCTGCC
GACTGAGTCTGATAGTCGAAAATCTGACAAGCCGAAGGCTGGCAAATGCGATGC
TCGAACAGACGGCTCAAATTAGGAAAAGGCTGATAATTGTCAGCAGCACCACAG
CTTCGACGGTGTTTAGGAAGATAGAGC
>index436_0
TTTCTGTTGGTGCTGATATTGCTTCCGGAGGATCGGACTTACCACCCATCCTAC
AACTCAGCAGTCTCCATGTTCGAAAGAACCCTCTACGACTTCAAAGGCGGGGCT
TGTCACCCGACCTATGACTTAAGTGGCCGAGGAGAAGGCGGTCGTATTAGTGTA
AGACACCGATACTCCGGTACCCTACAAGGCGCTTTTAGTGTTCTGCAAAGTAGA
CCCTATTATTACCAGCGAAGATAGAGC
>index437_0
TTTCTGTTGGTGCTGATATTGCTTCCGGAGGATCGGACTTACGGGTTTCCGGAC
TTTTCAACGGATGAAGTCAGTCCTATGAGCAACGTCCCAAATTAAGGCCTACAC
AGCGCCGCGCTCAGTACCCACTCAGACCTATGAAAAGGCTACTTACCCTAATAG
ATCCGTTGACCTCTAGGGTCCGCTCAAGGCTGTGTCAGGGTGAACCGTGCGCCA
TATGTTAGACCACGAGGAAGATAGAGC

```
>index438_0
TTTCTGTTGGTGCTGATATTGCTAAATCCGTTACGCACGTACCGACGAACAAAC
AACTCTACCAGGAGCACTAGGTCATGCCTCATCATGCCTGTTAAAGGCCACAGT
TCGGTTGGGCCGATTGAAAAGCACCAGTCGTTGCAAGGCCTGGTGAGATTGAGT
TGAACCTGACGCCTTATGAAGAAACAAGGCTTTATTATCATATCTGATATCTGG
TCAGGAGGAGTCGGGCGAAGATAGAGC
>index439_0
TTTCTGTTGGTGCTGATATTGCTAACAGGACCAGGCGAAGAACAGGACCAGGCG
AAGTCCCTAAGACGACCTCTATAAGAGTGCGCAACAAGCGAGGGCGCTTCTACT
TGATCGAACGGAAGCCAGTTCAGGGCGGCCATCCGCGCTTAACTCCCAGTGGAT
CAATACGACTAGCGGCATTATCGTAGCGCTAGAAGACTGACGGGTAGGACTTTT
GCTCAGGGAAGTCGCGGAAGATAGAGC
>index440_0
TTTCTGTTGGTGCTGATATTGCTAACAGGACCAGGCGAAGAAAGAAACAACGGA
TGATCAGCAGTCATAGACTGTCCTTCAGAACTGGATTCTGAGGGCGCTAACCGC
CTCGTCTCTGGATCAACTGCAACCAAAACCCCACGCGCTACCCAAGAGTCTTAG
TGGGTTTTGTCACGGCCCGTTCTCTGCGCTTATTTACTCTATGATTGCTCCGCT
TTCCCATGCCCGGGTTGAAGATAGAGC
>index441_0
TTTCTGTTGGTGCTGATATTGCTAACAGGACCAGGCGAAGAAAGTTTCCACCCA
CTCTCGCTTAGGTGATAGTCCTGGGGTAGACCTGAAAAGAACGGCGCTGGGCCA
TTGCCTCGTTCTGCTGTCGTAGGTAAGCTTGAGTGCGCTTTACAGTGCCACAAT
CTTAATGCGCATTGAATGTCGCGCGGCGCTCTGGCGCGGCACTCTTACGAGTTT
TAGCGGTCCGCAACACGAAGATAGAGC
>index442_0
TTTCTGTTGGTGCTGATATTGCTAACAGGACCAGGCGAAGAATGGAGGAAAGAA
GTCTCAGCTACGAGGTTGTAGGCTAATACGTTGCAGGAATGCGGCGCTAGGAGT
CAAACATGCAATGAAAAGTGACCGCTGTTTACCAGCGCTCAGCCAGTATGTACT
TACGAAACTGGGAAGGTTTCTAAGTGCGCTGCTCAACTTCTAGAGTGACATGGC
GCCTACCACTATCAGGGAAGATAGAGC
>index443_0
TTTCTGTTGGTGCTGATATTGCTAACAGGACCAGGCGAAGAAAGAAAGGAACAT
ACCTCATGCCACAGCTACTGCATCAGCAACTCAATACGGTTACGCGCTAAAGGA
TGGACCGTCCACTCTATGTTTCAACTGGGTCCTCGCGCTGTCCCCAAGCCGTGC
GCTTAAAGGTATTCGAGATCTTAAGGCGCTGTTGAGCACTAGTGGCGTCTAAAC
CGTACGTAAGGGCTCCGAAGATAGAGC
```

```
>index444_0
TTTCTGTTGGTGCTGATATTGCTCCGGCCTGAACTAGTTCCCGGCCTGAACTAG
TTCTCATGGGTTGCAGACATAGATGACTCCACGTAGAGACGAAGCGCTAGTGAA
AGCTGTTCACAACCAGGCAAGTGCTTCGTGTTCTGCGCTGCGATGGTCGCCAGA
TAGAGGAGTCGAGAATGACTCCCTGGCGCTAGCATACTTGACTTTACTTTAAGA
ATGGACAGCGGGTGTAGAAGATAGAGC
>index445_0
TTTCTGTTGGTGCTGATATTGCTCCGGCCTGAACTAGTTCAGTCGCTCAGGGAA
GATTCTGCAGTACATCGCCTCTTGGTCGAATCGCCTAATACCTGCGCTGCATCA
TCAAAATACGCGTCATTTAGCCCGGTAAATTTGCGCGCTGCTCGCACGTCATCC
GGAAAAGGTGCCGATTTGCCATCTCGCGCTGTGAAGTCTACAGAGCCTTTTGGC
TAAGAAAGAGTGGGCAGAAGATAGAGC
>index446_0
TTTCTGTTGGTGCTGATATTGCTCCGGCCTGAACTAGTTCAGGGAGGAAACAGA
TCATCACGATCCTGACGACATTTCACAACCCGAATAACTCTGCGCGCTTGCCGC
GATTTCGAGGTATAATCCACGATTCATTACGGAGGCGCTCTCTGTCGTCGCTTG
ACACACGGCATAGTAACCAGCTTGTGCGCTCACTGACGCGATTTGTTAATGGGC
GTGAATAGCGTATGGCGAAGATAGAGC
>index447_0
TTTCTGTTGGTGCTGATATTGCTAAACCACCAAGTCCCACTCAATAACCACCCA
CTCTCTGCACTGTGCTTAGCTGAAGCTCAGTCGGACCTTGAGGGCGCTCACTGT
TCCTACATTTCGGGGTAAGGTGATTTTAGCGACCGCGCTGTACACGGCGGTTGA
GACCGCGCAAGGTGTGAATAAACCAGCGCTATGAGAAGCCTAACAGCTGGATTT
TGTACTTGTCCAGTTTGAAGATAGAGC
>index448_0
TTTCTGTTGGTGCTGATATTGCTAAACCACCAAGTCCCACTCCTACAACAGGGA
GGATCACCAGAAGCGAGTTGAGCTCGTTTCATACCCAGTAACAGCGCTAAAGGA
GATGTCCTCGTCCGGCTGTGAGTGGACACCAATAGCGCTAGTTATGGCATATCG
CATCATCCTTGATTTTCTCGGGCGAGCGCTTGCGCTCTGCTCGTACCACGTCAC
AGCTCGACCAACGGAAGAAGATAGAGC
>index449_0
TTTCTGTTGGTGCTGATATTGCTAAACCACCAAGTCCCACGTTTGCTTCCCACG
TTCTCTGCTCAACGCTAAACCTATGTCACGAGAACCGCCAACAGCGCTGACTAT
GAATGATAGGTTCCAGGTTCGACGTCCCAGGCTGGCGCTTACCCATCTTGAATC
TTGTCCCCGAGGACCAGGCGGTACAGCGCTTACCCCGCGCTATAACACTGGACG
GTTCACCGACAATTAAGAAGATAGAGC
```

>index450_0
TTTCTGTTGGTGCTGATATTGCTAAAGACAGACCCTCCCAAACATGTTACCATC
TTTTCTGCAGTCTTTGGGGAGTCTGAAGGAAGTACTCAAAAGAGCGCTCTTCCG
TGGTGTTCTGCATCAGGTGGAGATCAGGGGTGTGGCGCTTTCACCGTATTCGAA
ATGGCGTAGTAGAAGACCTTATAAAGCGCTACCCAAGTGCTCAGCTGTGCCGCT
TGGGTAGACACTATGGGAAGATAGAGC
>index451_0
TTTCTGTTGGTGCTGATATTGCTAAAGACAGACCCTCCCAAGGCGAAGAAACCA
GACTCAGCATCCTTTCGTAGAGTTGGGCGGTGTGGAGTGACACGCGCTCTTTCA
ACATGGTATAATCTTAACAGGGTCAGTTAGTCGCGCGCTTTATGCTACACTGGG
AGCATAGACATGAAGGGATGTTCACGCGCTGTCGTTACACGCTCATTGAACCGC
GAATAATATCGCCCTCGAAGATAGAGC
>index452_0
TTTCTGTTGGTGCTGATATTGCTAAAGACAGACCCTCCCAACGGGTTTCCGGAC
TTTTCAGCAGTCAGATACCTTCGCTACGTCAAAATGTGAGCATGCGCTCGCGAT
CTTCAGAGAATAGGAAAGGCTGTCATCCGCAGCCGCGCTAGGGTTGTGATCTAC
AGCAGGCTGTGAGCCTTGGTTCAGAGCGCTCTTTATTTACCCAGATTCCTTGGT
GACTGACGTTTCCATAGAAGATAGAGC
>index453_0
TTTCTGTTGGTGCTGATATTGCTAAAGTTTCATTATGTTTCCGGCCTGAACTCA
CCCTCACCTTGCATGGTCAAACGCTCCGATATCGGAGTTACTGGCGCTGTTGCC
GTAGGCCGAACTGCGAGAGTCAGTGTGAGTGCTCGCGCTGATATTAGAGCGAAA
TCCACCCATAGAAATATGGGCTATCGCGCTACGAACGTTCACGAGAGTTGTGTG
TGCACCCACTGGAATGGAAGATAGAGC
>index454_0
TTTCTGTTGGTGCTGATATTGCTAAATTTCATCCGCAAAGAAACCACCAAGTCC
CACTCCCTATCCTTTGGCCTAGCTGTCCAGTCTCTCATCCACAGCGCTGTCCGT
TGTGTGCACGTTCCTTACTCTGCCACATTTTACTGCGCTGTCTCTCTTCCGTTG
GCACGCCTTGTATCGTCACTCCTAGGCGCTCTGTGAGAGAGCCCCGTAGCAGCT
GGATGTAACTCGGGCGGAAGATAGAGC
>index455_0
TTTCTGTTGGTGCTGATATTGCTAAATTTCATCCGCAAAGAAAGGGATGGGTAG
GTTTCGCTACTACGTCGTCATTTGACTGTTGTCGAAACAGTGTGCGCTGAAGAG
GTGATCTCATGACACATGTGCTCTGGCGGCTTTGGCGCTTCTCCTTACTCATCG
ATTGCGGGACACGGAGCTCAAGCGAGCGCTTGAGCCTAGGGGAAGGAGACGCTT
CGGACTCCGAGCACTGGAAGATAGAGC
>index456_0
TTTCTGTTGGTGCTGATATTGCTAACATGTTACCATCTTTAAACCACCAAGTCC
CACTCACGTAGGTAGCGTTGATAGCTCTATCACCAACCCAAGGGCGCTTAGAAT
TCTCGACAATGGATTCGAACCATGTGAGTGTAGTGCGCTCTTCAGAGAGCGCCA
AGAGGAGAGACCCATGGCAAGCTATGCGCTCTCCTAACAGCGTAACCCCGGCAC
CAACCACATCTAATGTGAAGATAGAGC
>index457_0
TTTCTGTTGGTGCTGATATTGCTAACATGTTACCATCTTTAGGTGCGTCAGGAA
AGATCAGCTGTTGAGGTGACGATAACCTGAGTCTAATGCAACGGCGCTAGATAC
ACGGCGCTCAACCAGTGTTTGCTTGGATCAAGAGGCGCTAGTGCGTTCAATGGG
TGCGACAGTCCCGCATGGCAGCCGGGCGCTGTGTATATTTTCAACAGAGATAAT
GATGTGGCCATCGCGTGAAGATAGAGC
>index458_0
TTTCTGTTGGTGCTGATATTGCTAACATGTTACCATCTTTCAGGCGTTTCCGCA
AAGTCAGGACATGGATAGACACTCTAGCCCGTGTTTACCACTCGCGCTCAGTGC
CGTGGCAATAGGTCCTGTTCTGTGGCAGGAAGTGGCGCTCATGGTCAAAAGTGG
TGGAGTCCCCGTCAAGGTCGGTAGTGCGCTCGATGGGTGTCTCTGAGTGTGACG
AAGCCACTCGAGAAGGGAAGATAGAGC
>index459_0
TTTCTGTTGGTGCTGATATTGCTAACATGTTACCATCTTTCCCACGTTCATCGT
CATTCACAGCTGAGCTATGGCTGAGTCGTGTAAGGCCTGAGAGGCGCTGTATCG
TTTGTCTTTTAAATAGACCCCAGGCAGCTGACGGGCGCTGTTCCGGGACAAATC
GATCCCCGATAAATGGGGAAAACCAGCGCTTGGTACGCCCCGCTGCTTGATTTT
ACAGGGTGCAGCATATGAAGATAGAGC
>index460_0
TTTCTGTTGGTGCTGATATTGCTAACATGTTACCATCTTTAACTCACCCACCAG
ATCTCAGCTTCCATTCGTTCTGTACCAGGTTGGGTGCCACAGGGCGCTGTCAAA
ACCGGTTCCGTCTCGTTTCTACAGCGACCCAACAGCGCTCAAACACCAGCAGAC
CCCAGAGTACCTACGGTTCACTGTTGCGCTTGATTTGGCTAATCGGCCAAACAT
GCGACATGTGAAGAGGGAAGATAGAGC
>index461_0
TTTCTGTTGGTGCTGATATTGCTAACATGTTACCATCTTTAAGTCGGAAAGGGC
GTCTCATGCCATGCGATATCCTAGCTTGTTCGTGCATGGGTCGGCGCTCAGACG
CTGTTGATACATGACACTACCTTCTCCCTGAGCCGCGCTAGAGCAGTGCCCGCT
CTTGGACCTCATAGAAATCATGCGGGCGCTATCCGGTGAGTCGATAGGACGCCT
TAATTTATGTTAAGCGGAAGATAGAGC

>index462_0
TTTCTGTTGGTGCTGATATTGCTAACATGTTACCATCTTTAGGGAGGAAAGTCC
CACTCAACCACAGGGTCTTATCTTCACGCAGACCTGGACCGATGCGCTATAAGT
CACCCACAAGGCGTAGACCCCTGGAGTCTAATCTGCGCTAGGTCTTAGTGTAGC
ACGATCGACAAAGCACTTAGTTTCAGCGCTGCCCACTTTCCCTTGGCGCTGATC
CTGTTCTAGGCGATAGGAAGATAGAGC
>index463_0
TTTCTGTTGGTGCTGATATTGCTAACTCGTTAATCATGTCAGTCGCTCACTCCA
TCGTCAGCACAGTTGTGCAGTCCTTCACTAACCCCTTTTCGAGGCGCTAAGGTA
TGCTATAGGTCGCCTTTGCTTACCCCAACTAACTGCGCTCCGCCAAGACCTTAC
GCTCACGGGTATAGGCTGTTGGGGAGCGCTTGCTAAATTAGACATACCTCGTGT
GGCGTATCACTATCTTGAAGATAGAGC
>index464_0
TTTCTGTTGGTGCTGATATTGCTAAGTAGTCAATCTGTTTACCACCCATCCTAC
AACTCAACGCAGTTAGTGGCAGTACGCGTGACTAGGATCAGGCGCGCTACGTGA
TTGGGTCCTTGGATTGAATCATGAAAGCAAGATCGCGCTACTAGATAGACACTA
TGATAACGGTGAGTCCTATCTGGGTGCGCTGCGGCGCTCATCGAATCAATACCA
ACACGATTGGAACCTGGAAGATAGAGC
>index465_0
TTTCTGTTGGTGCTGATATTGCTAAGTAGTCAATCTGTTTACCGTTCTTAGGGA
TCCTCAACCCAACGGTCATACTTGAGGGAATCGCTGGCGTGAAGCGCTCCCTTT
CCAATTAGGCAGTTACCATATTCGTGTTATCCAGGCGCTAGAAGCCATTCTGCA
TCGAGGAATCGCATATCTCTGCTACGCGCTATAAGCTATACCGTGGCAAATTCA
ACGATACCGTTCGCGGGAAGATAGAGC
>index466_0
TTTCTGTTGGTGCTGATATTGCTAATACCAGACCGATCTATGGCCGTCTTCGAT
GTCTCAGTCAGGATGCGATAACCTTGCAAGTTCTCCGCAGCACGCGCTATAGTA
TTCGCCGGCACCATCGACGGGTTAGCTAATGTAGGCGCTTCTCTTTCATCCCAG
GAGGGTTACAGATGAAGTCTTCCACGCGCTCCGGTTCAGCGGAGAATTGTAGGC
AATCCGAGTTCCATAGGAAGATAGAGC
>index467_0
TTTCTGTTGGTGCTGATATTGCTAATACCAGACCGATCTAAGGCGAAGAAAGAA
AGGTCAACGCAGTAGATCAGTCCCGCAGAGAGTGTGTTGTACTGCGCTCAATGT
AAGAGCTGCGCACGGGGCAACCGAACGCTCTGTGGCGCTTAGTAAGCTTCACCT
AGACAGCCGTATCAGTAAAGCGCCAGCGCTGGACCTTAGCGGTGCGTTTTAACG
CGACGTCATATCGTTGGAAGATAGAGC
>index468_0
TTTCTGTTGGTGCTGATATTGCTAATACCAGACCGATCTAATCGGACTTACCTC
ACCTCAGCTTGGTGGTCTATAGCTTTCCGTGGTTGAGGCGGAAGCGCTTTAGGT
TTGTGTACGGTATCATTGCACGGGACGTCGGGAAGCGCTGACATCATTTCTGGC
CATGGGAGCTAGAAAGACCAGGAGCGCGCTCCTTAAACCTAACTACCCCTCTAC
ATATGTGTTCAGATCCGAAGATAGAGC
>index469_0
TTTCTGTTGGTGCTGATATTGCTAATCTTGTCCTCAGTACAAAGACAGACCCTC
CCATCATGCTGTCATGGGGTAGTCATTGGCCTCTGCATTAGTCGCGCTATCCCT
TCCGATCTCAATCTGATTTGCCTTCTTCACTGTTGCGCTCCCGGTGGTCGACGG
AAGGCCCCTGGGCACACCCTCTGTTGCGCTAGCCCTTGAACACGCTACGGGTGA
TCCTCTAAACCGCTGCGAAGATAGAGC
>index470_0
TTTCTGTTGGTGCTGATATTGCTAATCTTGTCCTCAGTACAACATGTTACCGGA
GGATCAGCTGTGTTTCACTGACTTCAGGGCGCTATTCAGCGTAGCGCTACCATC
CAGAAGGGTCTAGCCGAAATAAGCCGACAGCTGAGCGCTTCAAACCTCCTCCAT
GAGAGGTACTGCGCTTCTGCGACGTGCGCTGTTGAAAAGCCCAGCTAATGGAGG
TCATTAGTGGTGCTACGAAGATAGAGC
>index471_0
TTTCTGTTGGTGCTGATATTGCTAATTTAACAGGCGAAAGAAAGACAGACCCTC
CCATCTACCACAGAACCATGACATCACCTACACGAACTCAGGCGCGCTTGACTG
CATCGATCGAGGCGCCACGACAACATCGGTTACAGCGCTGCCGCAACATAACTC
GGCTTACGTCTTTTCGGCGCATACAGCGCTGGAGGATCCCTTAGATCTTCCTAA
TGGGGCGCATGGGGTGGAAGATAGAGC
>index472_0
TTTCTGTTGGTGCTGATATTGCTAATTTAACAGGCGAAAGCCGGCCTGAAAGAA
ACATCCACAAGACTTCCCTAGCCTCTAATCCTTTCAAGTCTGCGCGCTATACCT
GATCTTCCAACTCTACTCTGGGATGTCGACTCATGCGCTCGTCATGAAGCAAAC
AAGCCGAACGATTCCCCTGGTACCGGCGCTTCACCACCCTAAAAGAAGAAAGTG
CGCCTCAGCAGAACATGAAGATAGAGC
>index473_0
TTTCTGTTGGTGCTGATATTGCTAATTTAACAGGCGAAAGACCTCACCATCGGT
TGTTCATGGCTGTTAGCAAGCGAGTCGCACAGAGCTCATCCTAGCGCTGTCATC
TAGTAACCGGGACAGGCTTGCTAGCGCTGAGAACGCGCTATCCATGCATCAAGA
AATGGATTGAACCACTTATACGAACGCGCTACCCTACTCGTGGAATGAGGAGTC
AGGGTGTTCAGAACTTGAAGATAGAGC

```
>index474_0
TTTCTGTTGGTGCTGATATTGCTACCACCCATCCTACAACAATCTTGTCCTCAG
TACTCACCTTGACGAACCTATCTGGTGCACCCAGCAGTATTTGGCGCTAGTAAT
GCCATCGTCCGGAGACTCAGCTGAGAGAAGAGGCGCGCTCTAGTCCGGGTACTC
TTGTTATGAACAGTGAGTAGCCAAAGCGCTTCGGCAATGATACCACCAGTTGAA
AAGAAGTGTTACATACGAAGATAGAGC
>index475_0
TTTCTGTTGGTGCTGATATTGCTACCACCCATCCTACAACAGTTAAGGAAATCA
AACTCAACGTCTGGCAATGAAGAGCGGCTGTCTTAGCCGAAGCGCGCTGATAGG
GAAATCAGCGGCTGGAGTCAGATCTACCATTTTCGCGCTGAAGCTACGTGGGTT
TGAGAATATTGTCTCGTGAAAGTAGGCGCTCACGCTTCAGGTATGTTAAGTACG
GGGCCTTGAGGTGTGCGAAGATAGAGC
>index476_0
TTTCTGTTGGTGCTGATATTGCTACCACCCATCCTACAACAAAGAAAGGAGGAA
AGATCGCTTAGCTTTCTCGTTGATGCTTCTGTCTCGAGGGTTAGCGCTGGTACA
ATTCATCGGCGCCATCCTCCCCACGTCCATCGAAGCGCTCGTTTGACAACTTTC
TTCGCCCAGTACAGACTAAATGTCTGCGCTGCATCAGTATACTTCCCTTGAGTT
GTTTGTAAAGGAGACCGAAGATAGAGC
>index477_0
TTTCTGTTGGTGCTGATATTGCTACCGACGAAACTTTGTTACTCAATAACCCGT
AGATCAGCTCTTCTACGTTCTGATCGGTCGTGAGCTATACGATGCGCTATCATC
TGTCTGGCTTGTGTAGTGCATACGAGCGACGGGAGCGCTCAATTTAGGACTGAT
GACTGACACGAGACGTCCCTGAACAGCGCTGATTTCCTCAATGAAAGTGTCTGT
CAAGTTAGCCTTTCACGAAGATAGAGC
>index478_0
TTTCTGTTGGTGCTGATATTGCTACCGACGAAACTTTGTTAATTCCGTTACTCG
GACTCCCTAACGAGAGGTTTACCGCCTTTACCTATAGCGATAAGCGCTTGCGCG
TAGCAAATTACCACCAAGTCTCCTCCTTCTCAAGGCGCTTTAATGGCAACAGCA
TCAAACACTTGGTAGCCCAATCGAAGCGCTCAAACCTTACTACGGGGAGATTCT
GTCACGGGGCTCACCAGAAGATAGAGC
>index479_0
TTTCTGTTGGTGCTGATATTGCTACCGACGAAACTTTGTTAGGCGAAGAATCTT
TACTCGCTTAGCACTGGTAGCAGGTTTGACGCTCACCATTTACGCGCTTCGCCG
AGAGTCCCTTAAGAGTGTAGGCTGGTTGTGTAGAGCGCTCTCTTAAAATTGGAC
TTATGGACCTGAATGTAGAGTGAAAGCGCTGCTCCATGCGCCCGTTTGGGGCGA
ATTAATGCAACTGACCGAAGATAGAGC
```

```
>index480_0
TTTCTGTTGGTGCTGATATTGCTACCGACGAAACTTTGTTTCCCGTTTAAGGGC
GTCTCTGCTCAGAGGAGAATAGAGATGCAAGTCCGTCGAAAGAGCGCTATTCCG
AATGCCCCACCGCCAGTTGGGTGCCCTCTTCCAGGCGCTTAACGGACGTAATCG
CCCTCTCCAGAGTGTGGTGTACGAAGCGCTCGGAGTGTGTATACGCCCATGACA
ACAGAGGAACCTTCAAGAAGATAGAGC
>index481_0
TTTCTGTTGGTGCTGATATTGCTACCGACGAAACTTTGTTACCACGGAAATCAA
TCATCAGCTCTACGGTATACCTACTCAACTGTCTTGCGTTGAGGCGCTGAATGC
AGGGCAATCGCACTGACGCAGTATTGTGCTGGACGCGCTCTGAGGCAGAGACAC
CGCACTTATAGGGTAGATGGGTTTCGCGCTCACAACTCCACAATCAAATACTAA
AAGAGTTTGCTCTCTCGAAGATAGAGC
>index482_0
TTTCTGTTGGTGCTGATATTGCTACCTAATCAATCAATCATCCGCCTCAATTTG
CTGTCTGCAGTACGCTCAGACTGTAACCGGTACACACCCTAAGGCGCTAGTCGG
ATCCTGTAAATCTAATCGCACGCACATACACCCGGCGCTTATGGGACTTGGTCG
CGAGTCGCTCTCCCACATGGTGATGGCGCTTCGAGTATCCTTCTTAGTCTATGA
CGTACGAGACGGCGTTGAAGATAGAGC
>index483_0
TTTCTGTTGGTGCTGATATTGCTACTCAATAACCCGTAGAACCGACGAAACTTT
GTTTCAGCTTCGTTGCAACCATTCTACCCTTCTCAGAGCACGTGCGCTGCTCCA
AAGGTGGTTGGCACACCTAATCCAGTCGCTTAGGGCGCTATCTAGGGAGTAGGA
AGATGGGATTTCCGCCTAACAGGTTGCGCTTAGCTAATAACTAACGGCAGGGTC
TGGTTTTACAGATCAGGAAGATAGAGC
>index484_0
TTTCTGTTGGTGCTGATATTGCTACTCAATAACCCGTAGAAAGTAGTCACGGAT
GACTCGCATGTCATCGGATCGTTATTCGATCCGGTGGTAGGATGCGCTCGTACC
GCGTGTCGTTGCATCCTGAACGAGAATTTCATAGGCGCTTGATGTAACCGTATC
AAACACAATTCGGTGGGTACTCGACGCGCTTGCGTTAATGCCACCATATTCCTG
GAATTAAGTGCTTACTGAAGATAGAGC
>index485_0
TTTCTGTTGGTGCTGATATTGCTACTCAATAACCCGTAGAAACCCTAAAGGGAA
GATTCAACCCATGCCACATTAGGTTTGAGTGCATGCCTCCTGGGCGCTCGTAGT
AAGGCGGGTCTTCAACTTTGAAGACTGTGTGGTAGCGCTTTCCCTTATTGTCGA
CGGTAATGTGGCACCAACTGGTCGTGCGCTTGACGATGCGAGCTAATGCACGGA
CCTTACGTATTTTAAAGAAGATAGAGC
```

```
>index486_0
TTTCTGTTGGTGCTGATATTGCTAGGCGAAGAAACCAGACAATCTTGTTCAGCT
GGATCACGTAGGTCCAATGCACATGCAGCGAATGACTGCGGACGCGCTCATTCT
TATGCACGGTGGAAGCATTACTATGACCATCCCTGCGCTTAGTACAAATTGTGT
CATGGTGCACCTTCAGGAGGGCGCGGCGCTAACGTCAGGTTTGACCTGTTCTCC
GCCTGAGAGCGCAAGTGAAGATAGAGC
```
